# Reduced insulin signalling in neurons induces sex-specific health benefits

**DOI:** 10.1101/2022.09.19.508502

**Authors:** Maarouf Baghdadi, Tobias Nespital, Andrea Mesaros, Sandra Buschbaum, Dominic J. Withers, Sebastian Grönke, Linda Partridge

## Abstract

Reduced activity of the insulin/IGF signalling (IIS) network extends healthspan and lifespan in mammals and possibly humans. Loss of the Irs1 gene increases survival in mice and causes tissue-specific changes in gene expression. However, the tissues underlying IIS mediated longevity are currently unknown. Here we measured survival and healthspan in male and female animals lacking Irs1 activity specifically in the liver, muscle, fat and brain. Tissue-specific loss of IRS1 did not increase survival, suggesting that lack of Irs1 in more than one tissue is required for lifespan extension. Furthermore, loss of Irs1 in liver, muscle and fat did not improve health at old age. In contrast, loss of neuronal Irs1 increased energy expenditure, locomotion and insulin sensitivity, specifically in old males. Neuronal loss of IRS1 also caused male-specific mitochondrial dysfunction, activation of Atf4 and metabolic adaptations consistent with an activated integrated stress response at old age. Thus, we identified a male-specific brain signature of ageing in response to reduced IIS associated with improved health outcomes at old age.

## Introduction

Human lifespan has been increasing in many parts of the world for the past two centuries, but healthy lifespan has not kept pace(*1*). Ageing is characterised by a general decline in physiological function and an increased prevalence of multiple age-related diseases, including cancer, cardiovascular and neurodegenerative diseases. The prevalence of age-related disease shows a clear sex difference in humans, whereby women on average live longer but suffer greater age-associated morbidity(*2*). Ageing is a malleable process that can be ameliorated by genetic, dietary and pharmacological interventions, with the potential to delay or even compress age-associated morbidity(*3*). However, the response to longevity interventions is often sex specific(*2*), emphasizing the need to include both sexes in longevity studies.

Reduced activity of the insulin/insulin-like growth factor 1 (IGF1) signalling (IIS) pathway is associated with increased longevity and improved health at old age in animal models, including worms(*4*), flies(*5*), fish(6) and mice(*7–10*). Due to its high evolutionary conservation, IIS has also been suggested to be involved in human longevity. Indeed, recent genome wide association studies have implicated variants in IIS pathway loci with longevity(*11*), and studies of rare genetic variants have found enrichment of IIS variants in centenarians, suggesting a relationship between IIS and longevity in humans(*12*). The finding that a longevity-associated allele from humans reduces IIS activity in cell culture(13) further supports this link. Therefore, understanding how reduced IIS mediates longevity will help to decipher the underlying biological mechanisms of ageing and the development of therapeutics for healthy ageing in the future.

The IIS network plays a central role in regulating growth, metabolism and survival. In mammals, intracellular IIS activity is initiated by two receptor tyrosine kinases, the insulin receptor (IR) and IGF1 receptors, which upon ligand binding phosphorylate insulin receptor substrate (IRS) proteins, which are key downstream mediators of pathway activity. While mice have four IRS proteins (IRS1-4), IRS3 is not present in humans(*14*). Mice globally lacking Irs1 activity (Irs1KO) are long lived(*15*). In contrast, Irs2 knockout mice showed reduced survival, suggesting a specific function for IRS1 in longevity(*15*). Importantly, Irs1KO mice were not only long-lived but also showed resistance to a variety of age-related pathologies including adiposity, ulcerative dermatitis, reduced bone volume, motor dysfunction and age-related glucose intolerance(*15*). However, global reduction of IIS also has drawbacks such as reduced body size, reduced fertility, and metabolic syndrome(*16*). It is unclear in which tissues IRS1 acts to affect longevity and what are the underlying molecular mechanisms.

Transcriptomic analysis in livers of Irs1KO mice have linked altered IIS to mitochondrial function(*15*), and subsequent molecular studies have found that livers of IIS mutant mice exhibit reduced mitochondrial respiration, ATP generation and membrane potential(*17*). Mitochondria are cellular organelles with a central role in energy production, cellular stress response and apoptosis. Mitochondrial function has long been associated with health and longevity, and mitochondrial dysfunction can cause complex multi-tissue diseases, including metabolic and neurodegenerative disorders(*18, 19*). Activating transcription factor 4 (ATF4) is a key mediator of mitochondrial stress in response to perturbations in mitochondrial proteostasis(*20*). In the liver, ATF4 is up-regulated in response to lifespan-extending interventions such as methionine restriction, rapamycin and acarbose treatment(*21*). In addition, different mitochondrial stressors, affecting mitochondrial translation, OXPHOS stability, mitochondrial membrane potential disruption and impairment of mitochondrial import, activate ATF4, which then up-regulates expression of cytoprotective genes that result in a down-regulation of mitochondrial respiration and activation of the integrated stress response (ISR) through cellular metabolic rewiring(*20*). Moreover, in response to stress, ATF4 induces ATF5(*22*), which activates a programme to rescue mitochondrial activity(*23*). These adaptations lead to increased cellular resistance and protect cells from mitochondrial stress and apoptosis. Importantly, over-expression of the ATF4 target gene FGF21 can extend lifespan in mice, suggesting that the ISR might also ameliorate ageing in mammals(*24*).

In this study, we addressed whether the benefits observed in Irs1KO mice were due to protection from age-associated decline, or instead reflected an age-independent effect. Moreover, we extended the phenotypic characterisation of age-related phenotypes to include male Irs1KO mice, which have previously been shown to be long-lived but have not been extensively studied with respect to ageing pathology. We also deleted IRS1 specifically in liver, muscle, fat and brain tissue of male and female mice and systematically assessed adult survival as well as health parameters in young and old mice. In contrast to global deletion of IRS1, we did not detect lifespan extension in the tissue-specific Irs1KO mice. However, we found that neuron-specific IRS1 deletion was unique in increasing energy expenditure (EE), locomotor activity and insulin sensitivity, specifically in old male mice. Reduced neuronal IIS induced male-specific and age-dependent mitochondrial dysfunction, leading to up-regulation of the ISR in the brain and systemic effects in peripheral tissues.

## Results

### Increased lifespan and health in Irs1KO mice

The effects of reduced IIS on metabolism and longevity are often sex-specific(*2*). For instance, female Irs1KO mice in a C57BL/6J background showed greater lifespan extension than male mice(*7*). Mutant females were also healthier than controls at old age(*15*). The health status of mutant males was not reported. The improved health at old age could indicate protection against the effects of ageing, or reflect an age-independent effect. To address these questions, we measured health parameters in 5 months (young) and 16 months (old) old female and male Irs1KO mice. We set out to perform these experiments using Irs1KO mice in a C57BL/6N background. However, homozygous mutant animals were not born in the expected Mendelian ratio and only 4% of homozygous Irs1KO animals were retrieved from matings between heterozygous females and males (10 males and 13 females out of 510 pups). In an earlier study(*15*), there was also a deficit of Irs1KO pups from double heterozygous matings in the C57BL/6J background, particularly for males (pers. comm.). We therefore generated these animals in a C3B6F1 hybrid background, in which homozygous animals were born in the expected 25% ratio. We assessed the health of young and old male and female Irs1KO mice in the C3B6F1 hybrid background.

#### Lifespan

To determine if lack of IRS1 can also extend lifespan in the C3B6F1 background, we measured survival of hybrid C3B6F1 Irs1KO mice and their wildtype littermates. Loss of IRS1 led to an extension of median lifespan by 7% and 11% and of maximum lifespan, as defined by the 80th percentile age, by 12% and 10% for males (Fig. 1a) and females (Fig. 1b), respectively. The lifespan-extending effect of loss of IRS1 is therefore robust to different genetic backgrounds. Moreover, there was no sex bias in lifespan extension in the C3B6F1 hybrid background (Cox proportional hazard test, sex*genotype interaction P = 0.4844).

**Figure 1:**
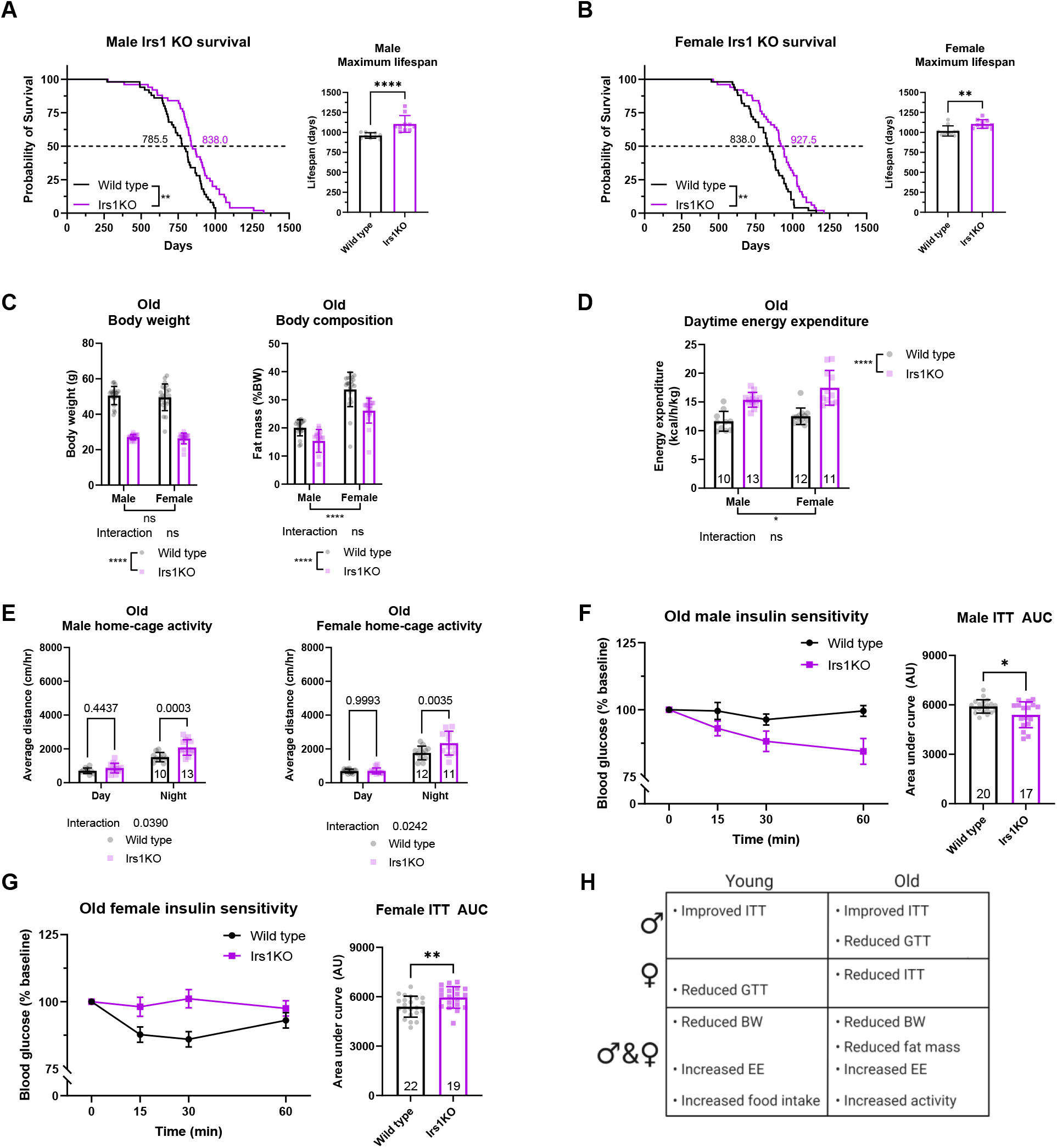
Increased lifespan and improved health parameters in Irs1KO mice. Kaplan Meier Plots depicting survival of **(A)** male and **(B)** female wild type and whole body Irs1KO mice (n=50 biologically independent animals per sex and genotype). **(C)** Body weight and composition of Irs1KO mice was measured at old age (16 months) (Wild type males n=20, Irs1KO males n=17, Wild type females n=21, Irs1KO females n=19). **(D)** Body weight normalised energy expenditure of singly housed old Irs1KO mice during daytime. **(E)** Spontaneous activity of old Irs1KO single housed mice during daytime (inactive phase) and nighttime (active phase) (Wild type males n=10, Irs1KO males n=13, Wild type females n=12, Irs1KO females n=11). Insulin tolerance test (ITT) performed on male **(F)** and female **(G)** Irs1KO mice at old age with respective AUC analysis revealed significantly higher insulin sensitivity in male Irs1KO mice but significantly lower insulin sensitivity in female Irs1KO mice. **(H)** Table summarising the phenotypes unique to and shared between male and female mutant mice. All error bars correspond to standard deviation except for longitudinal insulin sensitivity where standard error of the mean was reported. Number of animals reported at the bottom of the bars or in figure legends. Detailed statistical values found in Table S1.

Insulin mutant animals are not only long-lived but also show resistance to age-related diseases(*15*). Given the replication of the Irs1KO lifespan extension phenotype in a new mouse genetic background, we assessed whether female hybrid Irs1KO mice also presented with improved age-associated outcomes as reported in the original study(*15*), and extended the characterisation to include male Irs1KO mice.

#### Body weight and composition

Irs1KO C3B6F1 mice were dwarves, with a sex-specific reduction in body weight of 46% and 36% in young male and female mice, respectively (two-way ANOVA, sex*genotype interaction P<0.0001; F(1,76)=22.74, Supplementary Fig. 1a). The greater effect of the mutation on male body weight may have been in part attributable to differences in fat content, as young male Irs1KO mice showed a sex-specific reduction in fat mass, with no significant change in females (two-way ANOVA, sex*genotype interaction P=0.0041; F(1,76)=8.76, Supplementary Fig. 1a). Body weight and fat content were reduced in mutant mice of both sexes at old age, with no significant sex and genotype interaction (two-way ANOVA, body weight sex*genotype interaction term P=0.9049; F(1,74)=0.0144, fat content sex*genotype interaction term P=0.1787; F(1,74)=1,844, Fig. 1c). The decreased adiposity of old Irs1KO mice was not due to reduced food intake, because there was no difference in food consumption relative to body weight between old mutant and wild type animals of either sex (Supplementary Fig. 2a). In contrast, young Irs1KO mice of both sexes ate more food relative to their body weight (Supplementary Fig. 1b), while not showing increased fat mass, suggesting changes in energy expenditure.

#### Energy expenditure

Indirect calorimetry showed increased daytime energy expenditure (EE) in young Irs1KO mice of both sexes (Supplementary Fig. 1c), consistent with the hypothesis that body size and body weight-adjusted EE are inversely correlated(*25*). We also found increased EE in old Irs1KO mice of both sexes during daytime (Fig. 1d) and nighttime (Supplementary Fig. 2b). The decrease in adiposity was therefore probably not mediated by increased EE, as young female Irs1KO showed increased EE but no significant difference in adiposity (Supplementary Fig. 1a).

We next measured spontaneous home-cage locomotor activity to investigate if an increase in movement could account for the decreased adiposity observed in Irs1KO mice. Young male and female Irs1KO mice showed no significant difference in activity levels (Supplementary Fig. 1d). However, there was a trend that did not reach significance in nighttime male Irs1KO activity (Supplementary Fig. 1d) that may have contributed to the sex-specific reduction in adiposity observed in young male Irs1KO mice (Supplementary Fig. 1a). Analysis of activity levels in old male and female Irs1KO mice revealed a significant increase in nighttime, but not daytime, activity (Fig. 1e). Thus, increased locomotor activity might contribute to the reduced adiposity of old male and female Irs1KO mutant mice, but it cannot explain the reduced fat mass of young male Irs1KO mice.

Consistent with previous findings in female C57BL/6J Irs1KO mice, female hybrid Irs1KO developed an age-dependent reduction in adiposity(*15*). However, adiposity presented in a sex-specific manner, where male hybrid Irs1KO mice had significantly decreased fat content independent of age. Moreover, we detected a significant increase in locomotor activity in old male and female Irs1KO mice. One potential explanation could be an amelioration of the age-dependent decrease in locomotor activity observed in wild type mice(*26*). Interestingly, this is consistent with the original report in C57BL/6J female Irs1KO mice of a delay in age-dependent loss of locomotor coordination on the rotarod(*15*).

#### Peripheral metabolism

IIS is a major regulator of glucose metabolism(27), and insulin resistance is a causal factor for several age-related pathologies including obesity, type-2 diabetes and metabolic syndrome(*28*). Therefore, we assessed whether Irs1KO mutant mice showed age-related changes in glucose metabolism by performing insulin tolerance (ITT) and glucose tolerance tests (GTT) of young and old Irs1KO mice of both sexes as a read out for insulin sensitivity and pancreatic beta cell function, respectively. Area under the curve (AUC) analysis of ITT revealed that male Irs1KO mice showed significantly increased insulin sensitivity compared to controls at both young (Supplementary Fig. 1e) and old age (Fig. 1f). In contrast, loss-of Irs1 had no effect on insulin sensitivity in young females (Supplementary Fig. 1f) but caused insulin insensitivity in old females (Fig. 1g), consistent with results of female Irs1KO C57BL/6J mice (*15*). There was no difference in glucose clearance in young Irs1KO males (Supplementary Fig. 1g), but old Irs1KO males showed a significantly reduced response to the glucose challenge (Supplementary Fig. 2c). Conversely, young Irs1KO females presented with glucose intolerance (Supplementary Fig. 1h) while there was no significant difference in glucose tolerance in old Irs1KO females (Supplementary Fig. 2d). These findings are consistent with previous reports, which also detected no change in glucose tolerance in 16-month-old female Irs1KO C57BL/6J mice(*15*). Interestingly, glucose tolerance was improved in 23-month-old female Irs1KO C57BL/6J mice(15).

In summary, C3B6F1 hybrid Irs1KO mice showed no sex bias in lifespan or metabolic health parameters such as reduced adiposity, increased EE, and locomotor activity at old age. However, C3B6F1 Irs1KO males showed a sex-specific benefit in insulin sensitivity.

### Generation and validation of tissue-specific Irs1 knockout mice

We next investigated the role of the five major insulin-responsive metabolic organs, liver, muscle, fat, gut and nervous system, in mediating the effect of IRS1 deficiency on murine longevity and health. Irs1KO mice in the C3B6F1 hybrid background showed similar phenotypes as previously reported for C57BL/6J Irs1KO mice(*15*). Therefore, we conducted the tissue specific experiments in the C57BL/6N background, as the LoxP floxed Irs1 allele(*29*) and the tissue-specific Cre drivers were only available in this background. In contrast to the global deletion of IRS1 in the C57BL/6N background, mice with tissue-specific deletion of IRS1 were born in the expected Mendelian ratio, suggesting that development of the animals was not detrimentally affected.

Liver-specific Irs1 knockout mice were generated using Alfp-CreT (lKO)(*30*), muscle-specific (targeting skeletal and cardiac muscle tissue) using Ckmm-CreT (mKO)(*31*), fat-specific (targeting white adipose tissue (WAT) and brown adipose tissue (BAT)) using Adipoq-CreT (fKO)(*32*), gut-specific (targeting small and large intestine) using Villin1-CreT (gKO)(33) and neuron-specific using Syn1-CreT (nKO)(34) mice. Male C57BL/6N mice carrying the corresponding CreT transgenic constructs were mated to LoxP floxed Irs1(29) mutant C57BL/6N females to generate tissue-specific Irs1 knockout mice (CreT/+::Irs1fl/fl) and their corresponding LoxP floxed Irs1 littermate controls (Cre+/+::Irs1fl/fl). We first validated the efficiency of IRS1 depletion by measuring *Irs1* transcript levels by quantitative real-time PCR (Q-RT-PCR) in the corresponding target tissue (Supplementary Fig. 3a-f). This was done using old male and female mice to verify that the depletion of IRS1 is stable throughout life. Primers and Q-RT-PCR conditions were validated with cortical brain samples from global Irs1KO mice (Supplementary Fig. 3a). *Irs1* transcripts were strongly depleted in liver tissue of lKO (Supplementary Fig. 3b), hindlimb muscle tissue of mKO (Supplementary Fig. 3c), BAT tissue (subscapular) of fKO (Supplementary Fig. 3d) and partly depleted in cortical brain tissue of nKO mice (Supplementary Fig. 3e). The partial reduction in the cortex is probably explained by the residual expression of *Irs1* in glial cells, which are not targeted by the Syn1-CreT(*34*). In contrast, we did not detect depletion of *Irs1* transcripts in the small intestine (ileum) of gKO mice (Supplementary Fig. 3f), suggesting that Irs1 mutant cells were outcompeted by wild type cells in the gut epithelium during ageing. Therefore, gKO mice were excluded from further analysis.

We then tested the specificity of the tissue-specific knockout lines by measuring *Irs1* expression levels in untargeted tissues, namely brain cortex for lKO, mKO and fKO (Supplementary Fig. 3g) and liver tissue for the nKO (Supplementary Fig. 3h), respectively. There was no significant difference in the expression level of Irs1 in the non-targeted tissues, demonstrating the specificity of the generated tissue-specific knockout lines. As IRS1 and IRS2 have been suggested to have redundant functions, we used Q-RT-PCR to measure *Irs2* levels in the IRS1 tissue-specific knockout lines. However, except for a slight up-regulation of *Irs2* transcript levels in the liver of male IKO mice (Supplementary Fig. 3b), there was no compensatory up-regulation of *Irs2* gene expression in any of the tested knockout lines (Supplementary Fig. 3a-e). In summary, the four IRS1 tissue-specific knockout lines had efficient depletion of *Irs1* expression, without unspecific effects in other tissues or compensatory regulation of *Irs2* expression. Thus, these lines were suitable for addressing how tissue-specific depletion of IRS1 affects health and longevity.

### Tissue-specific knockout of Irs1 in liver, muscle, fat or neurons does not extend lifespan in mice

#### Lifespan

As global loss of IRS1 results in lifespan extension, we measured survival of tissue-specific Irs1 mutant male and female mice, to address which tissue might underlie the longevity effect. However, there was no lifespan extension detected by log-rank test in male or female lKO, mKO, fKO and nKO (Fig. 2) mice.

**Figure 2:**
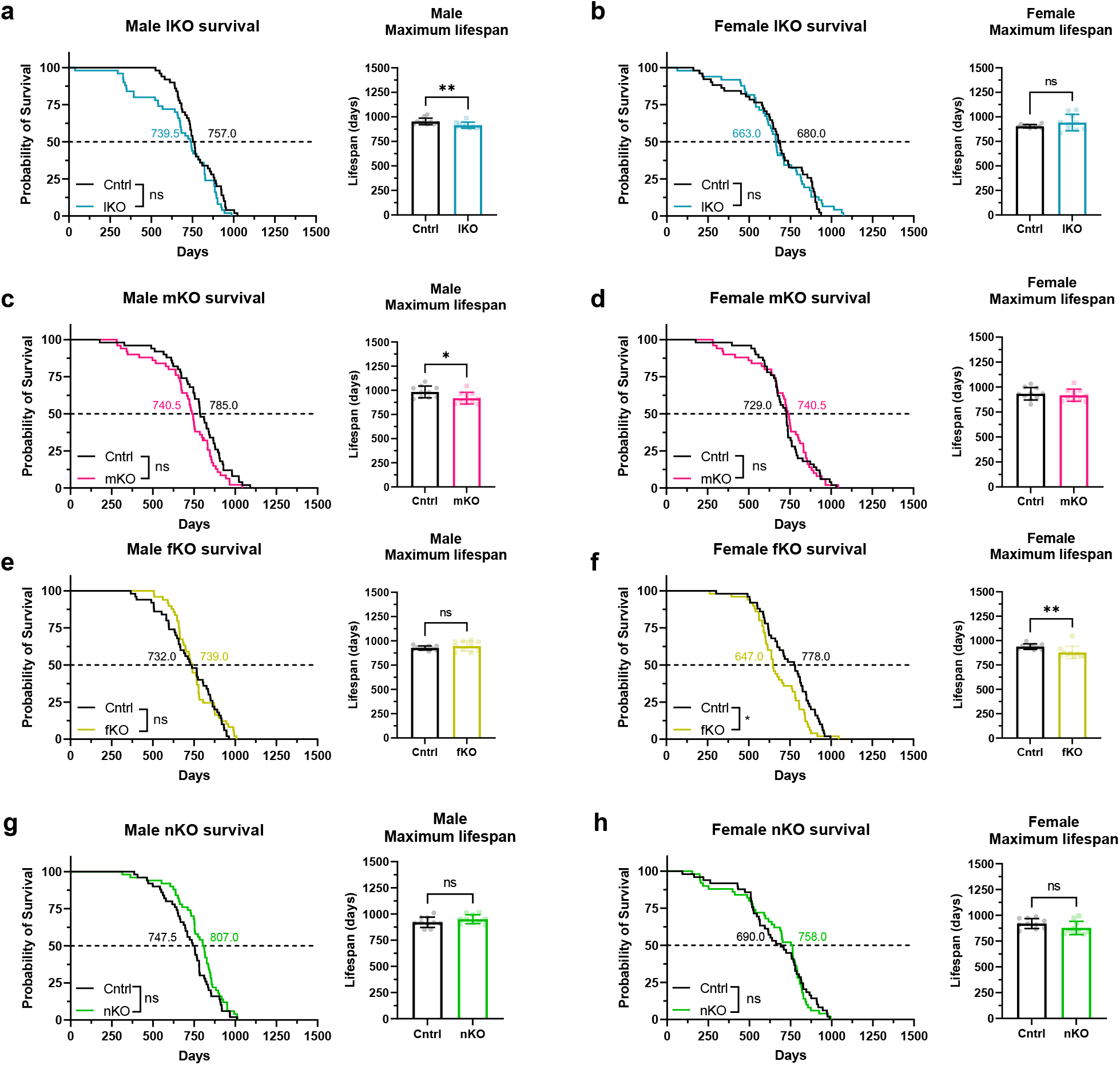
Tissue-specific deletion of IRS1 is not sufficient for lifespan extension. Kaplan Meier plots depicting survival of male and female mice. Log-rank tests were used to compare median survival (labelled on 50% survival probability line in days) and Mann Whitney tests to compare maximum lifespan (inset shows top 20% longest-lived mice) of all mutant mice. **(C)** Male and **(D)** female control (Cntrl) and liver-specific Irs1 knockout (lKO) mice (n=50 biologically independent animals for all groups). **(E)** Male and **(F)** female control (Cntrl) and muscle-specific Irs1 knockout (mKO) mice (n=50 biologically independent animals for all groups). **(G)** Male and **(H)** female control (Cntrl) and fat-specific Irs1 knockout (fKO) mice (n=50 biologically independent animals for all groups, n=49 for male fKO mice). **(I)** Male and **(J)** female control (Cntrl) and neuron-specific Irs1KO (nKO) mice (n=50 biologically independent animals for all groups, n=49 for female control). Detailed statistical values found in Table S1.

In summary, we did not detect lifespan extension in any of the tested Irs1 tissue-specific KO lines, suggesting that reduction of IIS in more than one tissue or in a tissue we have not targeted is required for lifespan extension in mice.

### IRS1 deletion in liver, muscle or fat does not improve health at old age

#### Body weight and composition

In order to address whether tissue-specific deletion of IRS1 would lead to health benefits, we conducted phenotyping of males and females of the Irs1 tissue-specific knockout lines. In contrast to whole body Irs1KO animals, which are dwarfs, there was no difference in body weight in young lKO, mKO or fKO animals (Supplementary Fig. 4a, e and i), indicating that deletion of IRS1 in these tissues did not interfere with overall growth.

The liver stores glucose after a meal as glycogen or converts excess glucose to fatty acids. It also oxidises fatty acids to provide energy for gluconeogenesis during fasting. Moreover, sex affects liver physiology, with significant consequences for systemic metabolism(*35*). IRS1 plays a role in muscle growth and in insulin-stimulated glucose transport into muscle in male mice(*36, 37*), but sex-specific effects of IRS1 in muscle tissue have not been fully characterised. Therefore, we assessed whether muscle contributed to the sex-specific health benefits observed in Irs1KO mice. The contribution of adipose tissue to the physiological insulin response is not certain. However, fat-specific IR knockout mice are protected against age-associated glucose intolerance and insulin insensitivity(*38*), and show enhanced lifespan(*39*). However, the role of a relatively modest IIS reduction in IRS1 deletion and the role of sex have not been assessed.

As we observed sex differences in the metabolic phenotypes of Irs1KO mice, we measured metabolic phenotypes in male and female lKO mice. lKO mice showed a sex-specific reduction in body weight only in old males (two-way ANOVA, sex*genotype interaction P=0.0015;F(1,45)=11.38, Supplementary Fig. 4c), which was probably due to a sex-specific decrease in fat mass (two-way ANOVA, sex*genotype interaction P=0.0284;F(1,45)=5.132, Supplementary Fig. 4d). Male and female mKO mice showed no change in body weight (Supplementary Fig. 4e and g), but a significant increase in fat mass relative to body weight at both young and old age in both sexes (Supplementary Fig. 4f and h). These results are consistent with findings using muscle-specific IR knockout animals, which also showed increased fat mass and reduced muscle mass(*40*). Although male and female fKO showed no change in body weight (Supplementary Fig. 4i and k), we measured a significant reduction in fat mass at young and old age in both male and female fKO mice (Supplementary Fig. 4j and l), suggesting that, similar to fat-specific IR knockout mice(*38, 41*), fKO mice have a reduced WAT mass. Changes in body weight and composition were not a result of altered food consumption as no significant differences were observed in young and old lKO, mKO or fKO animals (Supplementary Fig. 4m-r). There has been controversy whether IIS in fat can modulate feeding, as one report found a significant increase in fat-specific IR knockout mice(*41*), while another study found no difference(*38*). We did not detect any significant changes in food intake of young or old fKO mice of either sex (Supplementary Fig. 4q and r), which might be due to the less severe down-regulation of IIS upon loss of IRS1 compared to the IR.

#### Energy expenditure

There was no significant difference in EE in young or old male or female lKO mice (Supplementary Fig. 5a-d). IRS1 deficiency in muscle and fat tissue did not lead to a significant difference in EE in young or old male and female mKO and fKO mice (Supplementary Fig. 5e-l). Spontaneous locomotor activity did not reveal any significant difference in young or old male and female lKO (Supplementary Fig. 6a-b). Young male mKO mice showed increased spontaneous locomotor activity specifically during nighttime (Supplementary Fig. 6c), while no change was observed in mKO females or old mKO males (Supplementary Fig. 6d). Loss-of IRS1 in the fat did not affect spontaneous activity of young or old male or female mice (Supplementary Fig. 6e and f).

#### Peripheral metabolism

Insulin sensitivity of young or old male and female lKO mice was not changed compared to control animals (Supplementary Fig. 7a-d). Insulin sensitivity of mKO mice was also not significantly changed (Supplementary Fig. 7e-h), consistent with data from muscle-specific IR and IGF1R knockout mice, which had reduced activated IRS1 levels but no change in insulin sensitivity (31,42). Young male fKO mice had a significantly reduced insulin sensitivity (Supplementary Fig. 7i), while young females showed no significant difference (Supplementary Fig. 7j). Insulin sensitivity of old male fKO mice was not significantly changed (Supplementary Fig. 7k), however, females had significantly reduced insulin sensitivity (Supplementary Fig. 7l).

Consistent with previous glucose tolerance test studies(*43*), male lKO mice showed reduced glucose tolerance at young (Supplementary Fig. 8a), but not old age (Supplementary Fig. 8c). However, a hyperinsulinemic-euglycemic clamp study in another report found no evidence for any difference in glucose tolerance in young male lKO mice (*44*). In contrast to the males in our study, female lKO mice showed no difference in glucose tolerance at young or old age (Supplementary Fig. 8b and d). Young male and female mKO animals showed a significant reduction in glucose tolerance (Supplementary Fig. 8e and f), which was surprising, given that IR or IRS1+2 knockout mice did not show this effect (*31, 37*). However, the reduction in glucose tolerance was also observed in old male (Supplementary Fig. 8g), but not in old female mKO mice (Supplementary Fig. 8h). Glucose tolerance was significantly reduced in young female fKO mice (Supplementary Fig. 8j), but unaffected in young males (Supplementary Fig. 8e). Old female fKO mice were more sensitive to glucose than controls (Supplementary Fig. 8l), but no difference was detected in old males (Supplementary Fig. 8k).

In summary, loss of IRS1 in the liver affected body composition and glucose tolerance in males. Loss of IRS1 in muscle reduced lean mass and glucose tolerance at young age of both males and females, while locomotor activity was increased only in young males. Fat tissue specific loss of IRS1 function mostly affected peripheral metabolism in females. Thus, loss of IRS1 affects peripheral metabolism in a tissue and sex specific manner.

### Neuron-specific deletion of Irs1 causes male-specific improvement in metabolic health

The brain is a central mediator of metabolic function(*45*), and reduced IIS in the brain has been associated with changes in body weight, fat mass, glucose metabolism and feeding behaviour(*46*). However, peripheral, off-target recombination has been reported in kidney, pancreas and muscle of the Cre-driver lines used in these studies(*47*). The Syn1 Cre line used here is specific to neurons(34) and allowed us to specifically study the effects of reduced IIS in the brain on peripheral metabolism. Therefore, male and female nKO mice at young and old age were phenotyped. There was a slight sex-specific reduction in body weight in old male nKO mice (two-way ANOVA, sex*genotype interaction P<0.0172; F(1,51)=6.069, Fig. 3a) with no significant change in body composition of either young (Supplementary Fig. 9a) or old nKO (Fig. 3a) mice of either sex. Brain insulin signalling has been implicated in feeding and satiety(*48*). Thus, we measured food consumption in nKO mice, but did not detect any differences in young (Supplementary Fig. 9b) or old (Supplementary Fig. 10a) mice of either sex. Previous studies have uncovered a direct role of the brain in mediating EE through peripheral tissues such as BAT(*49, 50*). In young animals, there was no effect on EE in either male or female nKO mice (Supplementary Fig. 9c). However, EE was specifically increased in old male, but not female nKO mice during daytime (Fig. 3b) and nighttime (Supplementary Fig. 10b), indicating that nKO mice show an age-dependent and sex-specific increase in EE. Given the role of neuronal IIS in locomotor activity(51) we next measured spontaneous home-cage activity. Locomotor activity was not changed in young nKO males (Supplementary Fig. 9d), but was increased in young females during both daytime and nighttime. In contrast, old nKO males showed a significant increase in activity only during nighttime, while there was no change in old nKO females (Fig. 3c). The male-specific increase in activity was not observed during daytime, suggesting that the effect seen at old age was not due to general hyperactivity or involuntary movements. As neuronal IIS function has been implicated in mediating insulin sensitivity through circuitry with peripheral metabolic organs(*51*), we assessed insulin sensitivity in young and old nKO mice of both sexes. Consistent with male Irs1KO mice, insulin sensitivity was increased in young (Supplementary Fig. 9e) and old (Fig. 3d) male nKO mice. In contrast, young female nKO mice showed a significant reduction in glucose clearance (Supplementary Fig. 9f), which was not present at old age (Fig. 3e). Importantly, Syn1 Cre control mice did not show any significant changes in the corresponding phenotypes, suggesting that the observed differences in nKO mice are not caused by neuronal Cre expression (Supplementary Fig. 11a-j). Thus, IRS1 deletion in neurons was sufficient to induce male-specific benefits in metabolic outcomes that in part recapitulated the phenotypes of the global Irs1KO. Moreover, nKO mice did not show the negative consequences of globally reduced IIS in the C57BL/6 background such as reduced body size, viability and glucose tolerance (Supplementary Fig. 10c and d).

**Figure 3:**
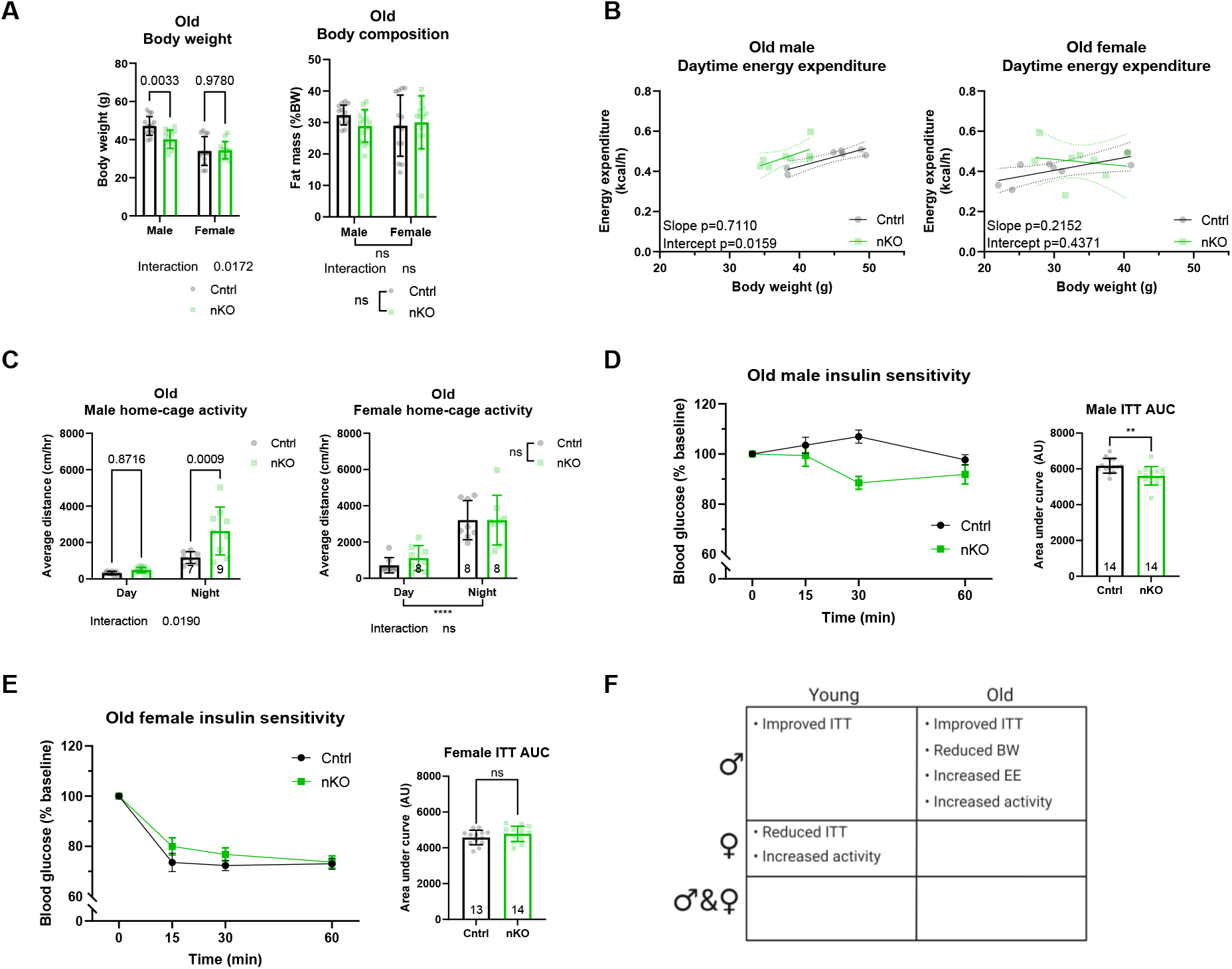
Neuron-specific Irs1 knockout (nKO) mice show male-specific improvement in metabolic health. **(A)** Body weight and composition of nKO mice was assessed at old age (16 months) (n = 14 biologically independent animals for all groups). **(B)** Daytime energy expenditure of male (control n=7 and nKO n=8) and female (n=8 female control and nKO) mice was analysed by linear regression of energy expenditure by body weight (ANCOVA). **(C)** Plotted spontaneous home-cage activity of old male (control n =7 and nKO n=9) and female (control n=7 and nKO n=9) single housed nKO mice showed a nighttime specific increase in activity of male nKO mice. **(D)** Analysis of insulin tolerance test (ITT) curves and AUC values of old male nKO showed a significant improvement in insulin sensitivity in nKO mice compared to controls. **(E)** Analysis of ITT curves and AUC values of old female nKO did not reveal any significant difference compared to controls. **(F)** Table summarising the phenotypes unique to and shared between male and female mutant mice, highlighting the enrichment of male-specific phenotypes in nKO mice. All error bars correspond to standard deviation except for longitudinal insulin sensitivity where standard error of the mean was reported. For ANCOVA analysis the 95% confidence interval is plotted. Number of animals reported at the bottom of the bars or in figure legends. Detailed statistical values found in Table S1.

In summary, while loss of IRS1 in liver, muscle or fat did not improve health at old age, deletion of IRS1 in neurons was sufficient to cause similar health benefits in old males as observed in Irs1KO mice. Thus, reduced neuronal IIS may contribute to the improved insulin sensitivity, increased EE and locomotor activity of male whole body Irs1KO mice.

### Loss of IRS1 function caused mitochondrial dysfunction in the brain of old males

We next asked which molecular mechanisms in the brain might underlie the improved metabolic health of nKO males. Gene expression studies in liver tissue of Irs1KO mice showed regulation of genes associated with oxidative phosphorylation and the tricarboxylic acid cycle(15), and activation of FOXO1 in the liver resulted in a reduction in the activity of the mitochondrial electron transport chain(*17*), suggesting a change in mitochondrial function upon reduced IIS. Mitochondrial function in neurons is essential for neurotransmission, synaptic maintenance and calcium homeostasis(*18*). Mitochondrial function has been implicated in the longevity-dependent response to IIS reduction in the brain of *Drosophila(52)*. However, the effect of reduced IIS on mitochondrial function has not yet been investigated in the mammalian brain. Therefore, we assessed the effect of loss of IRS1 in the brain on oxidative phosphorylation (OXPHOS) by performing high-resolution respirometry on permeabilized brain tissue of young and old Irs1KO mice. There was no difference in basal respiration in young (Supplementary Fig. 12a) or old (Fig. 4a) Irs1KO mice of either sex, suggesting that loss of IRS1 function did not affect mitochondrial function in baseline conditions. Next, we determined the mitochondrial spare respiratory capacity, which indicates how much capacity a cell has to deal with acute additional energy demands. Neuronal reduction in spare respiratory capacity has been linked to age-associated neurodegenerative disorders such as Parkinson’s disease(*53–55*). Old male Irs1KO mice showed a significantly reduced mitochondrial spare respiratory capacity (Fig. 4c). In contrast, old females or young animals showed no change in mitochondrial spare respiratory capacity (Supplementary Fig. 12b). To address whether this phenotype was caused specifically by the lack of IRS1 in neurons, we also measured spare respiratory capacity in nKO mice. Consistent with the results in Irs1KO mice, nKO mice showed a male-specific reduction in mitochondrial spare respiratory capacity at old age (Figure 4b, d), and this was unchanged in Syn1 Cre control mice (Supplementary Fig. 13a and b), indicating that this phenotype was specific to neuronal reduction of IIS. Thus, reduced neuronal IIS lowered mitochondrial spare respiratory capacity in old male Irs1KO and nKO mice, suggesting a low level of mitochondrial stress as basal respiration was unaffected(*56*).

**Figure 4:**
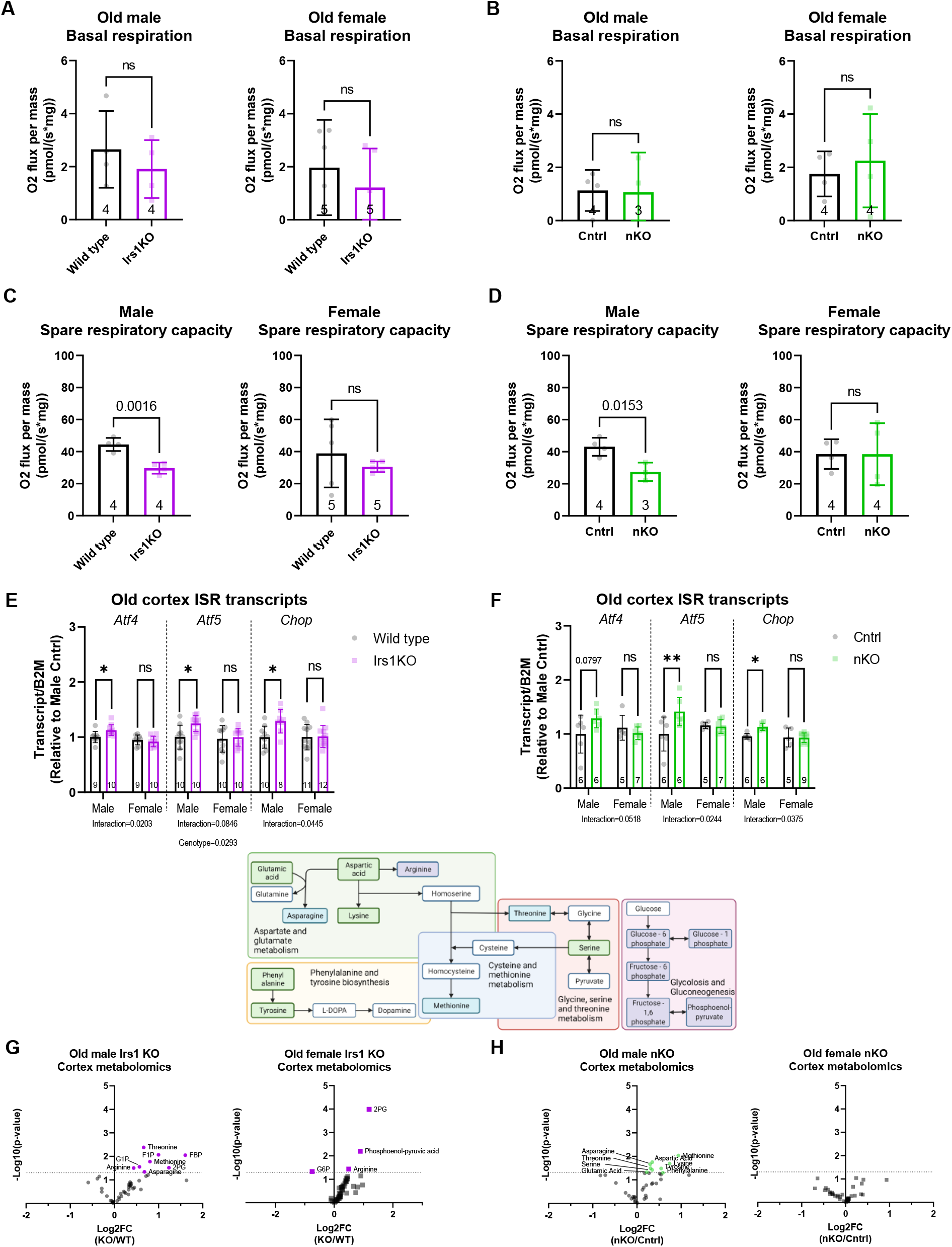
Investigation of mitochondrial function implicated the integrated stress response. **(A)** Measurement of basal oxygen consumption of brain tissue from old (19 months) Irs1KO mice did not detect any differences (male wild type and Irs1KO n=4, female wild type and Irs1KO n=5). **(B)** Measurement of basal oxygen consumption of brain tissue from old (22 months) nKO mice did not detect any differences (male control n=4 and nKO n=3, female control and nKO n=4). **(C)** Spare mitochondrial capacity in brain tissue of old Irs1KO mice was measured, after titration of a protonophore (FCCP) and subtracting the basal respiration, revealing significant reduction in male but not female Irs1KO brains (male wild type and Irs1KO n=4, female wild type and Irs1KO n=5). **(D)** Spare mitochondrial capacity in brain tissue of old nKO mice also revealed significant reduction in male but not female nKO brains (male control n=4 and nKO n=3, female control and nKO n=4). **(E)** Quantitative real-time PCR performed to measure transcript levels of several integrated stress response (ISR) markers on brain tissue of old Irs1KO mice and their wild type littermates found significant sex-specific up-regulation of ISR in male Irs1KO mice (male wild types n=9 and Irs1KO n=10, female wild types n=9 and Irs1KO n=10). **(F)** Transcripts of ISR markers were measured in brain tissue of old (16 months) nKO mice and their control littermates found significant sex-specific up-regulation of ISR in male nKO mice (male control and nKO n=6, female control n=5 and nKO n=7). **(G)** Semi-targeted metabolomics revealed only up-regulation of some metabolites in old brain tissue of Irs1KO mice compared to littermate wild types (male wild type n=5 and Irs1KO n=6, and female wild type and Irs1KO n=6). **(H)** While metabolomics analysis in old nKO mice also revealed only up-regulated metabolites, but exclusively in male brain tissue (male control and nKO n=6, female control and nKO n=5). Included above the volcano plots is a schematic of the metabolic pathways affected by reduced IIS in the brain, highlighting metabolites up-regulated in old Irs1KO only (purple box), old nKO males only (green boxes) and old Irs1KO and nKO males (blue boxes). All error bars correspond to standard deviation. Detailed statistical values found in Table S1. Full metabolites measured in Table S2.

### Lack of IRS1 causes a male-specific induction of the integrated stress response in aged brains

Impaired mitochondrial function has been shown to activate Atf4 signalling(*20*) and thereby the ISR(*57, 58*). Moreover, an increase in ATF4 activity has been identified as a common feature of interventions that increase lifespan in mice(*21*). Therefore, we measured *Atf4* transcript levels by Q-RT-PCR in the brains of Irs1KO mice, as neuronal ATF4 protein is usually rapidly degraded(*59*). *Atf4* mRNA levels were unchanged in female mice, but specifically increased in the brain of old male Irs1KO mice (Fig. 4e and Supplementary Fig. 12c), consistent with the male-specific mitochondrial dysfunction at old age. In line with this finding, *Atf5* levels were also only significantly up-regulated in old male Irs1KO mice (Fig. 4e). Expression level of *Chop*, an *Atf4* target gene commonly up-regulated in long lived mice(*21*), was also increased specifically in male Irs1KO mice (Fig. 4e). These results suggested that lack of IRS1 induced ISR in the brain of old males. To address whether deletion of IRS1 in neurons was sufficient to activate the ISR, we measured *Atf4, Atf5* and *Chop* expression in the brain of nKO mice. As in Irs1KO mice, no significant change in expression in these genes was observed in young male or in female nKO mice (Supplementary Fig. 12d). However, *Atf5* and *Chop* were significantly up-regulated in old male nKO mice and there was a tendency for *Atf4* to be up-regulated (Fig. 4f). Moreover, consistent with the lack of mitochondrial dysfunction in Syn1 Cre mice, we did not detect any difference in the expression of the ISR marker genes (Supplementary Fig. 13c). Thus, our data suggest that neuronal loss of IRS1 is sufficient to trigger the ISR specifically in the brain of old male mice.

### Loss-of neuronal IRS1 leads to male-specific metabolic adaptations during ageing

The ISR pathway has been shown to trigger ATF4-dependent cytoprotective metabolic adaptations(*20*), which are particularly relevant for metabolic rewiring in response to mitochondrial stress(*60–62*). Therefore, we used targeted metabolomics on mouse brains to measure metabolite levels that have been associated with the ISR response. Methionine, asparagine and threonine levels were increased specifically in the brain of old male but not female Irs1KO mice (Fig. 4g). These findings are consistent with previous reports that showed up-regulation of amino acid biosynthesis pathways in response to ISR, specifically threonine(*20*), methionine(*63*), glycine and serine(*64*). In contrast, these amino acids were not increased in the brain of young Irs1KO males (Supplementary Fig. 12e). Similar to the results observed in Irs1KO mice, threonine, methionine, glycine and serine were increased specifically in the brain of old nKO mice (Fig. 4h) but not young nKO mice (Supplementary Fig. 12f), suggesting that these metabolic changes are the results of reduced IIS in neurons. The observed metabolic changes are consistent with the hypothesis that loss of IRS1 in neurons of males during ageing triggers the ISR which then induces a cytoprotective metabolic programme.

### IRS1 deletion in peripheral tissues did not induce ISR gene expression

To assess if deletion of IRS1 in tissues other than neurons also induces the ISR, we measured *Atf4, Atf5* and *Chop* transcript level in the liver, muscle and adipose tissue of the respective tissue-specific Irs1 knockouts. In contrast to the findings in the nKO males, ISR marker genes were not up-regulated in the liver, muscle or BAT of old lKO, mKO or fKO mice respectively (Supplementary Fig. 14 a-c), suggesting that neuronal tissue is particularly susceptible to IRS1-deletion mediated ISR.

In summary, neuronal deletion of IRS1 caused a male-specific reduction in mitochondrial spare respiratory capacity, activation of Atf4 signalling and metabolic adaptations consistent with an activated ISR during ageing.

### Neuronal IRS1 deletion induces sex-specific mitochondrial ISR in non-affected tissues

Previous studies in C. elegans have reported that cell-non-autonomous signals due to electron transport chain disruption in neurons are sufficient to lead to lifespan extension by activating mitochondrial stress in the intestine(*65*). Whether mitochondrial ISR induction in one tissue sends stress signals in non-affected tissues in mammals is still unclear. An in-depth study using a mitochondrial myopathy mouse model characterised the temporal progression and inter-tissue response to muscle mitochondrial ISR, such as increased brain uptake of glucose, but did not detect up-regulation of mitochondrial ISR in non-affected tissues(*66*). We asked if IRS1-mediated ISR activation in the brain can activate a mitochondrial ISR signal in peripheral tissues. We measured ISR marker transcripts in liver, muscle, WAT, BAT and intestine (Fig. 5 a-e) in old male and female nKO mice. We detected a sexually dimorphic *Atf4* and *Chop* signal in the muscle, where transcripts were significantly up-regulated in nKO males but down-regulated in females (Fig. 5 b). Moreover, we found that WAT of nKO males presented with a sex-specific up-regulation of *Atf5* and *Chop* levels (Fig. 5 c). However, liver, BAT and gut ISR markers were not significantly changed (Fig. 5 a, d-e). These data suggest that the brain may be secreting a factor that is initiating mitochondrial ISR in some peripheral tissues. We tested whether IRS1 deletion in liver, muscle and fat tissue-specific Irs1KO mice led to an up-regulation of ISR markers in a non-affected tissue such as the brain (Supplementary Fig. 15 e) but we did not detect any evidence of ISR activation.

**Figure 5:**
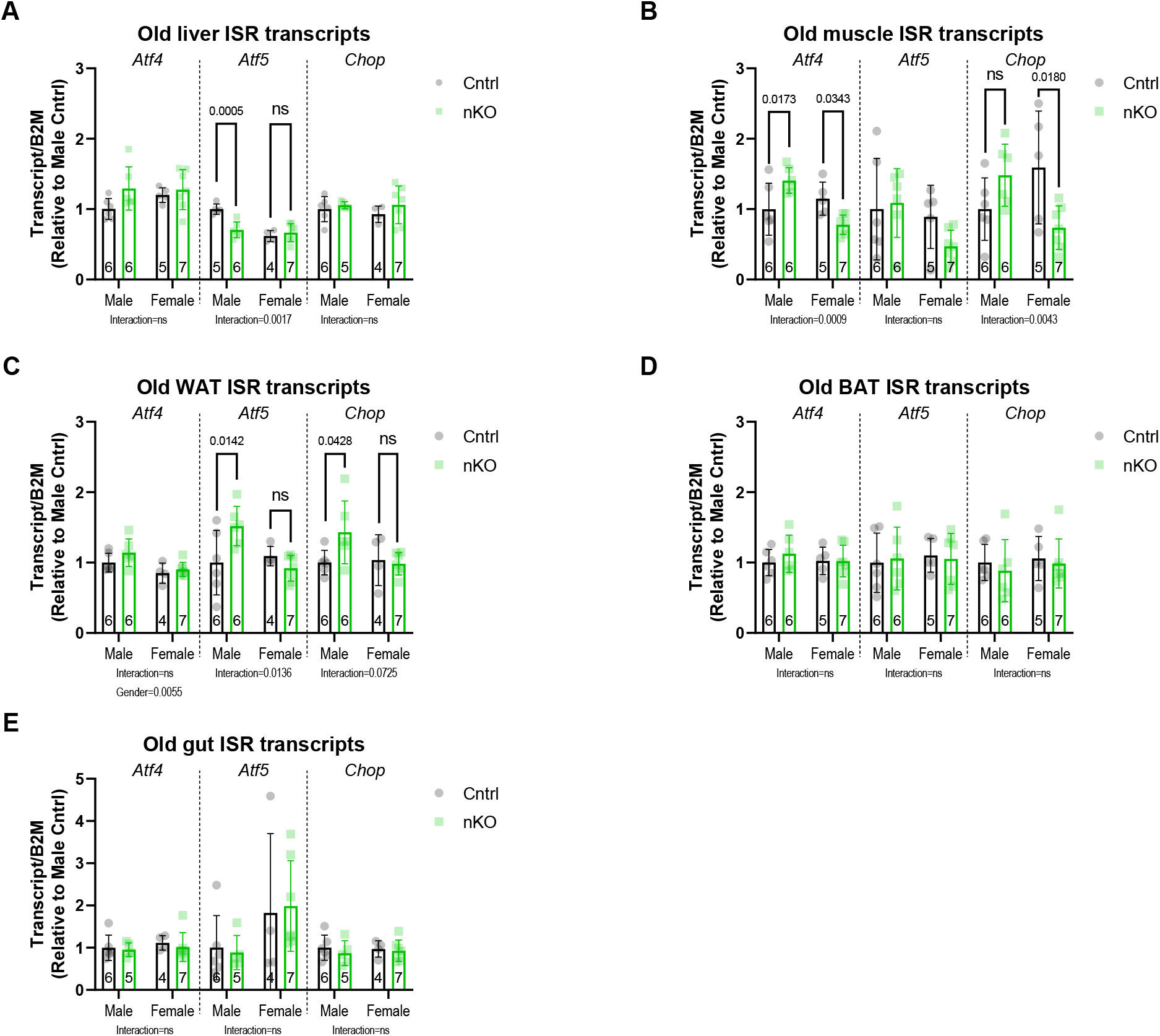
Sex-specific mitochondrial ISR can be activated in non-affected tissues. Quantitative real-time PCR performed on cortex of neuron-specific Irs1 knockout mice in old (16 months) mice targeting various integrated stress response (ISR) markers. **(A)** Liver ISR transcripts in nKO mice and their control littermates revealed a sex-specific downregulation in hepatic Atf5 levels of male nKO mice. **(B)** Muscle Atf4 and Chop transcripts were significantly upregulated in nKO males but significantly downregulated in females, while Atf5 levels were unchanged. **(C)** White adipose tissue Atf5 transcripts were significantly upregulated in nKO males in a sex-specific manner, while Chop levels showed a trend that did not reach significance. Brown adipose tissue **(D)** and gut **(E)** ISR transcripts were not affected in nKO mice. Detailed statistical values found in Table S1.

## Discussion

Our study provides a systematic study into the tissue-specific role of IRS1 deletion on longevity. We find that IRS1 deletion is a robust intervention capable of longevity induction in different mouse strains, suggesting that mechanisms leading to longevity are independent of genetic background and are broadly relevant to the ageing process. Tissue-specific IRS1 deletion in major insulin-sensitive organs was insufficient to extend lifespan. Specific IRS1 deletion in liver, muscle and fat tissue did not improve health at old age in male or female mice, whereas neuron-specific IRS1 deletion led to male-specific increase in health outcomes that recapitulate phenotypes of Irs1KO mice. These findings highlight the importance of neuronal IIS and sex in modulating health outcomes in mice. Furthermore, we found that reduced neuronal IIS led to male-specific mitochondrial dysfunction, activation of Atf4 signalling and metabolic adaptations consistent with an ISR response at old age. Importantly, we find evidence for mammalian mitochondrial ISR spread from the brain to peripheral organs through a sex-specific mechanism in old mice.

We employed a robust C3B6F1 hybrid mouse strain to mitigate confounds of inbred lines on complex phenotypes such as longevity(*67*). There was no effect on hybrid Irs1KO viability, and progeny were born in a Mendelian ratio. We replicated previous reports of Irs1KO mice leading to lifespan extension and improved health parameters in a new robust hybrid background. Previous reports have found that the mouse genetic background affects the extent of lifespan extension in dietary, genetic or pharmacological longevity interventions(*68*). Initially, we generated a global IRS1 deletion model under the ubiquitous Actin-Cre driver. However, there was a depletion of Actin Cre +/T::Irs1 fl/fl pups generated from Actin Cre +/+::Irs1 fl/+ and Actin Cre +/T::Irs1 fl/fl matings, suggesting reduced viability of IRS1 deletion mice. Nor could we successfully breed C57BL/6N Irs1KO mice, because Irs1KO progeny dropped in the third generation of backcrossing to approximately 4%. Haploinsufficiency in the IGF1R gene in the 129Sv mouse background led to a significant lifespan extension in females (33%) but only a trend in males (16%)(*69*). In contrast, Bokov and colleagues repeated the experiment in a C57BL/6 background, and found only a modest lifespan extension in females (5%) and males showed a small but insignificant reduction(*70*). Interestingly, when Bokov and colleagues tried the same intervention in the hybrid 129SBL6F1 background, no lifespan extension was detected in males or females(*70*). These findings suggest that reducing IIS in the C57BL/6 background may lead to a strain-specific sexual dimorphism leading to benefits in females but adverse effects in males. These two studies highlight the importance of replication and impact of inbred mouse strains on ageing studies and the effect of sex on reduced IIS interventions. Taken together, these reports fill us with confidence that the effect of IRS1 deletion on longevity is robust and is targeting biological mechanisms of ageing contrary to rescuing a deficit in an inbred mouse line.

The tissue-specific role of reduced IIS has been investigated extensively in the context of metabolic disorders to study the consequences of tissue-specific insulin resistance(*27*), but the effect of tissue-specific IIS disruption on lifespan is unknown. We measured the lifespan and healthspan of lifelong and tissue-specific IIS reduction by deletion of IRS1 in the liver, muscle, fat and brain of male and female mice. We found no evidence for significant extension of lifespan. Conversely, in *Drosophila* the causal mechanisms that contribute to extended lifespan are tissue-specific(*52*), potentially due to the lack of paralogues that could rescue pathway activity. Global reduction of IIS by deletion of IIS proteins upstream in the pathway, such as IR, IGF1R or IGF1 lead to early mortality(*71, 72*). However, partial disruption, either by targeting specific tissues such as fat(*39*) or brain tissue(*73*), or proteins that do not abrogate the pathway(*7*), can lead to lifespan extension. Recent reports of ectopic recombination in the central nervous system and in peripheral metabolic organs may confound the fat-(*74*) and brain-specific(*47*) findings respectively. Interestingly, when IR was deleted in all peripheral tissues after adulthood no lifespan extension was observed, but a large and significant reduction in male lifespan was detected(*75*). This finding again highlights the sensitivity of C57BL/6 males to reduced IIS, but also suggests that IIS reduction during development plays a role in lifespan modulation. Therefore, we built on previous approaches by using a constitutively active but partial IIS disruption, by deleting the IRS1 protein, in tissue-specific models with a proven record of specific recombination and confirmed by q-RT-PCR. Our findings suggest that either a tissue not tested in our study or multiple tissues are necessary for tissue-specific IRS1 deletion mediated lifespan extension.

We found that the deletion of IRS1 specifically in neurons led to improved systemic metabolic changes at old age, without detrimental effects on body size and viability observed in global C57BL/6J Irs1KO mice. Neuronal IIS reduction led to enhanced insulin sensitivity, increased EE and improved motor activity, specifically in old male mice. However, these changes were insufficient to extend lifespan. In mammalian models of neurodegenerative disease, manipulation of IIS in the central nervous system improves cellular resilience, especially with regards to proteotoxic stress(*76*). The role of IIS in the central nervous system in physiological ageing is still controversial in mammals(*77*). In *C. elegans* neuronal IIS reduction can induce longevity by increasing the resistance of neurons to free radicals(*78*) or by maintenance of mitochondrial integrity though non-cell-autonomous mechanisms. The healthspan benefits we observed in nKO mice are consistent with data obtained from fly models(*79, 80*). Moreover, similar changes in insulin sensitivity and locomotion were obtained from a study where IR was deleted only in male mice and specifically in the hypothalamus under high fat diet conditions(*51*). One possibility for the sex difference is the age-associated and tissue-specific difference in activation of the IIS pathway in male and female C57BL/6 mice(*81*). However, given the interaction of mouse background and IIS, this needs to be tested further in studies that include both male and female mice in different strains.

Previous studies found a reduction in hepatic mitochondrial function in response to IIS reduction but did not disclose the sex of mice used(*17*). Our study found that neuronal IRS1 deletion is sufficient to reduce mitochondrial spare respiratory capacity and up-regulate ISR in male brains in a sex-specific manner. The reduction in mitochondrial capacity without a change in basal mitochondrial respiration could suggest that physiological neuronal function is not impaired, but mitochondria are stressed(*56*). Indeed, the activation of mitochondrial ISR is a sign of a cellular response to stress in order to promote resilience(*20*). Previous reports investigated the effects of brain mitochondrial stress only in male mice, and found it sufficient to induce a systemic endocrine signal(*82*), but the role of sex and the effect of brain mitochondrial ISR on metabolic health and ageing remains to be elucidated. Another potential mechanism leading to systemic effects in non-affected tissues of nKO mice could be the presence of neural circuits innervating specific peripheral organs. Interestingly, although we observed similar systemic metabolic phenotypes in female Irs1KO mice, we could not detect mitochondrial dysfunction or a similar ISR response in female brain tissue. This suggests that the female phenotypes may be mediated through a different mechanism, or that the ISR in females functions in a different way. Indeed, a previous report of mitochondrial DNA-damage-induced mitochondrial stress in muscle tissue revealed sex differences in muscle free amino acids in aged mice(*83*), but sex differences in other parameters were not assessed. Mammalian models of nutrient-stress-induced systemic ISR implicate an endocrine signal leading to a male-specific lifespan(*84*) and healthspan(*85*) improvement, providing some evidence of sex-specificity in this pathway. Unfortunately, most previous studies in the ISR field only studied males, but our report suggests that sex as a variable should be considered in future studies.

### Limitations

We acknowledge several limitations in this study. First, given that all our interventions were constitutive, we cannot be certain to what extent the developmental effect of reduced IIS contributes to the observed phenotypes. Although we could not detect changes in energy expenditure, locomotor activity, ISR transcripts or mitochondrial function at young age, we cannot rule out lingering developmental effects later in life. Time-specific genetic tools could target the IIS intervention to adulthood and help eliminate any developmental confounds. Second, energy expenditure in Irs1KO mice is confounded by their reduced body size. The increased energy expenditure could be partly attributable to the body size difference, even with normalisation to lean mass or body mass, since we cannot adjust for size by ANCOVA because the difference in the range of values between wild type and mutant mice is too large. However, the finding that nKO mice, where we could adjust the data using ANCOVA, showed a similar phenotype might suggest that this phenotype is not simply caused by differences in body weight. Finally, we detected a significant improvement in insulin sensitivity of young male Irs1KO mice, whereas a previous hyperinsulinemic-euglycemic clamp study found evidence of insulin resistance in the muscle of Irs1KO mice(*36*). The clamp employed is a more robust method for measuring insulin sensitivity as a distinction between liver, fat, and muscle insulin resistance can be made. One potential explanation of this discrepancy could be the difference in mouse background and age in the two studies. The study(*36*) did not report the mouse strain, but they did mention the male Irs1KO mice used were 6 weeks old in contrast to males in this study which were 6 months old. The young wildtype hybrid C3B6F1 male mice did not respond to insulin at 6 months of age, which may have exacerbated the difference between young Irs1KO and their littermates (Supplementary Fig. 1e). However, young control C57BL/6N male mice responded to intraperitoneal insulin injection at 3 months of age (Supplementary Fig. 7), potentially highlighting the role of mouse background and age in this metabolic outcome.

In conclusion, we find evidence for a unique and causal role of neuronal IRS1 deletion in triggering systemic health benefits in males that recapitulate those found in Irs1KO mice in a sex-specific manner. Moreover, these data suggest that males and females respond differently to ISR in response to IRS1 deletion, and evidence for sex-specific ISR activation due to nutrient stress has been reported previously(*85*). However, it is currently unclear to what extent is ISR sex-specific in mammals and how different tissues react to the induction of ISR. Taken together, our study reveals details of how reduced IIS triggers a global response across the organism, some of which is tissue-specific and some sex-specific, reflecting a combination of local and systemic cues. Moreover, the newly identified sex-specific mechanisms in response to reduced neuronal IIS provide potential avenues for therapeutic interventions. However, some tissues were not included in this study and they may respond differently with different outcomes for the whole organism. We suggest that future research should investigate both sexes when studying the ISR pathway in response to various stressors and in a tissue-specific manner.

## Materials and Methods

### Mouse experiments and animal care

#### Mouse Generation

*Irs1* knockout (Irs1KO) mice were generated previously(*15*), and kindly provided by Dr. Dominic J. Withers. Previous data of Irs1KO mice were generated in the C57BL/6J background(*15*). However, due to genetic mutations in the C57BL/6J line that may confound metabolic traits, we chose to generate the Irs1KO mice in the C57BL/6N background(*86*). After embryo transfer of Irs1KO embryos from C57BL/6J into C57BL/6N mice, we observed Irs1KO progeny to drop dramatically in the third generation to approximately 1% Irs1KO mice being born. Following attempts to optimise diet and housing conditions, Irs1KO progeny continued to stay far below the expected ratio in subsequent generations (∼7% Irs1KO mice with normal levels of heterozygous and wild type mice observed in litters). Additionally, we tried to generate a Cre-specific whole body Irs1KO in the C57BL/6 background via the Actin Cre driver line. However, attempts to generate Actin Cre +/T:: Irs1 fl/fl led to similarly low rates of Irs1KO mice (∼2%). Therefore, we generated the Irs1KO mouse in a more robust C3B6 hybrid mouse background(*67*). *Irs* KO mice were backcrossed for 4 generations into the C57BL/6N (C57BL/6NCrl, Charles River) and C3H/HeOuJ (RRID: IMSR_JAX:000635, The Jackson Laboratory) background. Heterozygous C3H/HeOuJ *Irs1* -/+ females were mated with heterozygous C57BL/6N *Irs1*-/+ males to generate hybrid C3B6F1 whole body Irs1 knockout (Irs1KO) mice.

*Irs1loxP mice* were generated as previously described(*29*). For tissue-specific knockout of *Irs1, Irs1loxP/loxP* mice were crossed with mice expressing Cre-recombinase under the control of the mouse albumin enhancer and promoter and the mouse alpha-fetoprotein enhancers (*AlfpCre* mice(*30*)), the control of the creatine kinase promoter (CkmmCre mice(*31*)), the control of the adiponectin promoter (AdipoqCre(*32*)), the control of the villin promoter (Villin1Cre mice(*33*)) or the control of the rat Synapsin I promoter (*Syn1Cre* mice(*34*)). Breeding *Irs1loxP/loxP* AlfpCre mice with *Irs1loxP/loxP* mice resulted in hepatocyte specific *Irs1* deletion (AlfpCre::Irs1fl/fl denoted as lKO) and littermate control (Irsfl/fl) mice. Breeding *Irs1loxP/loxP* CkmmCre mice with *Irs1loxP/loxP* mice resulted in skeletal muscle specific *Irs1* deletion with partial deletion in cardiac muscle (CkmmCre::Irs1fl/fl denoted as mKO) and littermate control (Irsfl/fl) mice. Breeding *Irs1loxP/loxP* AdipoqCre mice with *Irs1loxP/loxP* mice resulted in fat specific *Irs1* deletion in both white and brown adipose tissue (AdipoqCre::Irs1fl/fl denoted as fKO) and littermate control (Irsfl/fl) mice. To avoid germline deletion in Syn1Cre in mice(*87*), only female *Irs1loxP/loxP* Syn1Cre mice were bred with male *Irs1loxP/loxP* mice to produce neuron specific *Irs1* deletion (Syn1Cre::Irsfl/fl denoted as nKO) and littermate control (Irsfl/fl) mice.

#### Mouse Husbandry

All mice were maintained in groups of four to five same-sex (only females were randomized) individuals under specific pathogen-free conditions in individually ventilated cages (Techniplast UK Ltd, Kettering, Northamptonshire, UK) to provide a controlled temperature and humidity environment with 12-h light/dark cycle (lights on from 06:00 - 18:00) and provided ad libitum access to food (ssniff® R/M-H phytoestrogen-poor (9% fat, 34% protein, 57% carbohydrate) ssniff Spezialdiäten GmbH, Soest, Germany) and sterile-filtered water. Sentinel mice in the animal room were regularly checked to be negative for mouse pathogens according to FELASA recommendations.

#### Mouse ethics

Mouse experiments were performed in accordance with the recommendations and guidelines of the Federation of the European Laboratory Animal Science Association (FELASA), with all protocols approved by the Landesamt fur Natur, Umwelt und Verbraucherschutz Nordrhein-Westfalen, Germany. Ethical permission requests were filed under 84-02.04.2014.A424 and 81-02.04.2019.A078.

#### Mouse genotyping

Mutant mice were identified by PCR genotyping using DNA extracted from ear clip biopsy and amplified using GoTaq® G2 DNA Polymerase, moreover tail clips were taken from mice at death for genotype confirmation. Primers used to genotype as well as expected size of amplicons, of Irs1 wild type, Irs1 knockout, Irs1 LoxP floxed allele, and different Cre lines are listed in Table S1.

#### Mouse tissue collection

Mice were euthanized by transcardial perfusion with PBS + EDTA (only for Irs1KO and nKO mice), after general anaesthesia with a cocktail of Ketamine (120 mg/kg) and Xylazine (10 mg/kg) with supplementary Isoflurane (5%) until no reflex response was observed. Then, blood was collected by cardiac puncture in tubes with EDTA, plasma-EDTA was isolated by centrifugation at 1,000g for at least 10 min at 4 °C before aliquoting and storage at −80 °C. Mice were rapidly decapitated, then the skull and body were dissected by two different scientists simultaneously to minimise tissue deterioration. The brain was removed from the skull and different brain regions were quickly isolated and snap-frozen in liquid nitrogen. The same cortical brain region was dissected for mitochondrial respirometry and prepared separately. The body of the animal was dissected and organs collected for histology in PFA or for molecular analysis were snap frozen in liquid nitrogen. The same procedure was performed for lKO, mKO and fKO with the exception of perfusion. Mice from lKO, mKO and fKO lines were sacrificed by cervical dislocation followed by rapid tissue removal as mentioned above.

### Mouse Metabolic Phenotyping

A longitudinal cohort of mice was assessed at young (Irs1KO = 6 months while lKO, mKO, fKO, and nKO = 4 months) and old (Irs1KO and lKO, mKO, fKO, and nKO = 16 months) age for general metabolic health outcomes.

#### Body composition

Body fat and lean mass content were measured in vivo by nuclear magnetic resonance using a minispec LF50H (Bruker Optics).

#### Collection of blood samples and determination of blood glucose levels

A small drop of blood was obtained from the tail of mice. Blood glucose levels were determined using an automatic glucose monitor (Accu-Check Aviva, Roche). Determination of blood glucose and collection of blood samples were always performed in the morning to avoid deviations due to circadian variations.

#### Insulin tolerance test

After determination of basal blood glucose levels, each animal received an intraperitoneal injection of insulin (0.75 U/kg body weight) (Sanofi). Blood glucose levels were measured 15, 30 and 60 min after insulin injection.

#### Glucose Tolerance test

Glucose tolerance tests were performed in the morning with animals after a 16 hour fast. After determination of fasted blood glucose levels, each animal received an intraperitoneal injection of 20% (w/v) glucose (10 ml/kg body weight). Blood glucose levels were measured 15, 30, 60 and 120 min after the glucose injection

#### Indirect calorimetry

Indirect calorimetry, locomotor activity, drinking and feeding were monitored for singly housed mice in purpose-built cages (Phenomaster, TSE Systems). Parameters such as food consumption, respiration, and locomotor activity were measured continuously for 48 hours after one day of acclimatisation and two days of training in similar cages. Values for locomotor activity were averaged for active and inactive phases separately for the 48 hour duration with the exception of the first and last hour of each phase. Metabolic rate was assessed by regression analysis using body weight as a covariate as recommended(*88*), except for Irs1KO mice due to the effect of the mutation on body size.

### Metabolomics

#### 2-phase metabolite extraction of polar and lipophilic metabolites of brain tissue

For the preparation of polar and lipophilic metabolites between 10 and 30 mg of snap-frozen mouse tissue (Irs1KO young = 5 months and old = 19 months, nKO young = 5 months and old = 22 months) was collected in 2 mL Eppendorf (www.eppendorf.com) tubes. For the extraction of the snap frozen material, the tissue was homogenised to a fine powder using a ball mill-type grinder (Tissuelyser II Qiagen, 85300). For the homogenization of the frozen material one liquid nitrogen cooled 5 mm stainless steel metal balls was added to each Eppendorf tube and the frozen material was disintegrated for 1 min at 25 Hz.

Metabolites were extracted by adding 1 mL of pre-cooled (−20°C) extraction buffer (methyl tert-butyl ether (MTBE): UPLC-grade methanol: UPLC-grade water 5:3:2 [v:v:v]), containing an equivalent 0.2 μL of EquiSplash Lipidomix (www.avantilipids.com) as an internal standard. The tubes were immediately vortexed until the sample was well re-suspended in the extraction buffer. The homogenised samples were incubated on a cooled (4°C) orbital mixer at 1500 rpm for 30 min. After this step, the metal ball was removed using a magnet and the samples were centrifuged for 10 min at 21,100 x g in a cooled table top centrifuge (4°C). The supernatant was transferred to a fresh 2 mL Eppendorf tube and 250 μL of MTBE and 150 μL of UPLC-grade water were added to each sample. The tubes were immediately vortexed before incubating them for an additional 10 min on a cooled (15°C) orbital mixer at 1500 rpm, before centrifuging them for 10 min at 15°C and 16,000 x g. After the centrifugation, the tubes contained two distinct phases. The upper MTBE phase contains the lipids, while the lower methanol-water phase contains the polar and semi-polar metabolites.

For the lipidomic analysis 600 μL of the upper lipid phase were collected into fresh 1.5 mL Eppendorf tubes, which were stored at -80°C, until mass spectrometric analysis. The remaining polar phase (∼800 μL) was immediately dried in a SpeedVac concentrator and stored dry at -80°C until mass spectrometric analysis.

#### Semi-targeted liquid chromatography-high-resolution mass spectrometry-based (LC-HRS-MS) analysis of amine-containing metabolites of brain tissue

The LC-HRMS analysis of amine-containing compounds was performed using an adapted benzoyl chloride-based derivatization method(*89*). In brief: The polar fraction of the metabolite extract was re-suspended in 200 μL of LC-MS-grade water (Optima-Grade, Thermo Fisher Scientific) and incubated at 4°C for 15 min on a thermomixer. The re-suspended extract was centrifuged for 5 min at 21,100 x g at 4°C and 50 μL of the cleared supernatant were mixed in an auto-sampler vial with a 200 μL glass insert (Chromatography Accessories Trott, Germany). The aqueous extract was mixed with 25 μl of 100 mM sodium carbonate (Sigma), followed by the addition of 25 μl 2% [v/v] benzoyl chloride (Sigma) in acetonitrile (Optima-Grade, Thermo Fisher Scientific). Samples were vortexed and kept at 20°C until analysis. For the LC-HRMS analysis, 1 μl of the derivatized sample was injected onto a 100 × 2.1 mm HSS T3 UPLC column (Waters). The flow rate was set to 400 μl/min using a binary buffer system consisting of buffer A (10 mM ammonium formate (Sigma), 0.15% [v/v] formic acid (Sigma) in LC-MS-grade water (Optima-Grade, Thermo Fisher Scientific). Buffer B consisted solely of acetonitrile (Optima-grade, Thermo Fisher-Scientific).

The mass spectrometer (Orbitrap ID-X, Thermo Fisher Scientific) was operated in positive ionisation mode recording the mass range m/z 100-1000. The heated ESI source settings of the mass spectrometer were: Spray voltage 3.5 kV, capillary temperature 300°C, sheath gas flow 60 AU, aux gas flow 20 AU at a temperature of 340°C and the sweep gas to 2 AU. The RF-lens was set to a value of 60%. Semi-targeted data analysis for the samples was performed using the TraceFinder software (Version 4.1, Thermo Fisher Scientific). The identity of each compound was validated by authentic reference compounds, which were run before and after every sequence. Peak areas of [M + nBz + H]^+^ ions were extracted using a mass accuracy (<5 ppm) and a retention time tolerance of <0.05 min. Areas of the cellular pool sizes were normalised to the internal standards (U-^15^N;U-^13^C amino acid mix (MSK-A2-1.2), Cambridge Isotope Laboratories), which were added to the extraction buffer, followed by a normalisation to the fresh weight of the analysed sample.

#### Anion-Exchange Chromatography Mass Spectrometry (AEX-MS) for the analysis of anionic metabolites of brain tissue

Extracted metabolites were re-suspended in 150-200 μl of Optima UPLC/MS grade water (Thermo Fisher Scientific). After 15 min incubation on a thermomixer at 4°C and a 5 min centrifugation at 21100 x g at 4°C, 100 μl of the cleared supernatant were transferred to polypropylene autosampler vials (Chromatography Accessories Trott, Germany) before AEX MS analysis. The samples were analysed using a Dionex ion chromatography system (Integrion, Thermo Fisher Scientific) as described previously(*90*). The detailed quantitative and qualitative transitions and electronic settings for the analysed metabolites are summarised in Table S2. For data analysis the area of the deprotonated [M-H+]-monoisotopic mass peak of each compound was extracted and integrated using a mass accuracy <5 ppm and a retention time (RT) tolerance of <0.1 min as compared to the independently measured reference compounds. Areas of the cellular pool sizes were normalised to the internal standards (citric acid D4), which were added to the extraction buffer, followed by a normalisation to the fresh weight of the analysed sample.

### Quantitative Real-time Polymerase Chain Reaction

RNA was extracted according to the manufacturer’s instructions using the TRIzol™ Reagent (ThermoFisher, 15596018) in Lysing Matrix D tubes (speed 6 for 40 seconds) (MP Biomedicals, 6913-500). RNA was precipitated with the aid of GlycoBlue Coprecipitant (ThermoFisher, AM9515) overnight at −80 °C. RNA was treated with DNase using the DNA-free™ kit (ThermoFisher, AM1906) according to manufacturer’s instructions. cDNA was prepared with the SuperScript® III First-Strand Synthesis SuperMix (ThermoFisher, 18080400) for q-RT-PCR. Samples of cDNA mixed with the PowerUp SYBR Green Master Mix (ThermoFisher, 4368706) and primers validated using a standard curve, were loaded in technical quadruplicates for q-RT-PCR on a QuantStudio™ 6 Flex Real-Time PCR System (ThermoFisher, 4485691). The ΔΔCT method was used to provide gene expression values after normalising to the known reference gene *B2M*. Primer sequences used for q-RT-PCR are shown in Table S3. Samples of cDNA mixed with the TaqMan™ Fast Advanced Master Mix (ThermoFisher, 4444557) and TaqMan® Assay probes, were loaded in technical quadruplicates for q-RT-PCR on a QuantStudio™ 6 Flex Real-Time PCR System (ThermoFisher, 4485691). The ΔΔCT method was used to provide gene expression values after normalising to the known reference gene *B2M*. Probe catalogue numbers used for q-RT-PCR are shown in Table S3.

### Mitochondrial respirometry

#### Mitochondrial mediums

Mediums were prepared as described in(*91*). Briefly, solution A contained 250 mM sucrose, 1 g/L BSA, 0.5 mM Na2EDTA, 10 mM Tris–HCl, pH 7.4. Solution B contained 20 mM taurine, 15 mM phosphocreatine, 20 mM imidazole, 0.5 mM DTT, 10 mM CaEGTA, 5.77 mM ATP, 6.56 mM MgCl2, 50 mM K-MES, pH 7.1. Solution C contained 0.5 mM EGTA, 60 mM K-lactobionate, 20 mM taurine, 10 mM KH2PO4, 3 mM MgCl2, 110 mM sucrose, 1 g/L fatty acid free BSA, 20 mM hepes, pH 7.1.

#### Tissue permeabilization

Protocol was adjusted from(*91*). In brief, cortical pieces (approximately 1 mm × 1 mm × 2 mm) were quickly removed and placed in cold solution A (250 mM sucrose, 1 g/L BSA, 0.5 mM Na2EDTA, 10 mM Tris–HCl, pH 7.4). Then, cortical tissue was weighed for normalisation of oxygen consumption and transferred into 2 mL tubes with 1 mL of cold solution B (20 mM taurine, 15 mM phosphocreatine, 20 mM imidazole, 0.5 mM DTT, 10 mM CaEGTA, 0.1 μM free Ca, 5.77 mM ATP, 6.56 mM MgCl2, 50 mM K-MES, pH 7.1). Medium was replaced by 2 mL of cold solution B complemented with 20 μL of a freshly prepared 5 mg/mL saponin solution. After 30 min at 4 °C under gentle agitation on an orbital shaker, samples were rinsed in cold solution C (0.5 mM EGTA, 60 mM K-lactobionate, 20 mM taurine, 10 mM KH2PO4, 3 mM MgCl2, 110 mM sucrose, 1 g/L fatty acid free BSA, 20 mM hepes, pH 7.1) three times for two minutes each, and further incubated on ice until measurements were taken.

#### Oxygen consumption measurement

Oxygen consumption of intact permeabilized cortical sections were measured using a respirometer (Oxygraph-2k, Oroboros Instruments). Measurements were performed under continuous stirring in 2 mL of solution C at 37 °C. The solution was equilibrated in air for at least 30 minutes, and permeabilized cortical tissue was transferred into the instrument chambers. Mutant mice with respective controls were run in parallel in the instrument’s two chambers simultaneously to minimise and day to day variability. Only after stabilisation of the initial mitochondrial oxygen consumption, mitochondrial respiration was stimulated by successive addition of substrates and inhibitors: First, 5 ul of 2M pyruvate and 5 ul of 800 mM Malate were added. Second, 10 ul of 500 mM ADP (with 300 mM free Mg2+) was added to measure O2 consumption under normal phosphorylating state. Third, 5 ul of 4 mM Cytochrome C was added to check for mitochondrial membrane integrity, any samples that responded with a significant increase in O2 consumption were removed from further analysis. Fourth, 10 ul of 2 M glutamate was added. Fifth, 20 ul of 1 M succinate was added. Sixth, 1 ul of 5 mM oligomycin, an ATP synthase inhibitor, was added. Then, maximum mitochondrial capacity was assessed by adding gradual volumes of 1 mM FCCP until maximum oxygen consumption was reached. Then, 1 ul of 1 mM rotenone, a complex one inhibitor, was added to measure complex two activity. Finally, 1 ul of 5 mM antimycin A, a complex three inhibitor, was added to measure non-mitochondrial O2 consumption (residual oxygen flux) due to cytosolic oxidases. Residual oxygen flux was subtracted from all other measurements to report baseline mitochondrial oxygen consumption. Mitochondrial oxygen consumption was calculated using DataGraph software from the manufacturer (Oroboros Instruments).

### Statistical analyses

Mean lifespan was assessed, and survivorship was analysed using log rank test or Cox proportional hazard analysis where denoted. Two-group comparisons were made using two-tailed, unpaired Student’s t-test unless otherwise stated. For comparisons of multiple factors (for example, phase * genotype or sex*genotype), two-way ANOVA was reported, followed by Sidak’s post-test if interaction between main effects was significant.

Numbers of mice were estimated to be sufficient to detect statistically meaningful differences of at least 20% between or among groups using standard power calculations with α = 0.05 and power of 0.8 on the basis of similar experiments conducted in our group. Homogeneity of variance and normality of residuals were assessed, and appropriate corrections were made if necessary. All experiments were performed in a randomised and blinded fashion when possible. Data were analysed statistically using GraphPad Prism 9.4, outliers were removed from analysis based on a Grubb’s test. The value of α was 0.05, and data were expressed as *P < 0.05; **P < 0.01; ***P < 0.001; ****P < 0.0001. Number of animals reported at the bottom of the bars for each condition or in figure legends. All error bars correspond to standard deviation except for longitudinal glucose and insulin sensitivity where standard error of the mean is reported. ANCOVA analyses were plotted with 95% confidence interval bands. Detailed P values for non-significant comparisons, test statistic values, and degrees of freedom included in Table S1.

## Supporting information

All supplementary material

## Acknowledgments

We thank the metabolomics, comparative and phenotyping facility of the Max Planck Institute for the Biology of Ageing for outstanding technical and analytical help and advice, Aleksandra Trifunovic for critical discussions, and Martin Purrio for technical assistance with mouse phenotyping.

## Author contributions

Conceptualization: MB, LP

Resources: MB, TN

Formal analysis: MB

Supervision: SG, LP

Funding acquisition: MB, LP

Investigation: MB

Visualisation: MB

Methodology: MB, AM

Writing: MB, SG, LP

## Data and materials availability

All data needed to evaluate the conclusions in the paper are present in the paper and/or the Supplementary Materials. The mouse lines or collected organs can be provided by L.P. pending scientific review and a completed material transfer agreement.

## Supplementary Materials

Figs. S1 to S15

Tables S1 to S3

## References

1. L. Partridge, J. Deelen, P. E. Slagboom, Facing up to the global challenges of ageing. Nature. 561, 45–56 (2018).

2. S. N. Austad, K. E. Fischer, Sex Differences in Lifespan. Cell Metab. 23 (2016), doi:10.1016/j.cmet.2016.05.019.

3. L. Fontana, L. Partridge, Promoting health and longevity through diet: from model organisms to humans. Cell. 161 (2015), doi:10.1016/j.cell.2015.02.020.

4. C. J. Kenyon, The genetics of ageing. Nature. 464, 504–512 (2010).

5. M. D. W. Piper, L. Partridge, Drosophila as a model for ageing. Biochim. Biophys. Acta Mol. Basis Dis. 1864 (2018), doi:10.1016/j.bbadis.2017.09.016.

6. J. Yuyang Lu, A. Seluanov, V. Gorbunova, Long-lived fish in a big pond. Science. 374, pp. 824–825.

7. C. Selman, L. Partridge, D. J. Withers, Replication of extended lifespan phenotype in mice with deletion of insulin receptor substrate 1. PLoS One. 6 (2011), doi:10.1371/journal.pone.0016144.

8. A. Nojima, M. Yamashita, Y. Yoshida, I. Shimizu, H. Ichimiya, N. Kamimura, Y. Kobayashi, S. Ohta, N. Ishii, T. Minamino, Haploinsufficiency of akt1 prolongs the lifespan of mice. PLoS One. 8 (2013), doi:10.1371/journal.pone.0069178.

9. C. Selman, J. M. Tullet, D. Wieser, E. Irvine, S. J. Lingard, A. I. Choudhury, M. Claret, H. Al-Qassab, D. Carmignac, F. Ramadani, A. Woods, I. C. Robinson, E. Schuster, R. L. Batterham, S. C. Kozma, G. Thomas, D. Carling, K. Okkenhaug, J. M. Thornton, L. Partridge, D. Gems, D. J. Withers, Ribosomal protein S6 kinase 1 signaling regulates mammalian life span. Science. 326 (2009), doi:10.1126/science.1177221.

10. J. J. Wu, J. Liu, E. B. Chen, J. J. Wang, L. Cao, N. Narayan, M. M. Fergusson, I. I. Rovira, M. Allen, D. A. Springer, C. U. Lago, S. Zhang, W. DuBois, T. Ward, R. deCabo, O. Gavrilova, B. Mock, T. Finkel, Increased mammalian lifespan and a segmental and tissue-specific slowing of aging after genetic reduction of mTOR expression. Cell Rep. 4 (2013), doi:10.1016/j.celrep.2013.07.030.

11. J. Deelen, D. S. Evans, D. E. Arking, N. Tesi, M. Nygaard, X. Liu, M. K. Wojczynski, M. L. Biggs, A. van der Spek, G. Atzmon, E. B. Ware, C. Sarnowski, A. V. Smith, I. Seppälä, H. J. Cordell, J. Dose, N. Amin, A. M. Arnold, K. L. Ayers, N. Barzilai, E. J. Becker, M. Beekman, H. Blanché, K. Christensen, L. Christiansen, J. C. Collerton, S. Cubaynes, S. R. Cummings, K. Davies, B. Debrabant, J.-F. Deleuze, R. Duncan, J. D. Faul, C. Franceschi, P. Galan, V. Gudnason, T. B. Harris, M. Huisman, M. A. Hurme, C. Jagger, I. Jansen, M. Jylhä, M. Kähönen, D. Karasik, S. L. R. Kardia, A. Kingston, T. B. L. Kirkwood, L. J. Launer, T. Lehtimäki, W. Lieb, L.-P. Lyytikäinen, C. Martin-Ruiz, J. Min, A. Nebel, A. B. Newman, C. Nie, E. A. Nohr, E. S. Orwoll, T. T. Perls, M. A. Province, B. M. Psaty, O. T. Raitakari, M. J. T. Reinders, J.-M. Robine, J. I. Rotter, P. Sebastiani, J. Smith, T. I. A. Sørensen, K. D. Taylor, A. G. Uitterlinden, W. van der Flier, S. J. van der Lee, C. M. van Duijn, D. van Heemst, J. W. Vaupel, D. Weir, K. Ye, Y. Zeng, W. Zheng, H. Holstege, D. P. Kiel, K. L. Lunetta, P. E. Slagboom, J. M. Murabito, A meta-analysis of genome-wide association studies identifies multiple longevity genes. Nat. Commun. 10, 1–14 (2019).

12. J.-R. Lin, P. Sin-Chan, V. Napolioni, G. G. Torres, J. Mitra, Q. Zhang, M. R. Jabalameli, Z. Wang, N. Nguyen, T. Gao, M. Laudes, S. Görg, A. Franke, A. Nebel, M. D. Greicius, G. Atzmon, K. Ye, V. Gorbunova, W. C. Ladiges, A. R. Shuldiner, L. J. Niedernhofer, P. D. Robbins, S. Milman, Y. Suh, J. Vijg, N. Barzilai, Z. D. Zhang, Rare genetic coding variants associated with human longevity and protection against age-related diseases. Nature Aging. 1, 783–794 (2021).

13. C. Tazearslan, J. Huang, N. Barzilai, Y. Suh, Impaired IGF1R signaling in cells expressing longevity-associated human IGF1R alleles. Aging Cell. 10 (2011), doi:10.1111/j.1474-9726.2011.00697.x.

14. M. Björnholm, A. R. He, A. Attersand, S. Lake, S. C. Liu, G. E. Lienhard, S. Taylor, P. Arner, J. R. Zierath, Absence of functional insulin receptor substrate-3 (IRS-3) gene in humans. Diabetologia. 45 (2002), doi:10.1007/s00125-002-0945-z.

15. C. Selman, S. Lingard, A. I. Choudhury, R. L. Batterham, M. Claret, M. Clements, F. Ramadani, K. Okkenhaug, E. Schuster, E. Blanc Piper, H. Al-Qassab, J. R. Speakman, D. Carmignac, I. C. Robinson, J. M. Thornton, D. Gems, L. Partridge, D. J. Withers, Evidence for lifespan extension and delayed age-related biomarkers in insulin receptor substrate 1 null mice. FASEB J. 22 (2008), doi:10.1096/fj.07-9261com.

16. M. Rincon, R. Muzumdar, G. Atzmon, N. Barzilai, The paradox of the insulin/IGF-1 signaling pathway in longevity. Mech. Ageing Dev. 125 (2004), doi:10.1016/j.mad.2004.03.006.

17. Z. Cheng, S. Guo, K. Copps, X. Dong, R. Kollipara, J. T. Rodgers, R. A. Depinho, P. Puigserver, M. F. White, Foxo1 integrates insulin signaling with mitochondrial function in the liver. Nat. Med. 15 (2009), doi:10.1038/nm.2049.

18. J. Nunnari, A. Suomalainen, Mitochondria: in sickness and in health. Cell. 148 (2012), doi:10.1016/j.cell.2012.02.035.

19. M. T. Lin, M. Flint Beal, Mitochondrial dysfunction and oxidative stress in neurodegenerative diseases. Nature. 443 (2006), pp. 787–795.

20. P. M. Quirós, M. A. Prado, N. Zamboni, D. D’Amico, R. W. Williams, D. Finley, S. P. Gygi, J. Auwerx, Multi-omics analysis identifies ATF4 as a key regulator of the mitochondrial stress response in mammals. J. Cell Biol. 216 (2017), doi:10.1083/jcb.201702058.

21. W. Li, X. Li, R. A. Miller, ATF4 activity: a common feature shared by many kinds of slow-aging mice. Aging Cell. 13 (2014), doi:10.1111/acel.12264.

22. D. Zhou, L. R. Palam, L. Jiang, J. Narasimhan, K. A. Staschke, R. C. Wek, Phosphorylation of eIF2 directs ATF5 translational control in response to diverse stress conditions. J. Biol. Chem. 283 (2008), doi:10.1074/jbc.M708530200.

23. C. J. Fiorese, A. M. Schulz, Y. F. Lin, N. Rosin, M. W. Pellegrino, C. M. Haynes, The Transcription Factor ATF5 Mediates a Mammalian Mitochondrial UPR. Curr. Biol. 26 (2016), doi:10.1016/j.cub.2016.06.002.

24. Y. Zhang, Y. Xie, E. D. Berglund, K. C. Coate, T. T. He, T. Katafuchi, G. Xiao, M. J. Potthoff, W. Wei, Y. Wan, R. T. Yu, R. M. Evans, S. A. Kliewer, D. J. Mangelsdorf, The starvation hormone, fibroblast growth factor-21, extends lifespan in mice. Elife. 1 (2012), doi:10.7554/eLife.00065.

25. J. R. Speakman, Body size, energy metabolism and lifespan. J. Exp. Biol. 208 (2005), doi:10.1242/jeb.01556.

26. S. Yanai, S. Endo, Functional Aging in Male C57BL/6J Mice Across the Life-Span: A Systematic Behavioral Analysis of Motor, Emotional, and Memory Function to Define an Aging Phenotype. Front. Aging Neurosci. 13, 697621 (2021).

27. M. F. White, C. Ronald Kahn, Insulin action at a molecular level – 100 years of progress. Molecular Metabolism. 52 (2021), doi:10.1016/j.molmet.2021.101304.

28. M. P. Czech, Insulin action and resistance in obesity and type 2 diabetes. Nat. Med. 23 (2017), doi:10.1038/nm.4350.

29. P. Essers, L. S. Tain, T. Nespital, J. Goncalves, J. Froehlich, L. Partridge, Reduced insulin/insulin-like growth factor signaling decreases translation in Drosophila and mice. Sci. Rep. 6, 30290 (2016).

30. C. Kellendonk, C. Opherk, K. Anlag, G. Schütz, F. Tronche, Hepatocyte-specific expression of Cre recombinase. Genesis. 26 (2000), doi:10.1002/(sici)1526-968x(200002)26:2<151::aid-gene17>3.0.co;2-e.

31. J. C. Brüning Michael, J. N. Winnay, T. Hayashi, D. Hörsch, D. Accili, L. J. Goodyear, C. R. Kahn, A muscle-specific insulin receptor knockout exhibits features of the metabolic syndrome of NIDDM without altering glucose tolerance. Mol. Cell. 2 (1998), doi:10.1016/s1097-2765(00)80155-0.

32. J. Eguchi, X. Wang, S. Yu, E. E. Kershaw, P. C. Chiu, J. Dushay, J. L. Estall, U. Klein, E. Maratos-Flier, E. D. Rosen, Transcriptional control of adipose lipid handling by IRF4. Cell Metab. 13, 249 (2011).

33. B. B. Madison, L. Dunbar, X. T. Qiao, K. Braunstein, E. Braunstein, D. L. Gumucio, Cis elements of the villin gene control expression in restricted domains of the vertical (crypt) and horizontal (duodenum, cecum) axes of the intestine. J. Biol. Chem. 277 (2002), doi:10.1074/jbc.M204935200.

34. Y. Zhu, M. I. Romero, P. Ghosh, Z. Ye, P. Charnay, E. J. Rushing, J. D. Marth, L. F. Parada, Ablation of NF1 function in neurons induces abnormal development of cerebral cortex and reactive gliosis in the brain. Genes Dev. 15, 859 (2001).

35. S. Della Torre, N. Mitro, C. Meda, F. Lolli, S. Pedretti, M. Barcella, L. Ottobrini, D. Metzger, D. Caruso, A. Maggi, Short-Term Fasting Reveals Amino Acid Metabolism as a Major Sex-Discriminating Factor in the Liver. Cell Metab. 28, 256 (2018).

36. S. F. Previs, D. J. Withers, J. M. Ren, M. F. White, G. I. Shulman, Contrasting effects of IRS-1 versus IRS-2 gene disruption on carbohydrate and lipid metabolism in vivo. J. Biol. Chem. 275 (2000), doi:10.1074/jbc.M006490200.

37. Y. C. Long, Z. Cheng, K. D. Copps, M. F. White, Insulin receptor substrates Irs1 and Irs2 coordinate skeletal muscle growth and metabolism via the Akt and AMPK pathways. Mol. Cell. Biol. 31 (2011), doi:10.1128/MCB.00983-10.

38. M. Blüher Michael, O. D. Peroni, K. Ueki, N. Carter, B. B. Kahn, C. R. Kahn, Adipose tissue selective insulin receptor knockout protects against obesity and obesity-related glucose intolerance. Dev. Cell. 3 (2002), doi:10.1016/s1534-5807(02)00199-5.

39. M. Blüher, B. B. Kahn, C. Ronald Kahn, Extended Longevity in Mice Lacking the Insulin Receptor in Adipose Tissue. Science (2003), doi:10.1126/science.1078223.

40. A Muscle-Specific Insulin Receptor Knockout Exhibits Features of the Metabolic Syndrome of NIDDM without Altering Glucose Tolerance. Mol. Cell. 2, 559–569 (1998).

41. J. Boucher, S. Softic, A. El Ouaamari, M. T. Krumpoch, A. Kleinridders, R. N. Kulkarni, B. T. O’Neill, C. R. Kahn, Differential Roles of Insulin and IGF-1 Receptors in Adipose Tissue Development and Function. Diabetes. 65 (2016), doi:10.2337/db16-0212.

42. B. T. O’Neill, H. P. Lauritzen, M. F. Hirshman, G. Smyth, L. J. Goodyear, C. R. Kahn, Differential Role of Insulin/IGF-1 Receptor Signaling in Muscle Growth and Glucose Homeostasis. Cell Rep. 11 (2015), doi:10.1016/j.celrep.2015.04.037.

43. X. C. Dong, K. D. Copps, S. Guo, Y. Li, R. Kollipara, R. A. DePinho, M. F. White, Inactivation of hepatic Foxo1 by insulin signaling is required for adaptive nutrient homeostasis and endocrine growth regulation. Cell Metab. 8 (2008), doi:10.1016/j.cmet.2008.06.006.

44. N. Kubota, T. Kubota, S. Itoh, H. Kumagai, H. Kozono, I. Takamoto, T. Mineyama, H. Ogata, K. Tokuyama, M. Ohsugi, T. Sasako, M. Moroi, K. Sugi, S. Kakuta, Y. Iwakura, T. Noda, S. Ohnishi, R. Nagai, K. Tobe, Y. Terauchi, K. Ueki, T. Kadowaki, Dynamic functional relay between insulin receptor substrate 1 and 2 in hepatic insulin signaling during fasting and feeding. Cell Metab. 8 (2008), doi:10.1016/j.cmet.2008.05.007.

45. J. Ruud, S. M. Steculorum, J. C. Brüning, Neuronal control of peripheral insulin sensitivity and glucose metabolism. Nat. Commun. 8, 1–12 (2017).

46. C. M. Hill, T. Laeger, M. Dehner, D. C. Albarado, B. Clarke, D. Wanders, S. J. Burke, J. J. Collier, E. Qualls-Creekmore, S. M. Solon-Biet, S. J. Simpson, H.-R. Berthoud, H. Münzberg, C. D. Morrison, FGF21 Signals Protein Status to the Brain and Adaptively Regulates Food Choice and Metabolism. Cell Rep. 27, 2934–2947.e3 (2019).

47. E. Harno, E. C. Cottrell, A. White, Metabolic pitfalls of CNS Cre-based technology. Cell Metab. 18 (2013), doi:10.1016/j.cmet.2013.05.019.

48. A. C. Könner, S. Hess, S. Tovar, A. Mesaros, C. Sánchez-Lasheras, N. Evers, L. A. Verhagen, H. S. Brönneke, A. Kleinridders, B. Hampel, P. Kloppenburg, J. C. Brüning, Role for insulin signaling in catecholaminergic neurons in control of energy homeostasis. Cell Metab. 13 (2011), doi:10.1016/j.cmet.2011.03.021.

49. Y. Zhang, I. A. Kerman, A. Laque, P. Nguyen, M. Faouzi, G. W. Louis, J. C. Jones, C. Rhodes, H. Münzberg, Leptin-receptor-expressing neurons in the dorsomedial hypothalamus and median preoptic area regulate sympathetic brown adipose tissue circuits. J. Neurosci. 31 (2011), doi:10.1523/JNEUROSCI.3223-10.2011.

50. K. Rezai-Zadeh, S. Yu, Y. Jiang, A. Laque, C. Schwartzenburg, C. D. Morrison, A. V. Derbenev, A. Zsombok, H. Münzberg, Leptin receptor neurons in the dorsomedial hypothalamus are key regulators of energy expenditure and body weight, but not food intake. Molecular metabolism. 3 (2014), doi:10.1016/j.molmet.2014.07.008.

51. A. C. Hausen, J. Ruud, H. Jiang, S. Hess, H. Varbanov, P. Kloppenburg, J. C. Brüning, Insulin-Dependent Activation of MCH Neurons Impairs Locomotor Activity and Insulin Sensitivity in Obesity. Cell Rep. 17 (2016), doi:10.1016/j.celrep.2016.11.030.

52. L. S. Tain, R. Sehlke, C. Jain, M. Chokkalingam, N. Nagaraj, P. Essers, M. Rassner, S. Grönke, J. Froelich, C. Dieterich, M. Mann, N. Alic, A. Beyer, L. Partridge, A proteomic atlas of insulin signalling reveals tissue-specific mechanisms of longevity assurance. Mol. Syst. Biol. 13, 939 (2017).

53. N. Yadava, D. G. Nicholls, Spare respiratory capacity rather than oxidative stress regulates glutamate excitotoxicity after partial respiratory inhibition of mitochondrial complex I with rotenone. J. Neurosci. 27 (2007), doi:10.1523/JNEUROSCI.0212-07.2007.

54. L. I. Johnson-Cadwell, M. B. Jekabsons, A. Wang, B. M. Polster, D. G. Nicholls, “Mild Uncoupling” does not decrease mitochondrial superoxide levels in cultured cerebellar granule neurons but decreases spare respiratory capacity and increases toxicity to glutamate and oxidative stress. J. Neurochem. 101 (2007), doi:10.1111/j.1471-4159.2007.04516.x.

55. S. W. Choi, A. A. Gerencser, D. G. Nicholls, Bioenergetic analysis of isolated cerebrocortical nerve terminals on a microgram scale: spare respiratory capacity and stochastic mitochondrial failure. J. Neurochem. 109 (2009), doi:10.1111/j.1471-4159.2009.06055.x.

56. P. Marchetti, Q. Fovez, N. Germain, R. Khamari, J. Kluza, Mitochondrial spare respiratory capacity: Mechanisms, regulation, and significance in non-transformed and cancer cells. FASEB J. 34 (2020), doi:10.1096/fj.202000767R.

57. H. P. Harding, Y. Zhang, H. Zeng, I. Novoa, P. D. Lu, M. Calfon, N. Sadri, C. Yun, B. Popko, R. Paules, D. F. Stojdl, J. C. Bell, T. Hettmann, J. M. Leiden, D. Ron, An integrated stress response regulates amino acid metabolism and resistance to oxidative stress. Mol. Cell. 11 (2003), doi:10.1016/s1097-2765(03)00105-9.

58. S. K. Young, R. C. Wek, Upstream Open Reading Frames Differentially Regulate Gene-specific Translation in the Integrated Stress Response. J. Biol. Chem. 291 (2016), doi:10.1074/jbc.R116.733899.

59. F. Amar, C. Corona, J. Husson, J. Liu, M. Shelanski, L. Greene, Rapid ATF4 Depletion Resets Synaptic Responsiveness after cLTP. eNeuro. 8 (2021), doi:10.1523/ENEURO.0239-20.2021.

60. I. Kühl, M. Miranda, I. Atanassov, I. Kuznetsova, Y. Hinze, A. Mourier, A. Filipovska, N.-G. Larsson, Transcriptomic and proteomic landscape of mitochondrial dysfunction reveals secondary coenzyme Q deficiency in mammals (2017), doi:10.7554/eLife.30952.

61. N. A. Khan, J. Nikkanen, S. Yatsuga, C. Jackson, L. Wang, S. Pradhan, R. Kivelä, A. Pessia, V. Velagapudi, A. Suomalainen, mTORC1 Regulates Mitochondrial Integrated Stress Response and Mitochondrial Myopathy Progression. Cell Metab. 26 (2017), doi:10.1016/j.cmet.2017.07.007.

62. J. Nikkanen, S. Forsström, L. Euro, I. Paetau, R. A. Kohnz, L. Wang, D. Chilov, J. Viinamäki, A. Roivainen, P. Marjamäki, H. Liljenbäck, S. Ahola, J. Buzkova, M. Terzioglu, N. A. Khan, S. Pirnes-Karhu, A. Paetau, T. Lönnqvist, A. Sajantila, P. Isohanni, H. Tyynismaa, D. K. Nomura, B. J. Battersby, V. Velagapudi, C. J. Carroll, A. Suomalainen, Mitochondrial DNA Replication Defects Disturb Cellular dNTP Pools and Remodel One-Carbon Metabolism. Cell Metab. 23, 635–648 (2016).

63. V. Byles, Y. Cormerais, K. Kalafut, V. Barrera, H. H. Je, S. H. Sui, J. M. Asara, C. M. Adams, G. Hoxhaj, I. Ben-Sahra, B. D. Manning, Hepatic mTORC1 signaling activates ATF4 as part of its metabolic response to feeding and insulin. Molecular metabolism. 53 (2021), doi:10.1016/j.molmet.2021.101309.

64. E. Motori, I. Atanassov, S. M. V. Kochan, K. Folz-Donahue, V. Sakthivelu, P. Giavalisco, N. Toni, J. Puyal, N. G. Larsson, Neuronal metabolic rewiring promotes resilience to neurodegeneration caused by mitochondrial dysfunction. Science advances. 6 (2020), doi:10.1126/sciadv.aba8271.

65. J. Durieux, S. Wolff, A. Dillin, The cell-non-autonomous nature of electron transport chain-mediated longevity. Cell. 144 (2011), doi:10.1016/j.cell.2010.12.016.

66. Fibroblast Growth Factor 21 Drives Dynamics of Local and Systemic Stress Responses in Mitochondrial Myopathy with mtDNA Deletions. Cell Metab. 30, 1040–1054.e7 (2019).

67. in Handbook of Models for Human Aging (Academic Press, 2006; http://dx.doi.org/10.1016/B978-012369391-4/50034-5), pp. 393–401.

68. L. Mulvey, A. Sinclair, C. Selman, Lifespan modulation in mice and the confounding effects of genetic background. J. Genet. Genomics. 41, 497–503 (2014).

69. M. Holzenberger, J. Dupont, B. Ducos, P. Leneuve, A. Géloën, P. C. Even, P. Cervera, Y. Le Bouc, IGF-1 receptor regulates lifespan and resistance to oxidative stress in mice. Nature. 421, 182–187 (2002).

70. A. F. Bokov, N. Garg, Y. Ikeno, S. Thakur, N. Musi, R. A. DeFronzo, N. Zhang, R. C. Erickson, J. Gelfond, G. B. Hubbard, M. L. Adamo, A. Richardson, Does Reduced IGF-1R Signaling in Igf1r +/-Mice Alter Aging? PLoS One. 6 (2011), doi:10.1371/journal.pone.0026891.

71. J. P. Liu, J. Baker, A. S. Perkins, E. J. Robertson, A. Efstratiadis, Mice carrying null mutations of the genes encoding insulin-like growth factor I (Igf-1) and type 1 IGF receptor (Igf1r). Cell. 75 (1993) (available at https://pubmed.ncbi.nlm.nih.gov/8402901/).

72. D. Accili, J. Drago, E. J. Lee Johnson, M. H. Cool, P. Salvatore, L. D. Asico, P. A. José, S. I. Taylor, H. Westphal, Early neonatal death in mice homozygous for a null allele of the insulin receptor gene. Nat. Genet. 12 (1996), doi:10.1038/ng0196-106.

73. A. Taguchi, L. M. Wartschow, M. F. White, Brain IRS2 signaling coordinates life span and nutrient homeostasis. Science. 317 (2007), doi:10.1126/science.1142179.

74. K. Martens, A. Bottelbergs, M. Baes, Ectopic recombination in the central and peripheral nervous system by aP2/FABP4-Cre mice: implications for metabolism research. FEBS Lett. 584 (2010), doi:10.1016/j.febslet.2010.01.061.

75. T. L. Merry, D. Kuhlow, B. Laube, D. Pöhlmann, A. F. H. Pfeiffer, C. R. Kahn, M. Ristow, K. Zarse, Impairment of insulin signalling in peripheral tissue fails to extend murine lifespan. Aging Cell. 16 (2017), doi:10.1111/acel.12610.

76. E. Cohen, J. F. Paulsson, P. Blinder, T. Burstyn-Cohen, D. Du, G. Estepa, A. Adame, H. M. Pham, M. Holzenberger, J. W. Kelly, E. Masliah, A. Dillin, Reduced IGF-1 signaling delays age-associated proteotoxicity in mice. Cell. 139 (2009), doi:10.1016/j.cell.2009.11.014.

77. S. M. Steculorum, M. Solas, J. C. Brüning, The paradox of neuronal insulin action and resistance in the development of aging-associated diseases. Alzheimers. Dement. 10, S3–11 (2014).

78. C. A. Wolkow, K. D. Kimura, M. S. Lee, G. Ruvkun, Regulation of C. elegans life-span by insulinlike signaling in the nervous system. Science. 290 (2000), doi:10.1126/science.290.5489.147.

79. M. Z. B. Ismail, M. D. Hodges, M. Boylan, R. Achall, A. Shirras, S. J. Broughton, The Drosophila Insulin Receptor Independently Modulates Lifespan and Locomotor Senescence. PLoS One. 10, e0125312 (2015).

80. H. Augustin, K. McGourty, M. J. Allen, S. K. Madem, J. Adcott, F. Kerr, C. T. Wong, A. Vincent, T. Godenschwege, E. Boucrot, L. Partridge, Reduced insulin signaling maintains electrical transmission in a neural circuit in aging flies. PLoS Biol. 15, e2001655 (2017).

81. E. L. Baar, K. A. Carbajal, I. M. Ong, D. W. Lamming, Sex- and tissue-specific changes in mTOR signaling with age in C57BL/6J mice. Aging Cell. 15, 155 (2016).

82. L. M. Restelli, B. Oettinghaus, M. Halliday, C. Agca, M. Licci, L. Sironi, C. Savoia, J. Hench, M. Tolnay, A. Neutzner, A. Schmidt, A. Eckert, G. Mallucci, L. Scorrano, S. Frank, Neuronal Mitochondrial Dysfunction Activates the Integrated Stress Response to Induce Fibroblast Growth Factor 21. Cell Rep. 24 (2018), doi:10.1016/j.celrep.2018.07.023.

83. H. Tyynismaa, C. J. Carroll, N. Raimundo, S. Ahola-Erkkilä, T. Wenz, H. Ruhanen, K. Guse, A. Hemminki, K. E. Peltola-Mjøsund, V. Tulkki, M. Orešič, C. T. Moraes, K. Pietiläinen, I. Hovatta, A. Suomalainen, Mitochondrial myopathy induces a starvation-like response. um. Mol. Genet. 19, 3948–3958 (2010).

84. N. E. Richardson, E. N. Konon, H. S. Schuster, A. T. Mitchell, C. Boyle, A. C. Rodgers, M. Finke, L. R. Haider, D. Yu, V. Flores, H. H. Pak, S. Ahmad, S. Ahmed, A. Radcliff, J. Wu, E. M. Williams, L. Abdi, D. S. Sherman, T. A. Hacker, D. W. Lamming, Lifelong restriction of dietary branched-chain amino acids has sex-specific benefits for frailty and life span in mice. Nature Aging. 1, 73–86 (2021).

85. D. Yu, S. E. Yang, B. R. Miller, J. A. Wisinski, D. S. Sherman, J. A. Brinkman, J. L. Tomasiewicz, N. E. Cummings, M. E. Kimple, V. L. Cryns, D. W. Lamming, Short-term methionine deprivation improves metabolic health via sexually dimorphic, mTORC1-independent mechanisms. The FASEB Journal. 32, 3471 (2018).

86. D. A. Fontaine, D. B. Davis, Attention to Background Strain Is Essential for Metabolic Research: C57BL/6 and the International Knockout Mouse Consortium. Diabetes. 65, 25–33 (2016).

87. D. Rempe, G. Vangeison, J. Hamilton, Y. Li, M. Jepson, H. J. Federoff, Synapsin I Cre transgene expression in male mice produces germline recombination in progeny. Genesis. 44 (2006), doi:10.1002/gene.20183.

88. T. D. Müller, M. Klingenspor, M. H. Tschöp, Revisiting energy expenditure: how to correct mouse metabolic rate for body mass. Nature Metabolism. 3, 1134–1136 (2021).

89. J. M. Wong, P. A. Malec, O. S. Mabrouk, J. Ro, M. Dus, R. T. Kennedy, Benzoyl chloride derivatization with liquid chromatography-mass spectrometry for targeted metabolomics of neurochemicals in biological samples. J. Chromatogr. A. 1446 (2016), doi:10.1016/j.chroma.2016.04.006.

90. M. Schwaiger, E. Rampler, G. Hermann, W. Miklos, W. Berger, G. Koellensperger, Anion-Exchange Chromatography Coupled to High-Resolution Mass Spectrometry: A Powerful Tool for Merging Targeted and Non-targeted Metabolomics. Anal. chem. 89 (2017), doi:10.1021/acs.analchem.7b01624.

91. A. Benani, V. Barquissau, L. Carneiro, B. Salin, A.-L. Colombani, C. Leloup, L. Casteilla, M. Rigoulet, L. Pénicaud, Method for functional study of mitochondria in rat hypothalamus. J. Neurosci. Methods. 178, 301–307 (2009).

