## Supplementary material for "Reduced insulin signalling in neurons induces sex-specific health benefits": All supplementary material

**This PDF file includes:**

Supplementary Text
Figs. S1 to S15
Tables S1 to S3

*Supplementary Figure 1*

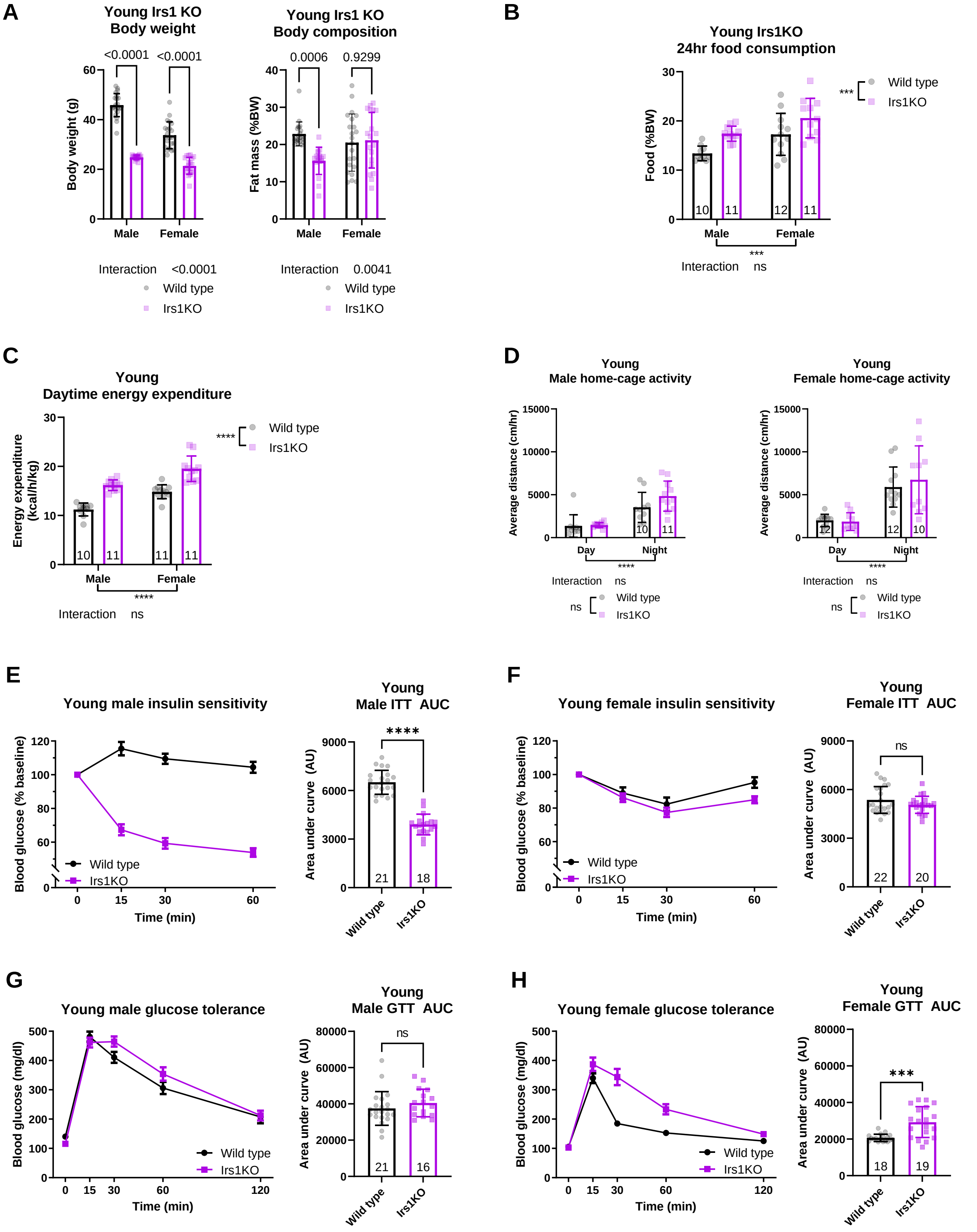


### Supplementary Figure 1: Characterisation of young Irs1KO

**(A)** Body weight and body composition of Irs1KO mice measured at young age (5 months) (Wild type males n=21, Irs1KO males n=18, Wild type females n=22, Irs1KO females n=19). **(B)** Measurement of food consumption of young Irs1KO and wild type mice revealed a significant increase in food consumption of single housed Irs1KO male and female mice. **(C)** Young body weight normalised energy expenditure of singly housed male and female mice during daytime as assessed by metabolic chambers showing an increase in energy expenditure in Irs1KO mice. **(D)** Spontaneous activity of Irs1KO single housed mice during their inactive cycle or daytime showed no significant difference in activity at young age (Wild type males n=10, Irs1KO males n=11, Wild type females n=12, Irs1KO females n=10). Insulin tolerance test (ITT) of young male **(E)** Irs1KO mice revealed a clear enhanced systemic insulin sensitivity in male Irs1KO mice, while no difference was observed between female **(F)** Irs1KO mice and their wild type littermates. **(G)** Glucose tolerance test (GTT) did not show any significant difference in glucose tolerance in young male Irs1KO and wild type littermate mice. **(H)** AUC analysis of GTT in young female Irs1KO mice revealed a significantly reduced glucose tolerance in Irs1KO females compared to wild type littermate mice. All error bars correspond to standard deviation except for longitudinal glucose and insulin sensitivity where standard error of the mean is reported. Number of animals reported at the bottom of the bars for each condition. Detailed statistical values found in Table S1.

*Supplementary Figure 2*

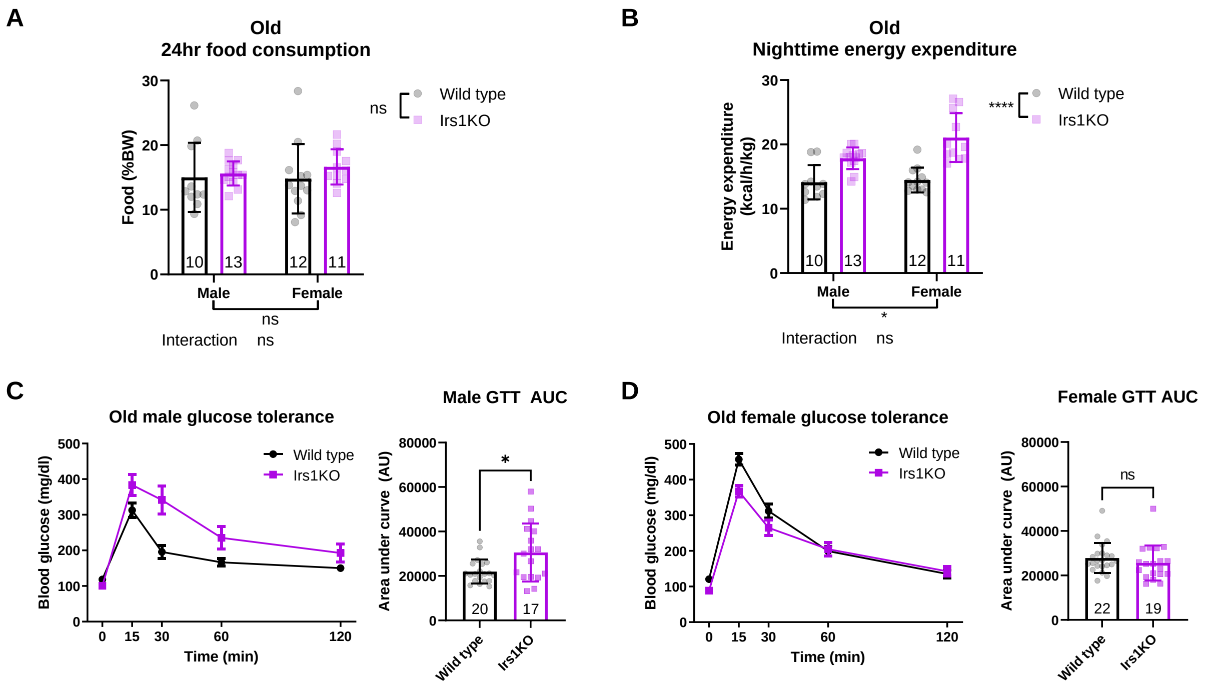


### Supplementary Figure 2: Additional parameters of old Irs1KO

**(A)** Old (16 months) Irs1KO and wild type food consumption measured in single housed animals shows no significant differences. **(B)** Body weight normalised energy expenditure of old Irs1KO mice during nighttime in individually housed animals showing significant increase in old Irs1KO mice. Glucose tolerance test (GTT) was administered to old male **(C)** and female **(D)** Irs1KO mice, by measuring blood glucose levels in response to a body weight adjusted glucose bolus. Old male Irs1KO mice showed significant reduction in glucose sensitivity. Detailed statistical values found in Table S1.

*Supplementary Figure 3*

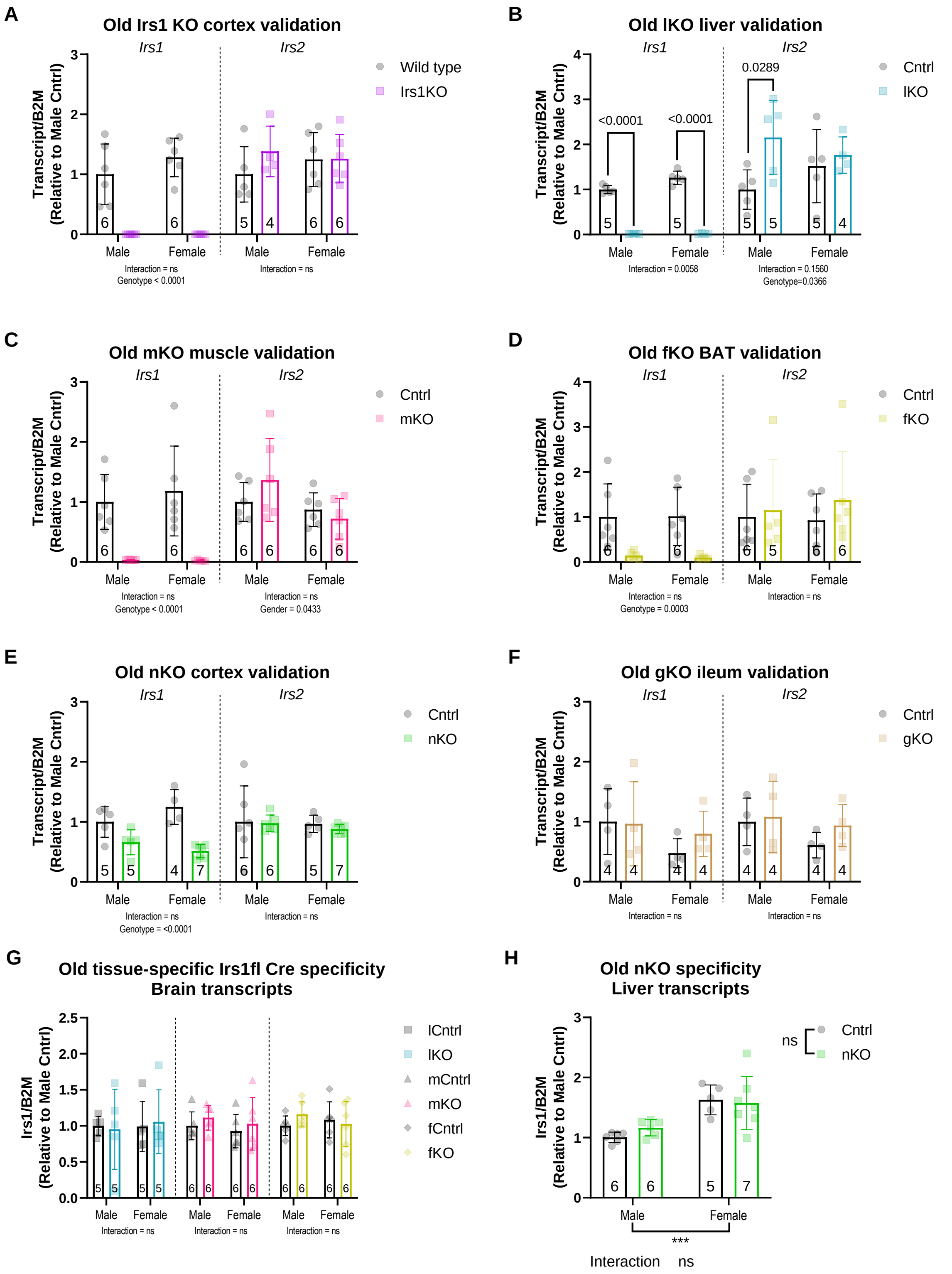


### Supplementary Figure 3: Validation of mouse models used in the study

**(A)** Quantitative real-time PCR of cortical tissue in Irs1KO mice shows depletion of Irs1 transcripts. Irs2 transcript levels show no compensatory effect. **(B)** Liver samples of lKO mice show depletion of Irs1 transcripts and compensatory Irs2 upregulation in male lKO mice. **(C)** Hindlimb muscle samples revealed depletion of Irs1 transcript levels in mKO mice, but no effect on Irs2 transcript levels. **(D)** Supraclavicular brown adipose tissue (BAT) samples of fKO mice show depletion of Irs1 transcripts with no compensatory Irs2 upregulation. **(E)** Cortex samples of nKO mice show significant reduction but not depletion of Irs1 transcripts in nKO mice, with no effect on Irs2 transcript levels. **(F)** lKO, mKO and fKO cortical samples used to assess Irs1 transcript levels did not reveal any non-specific Irs1 deletion in brain tissue. **(G)** Liver samples from nKO mice not showing any non-specific Irs1 deletion in liver tissue. All error bars correspond to standard deviation. Number of animals reported at the bottom of the bars or in figure legends. Detailed statistical values found in Table S1.

*Supplementary Figure 4*

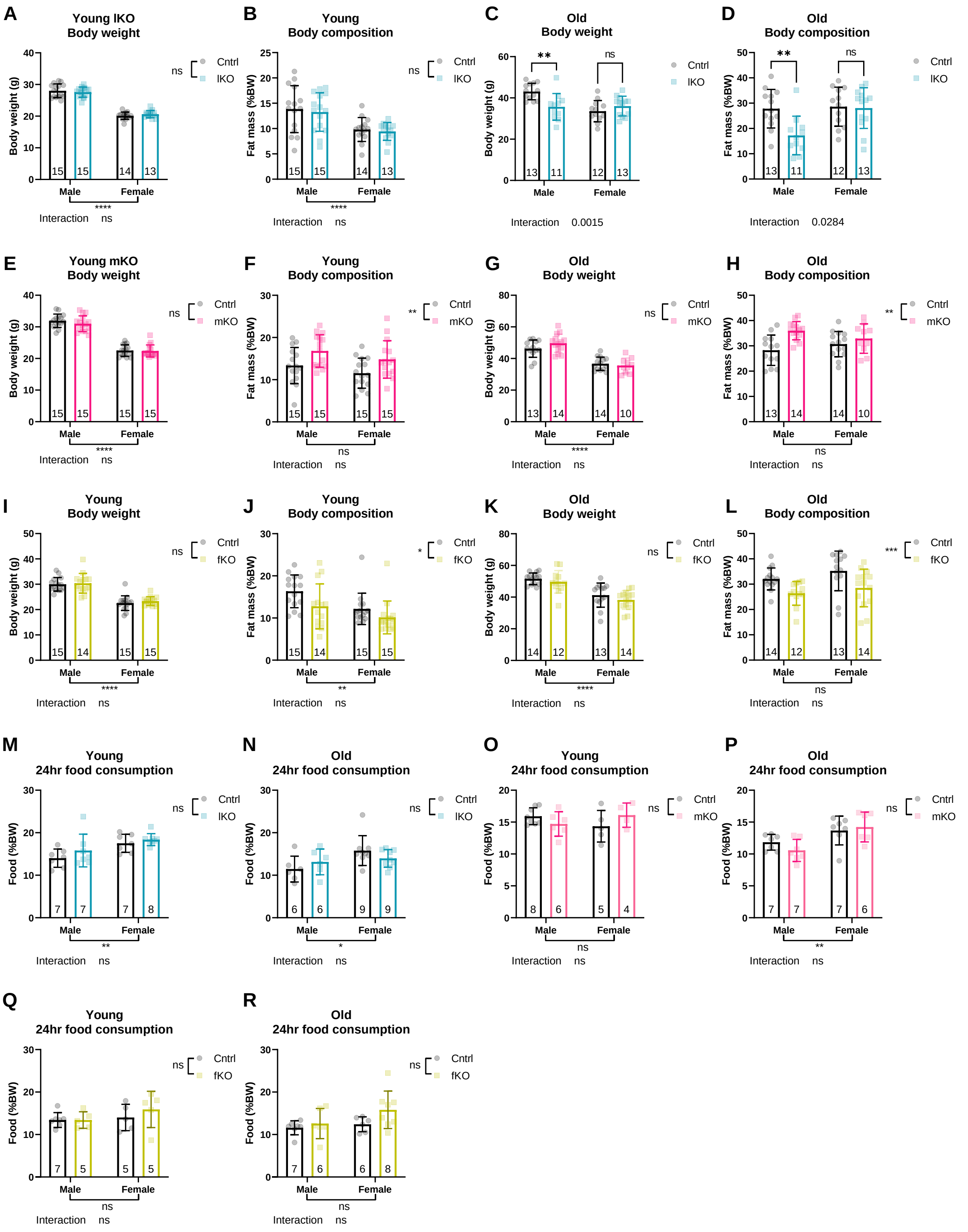


#### Supplementary Figure 4: Body weight and composition of tissue-specific Irs1KO mice

Body weight **(A)** and body composition **(B)** of young (4 months) male and female lKO mice revealed no significant difference between lKO and control littermates. **(C)** Body weight of old (16 months) male and female lKO revealed a sex-specific significant reduction in body weight of male lKO mice. **(D)** Body composition at old age also revealed sex-specific significant reduction in body weight adjusted fat mass of male lKO mice. Body weight of young **(E)** (4 months) and old **(F)** (16 months) mKO mice showed no significant difference. Body composition in young **(G)** and old **(H)** mKO mice revealed a significant age-independent increase in fat mass of mKO mice compared to control littermates. Body weight of young **(I)** (4 months) and old **(K)** (16 months) fKO mice showed no significant difference. Body composition in young **(J)** and old **(L)** fKO mice revealed a significant age-independent decrease in fat mass of fKO mice compared to control littermates. Measurement of food consumption of young lKO **(M)** and old lKO **(N)** revealed no significant difference in food consumption of single housed animals relative to the corresponding littermate controls. Measurement of food consumption of young mKO **(O)** and old mKO **(P)** revealed no significant difference in food consumption of single housed animals relative to the corresponding littermate controls. Measurement of food consumption of young fKO **(Q)** and old fKO **(R)** revealed no significant difference in food consumption of single housed animals relative to the corresponding littermate controls. Number of animals reported at the bottom of the bars for each condition. Detailed statistical values found in Table S1.

*Supplementary Figure 5*

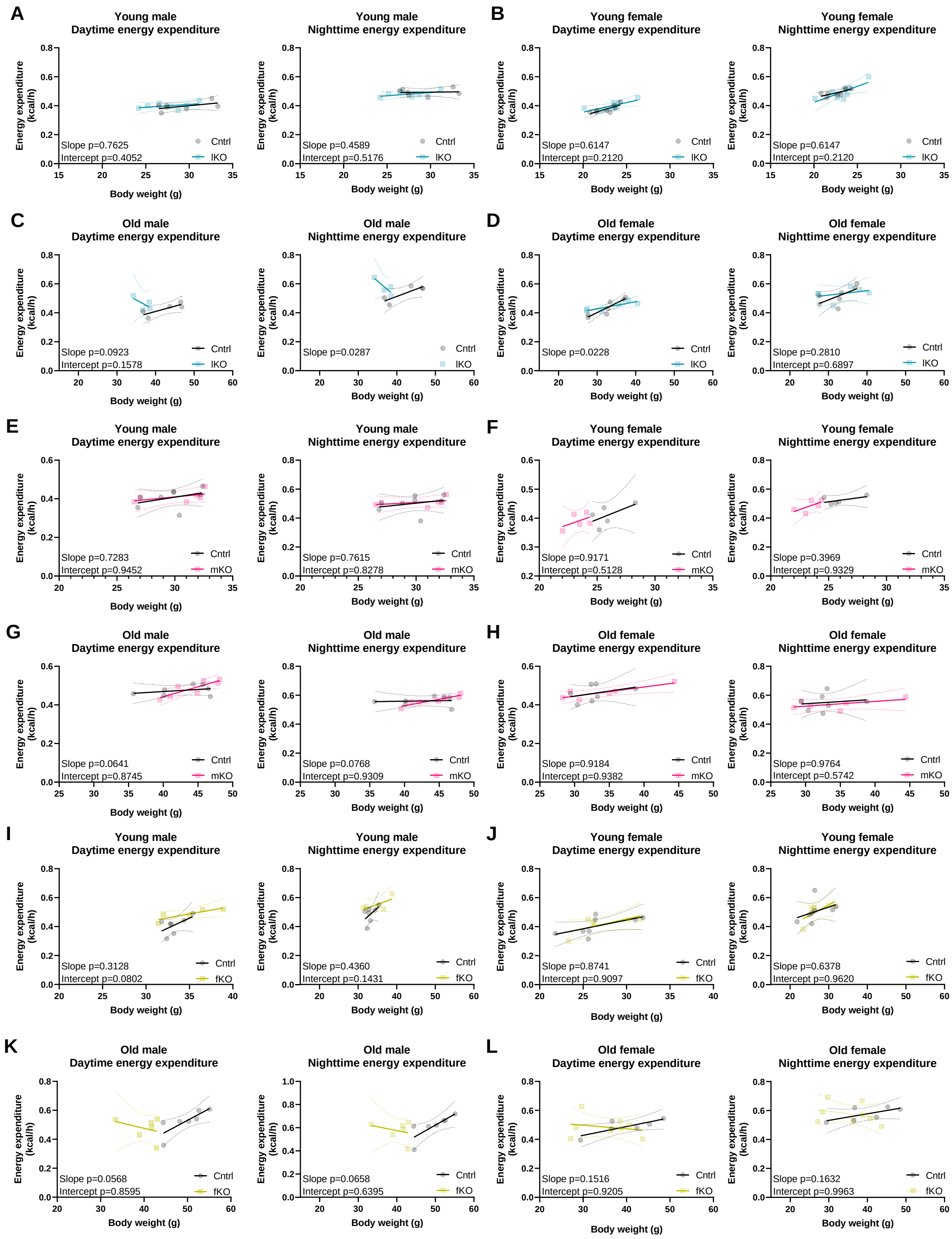


### Supplementary Figure 5: Energy expenditure of tissue-specific Irs1KO mice

Daytime and nighttime energy expenditure were analysed by linear regression of energy expenditure by body weight (ANCOVA). No difference in energy expenditure was detected in young male **(A)** or female **(B)** lKOs (male control and lKO n=7, female controls and lKO n=8) or old male **(C)** and female **(D)** lKOs (male control n=5 and lKO n=4, female controls n=7 and lKO n=6). No difference in energy expenditure was detected in young male **(E)** and female **(F)** mKOs (male control n=8 and mKO n=6, female controls and mKO n=5) or old male **(G)** and female **(H)** mKOs (male control and mKO n=7, female control n=7 and mKO n=6). No difference in energy expenditure was detected in young male **(I)** and female **(J)** fKOs (male control n=7 and fKO n=5, female control n=8 and fKO n=6) or old male **(K)** and female **(L)** fKOs (male control n=7 and fKO n=6, female control n=6 and fKO n=8). Detailed statistical values found in Table S1.

*Supplementary Figure 6*

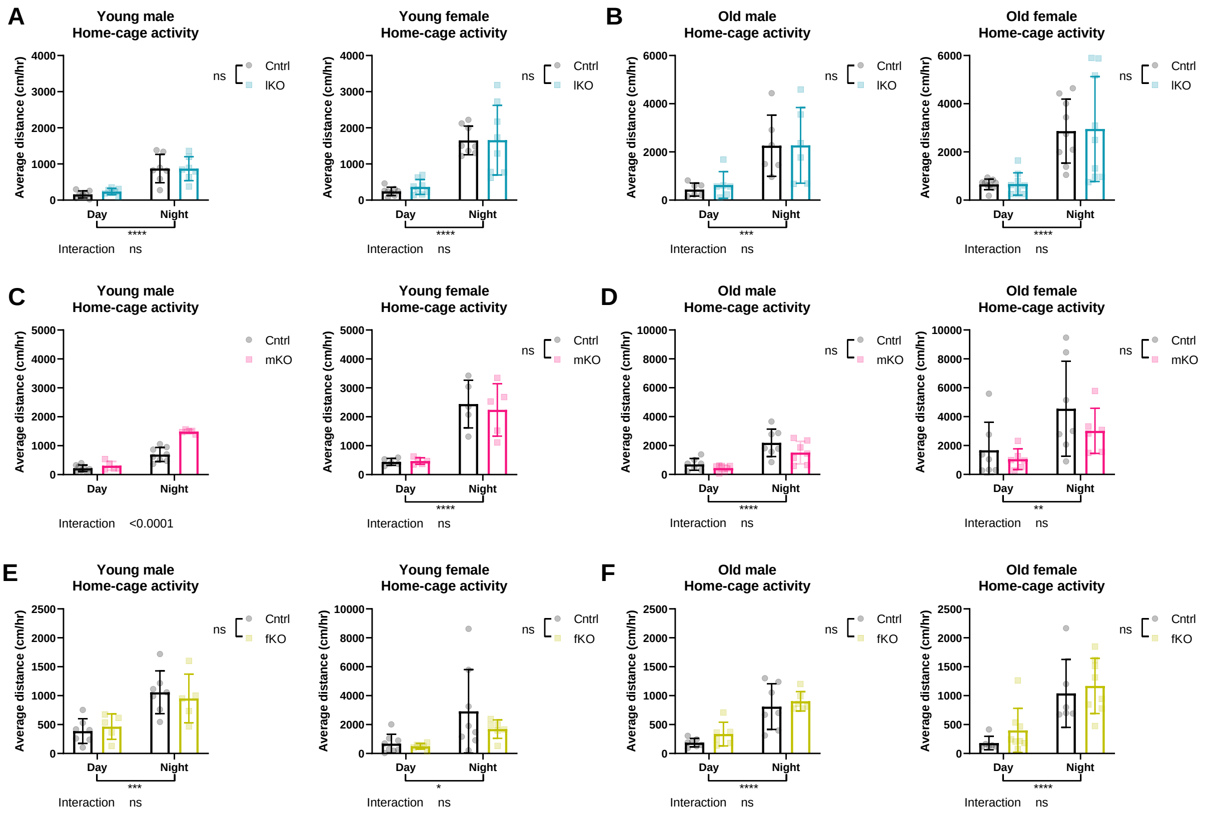


### Supplementary Figure 6: Locomotor activity of tissue-specific Irs1KO mice

Measurement of spontaneous locomotor activity of singly housed young lKO **(A)** (male control and lKO n=7, female controls and lKO n=8) and old lKO **(B)** (male control and lKO n=6, female controls and lKO n=9) mice during daytime and nighttime did not reveal any significant difference due to genotype. Spontaneous locomotor activity of singly housed young male mKO mice **(C)** (control n=8 and mKO n=5) revealed higher activity levels during nighttime compared to controls. Spontaneous activity of young female mKO **(C)** (control and mKO n=5), old male mKO **(D)** (control and mKO n=7) or old female mKO **(D)** (control n=7 and mKO n=6) mice during daytime and nighttime did not reveal any significant difference due to genotype. Spontaneous locomotor activity of singly housed young fKO **(E)** (male control n=7 and fKO n=5, female controls n=8 and fKO n=6) and old fKO **(F)** (male control n=7 and fKO n=6, female controls n=6 and fKO n=8) mice during daytime and nighttime did not reveal any significant differences. Detailed statistical values found in Table S1.

*Supplementary Figure 7*

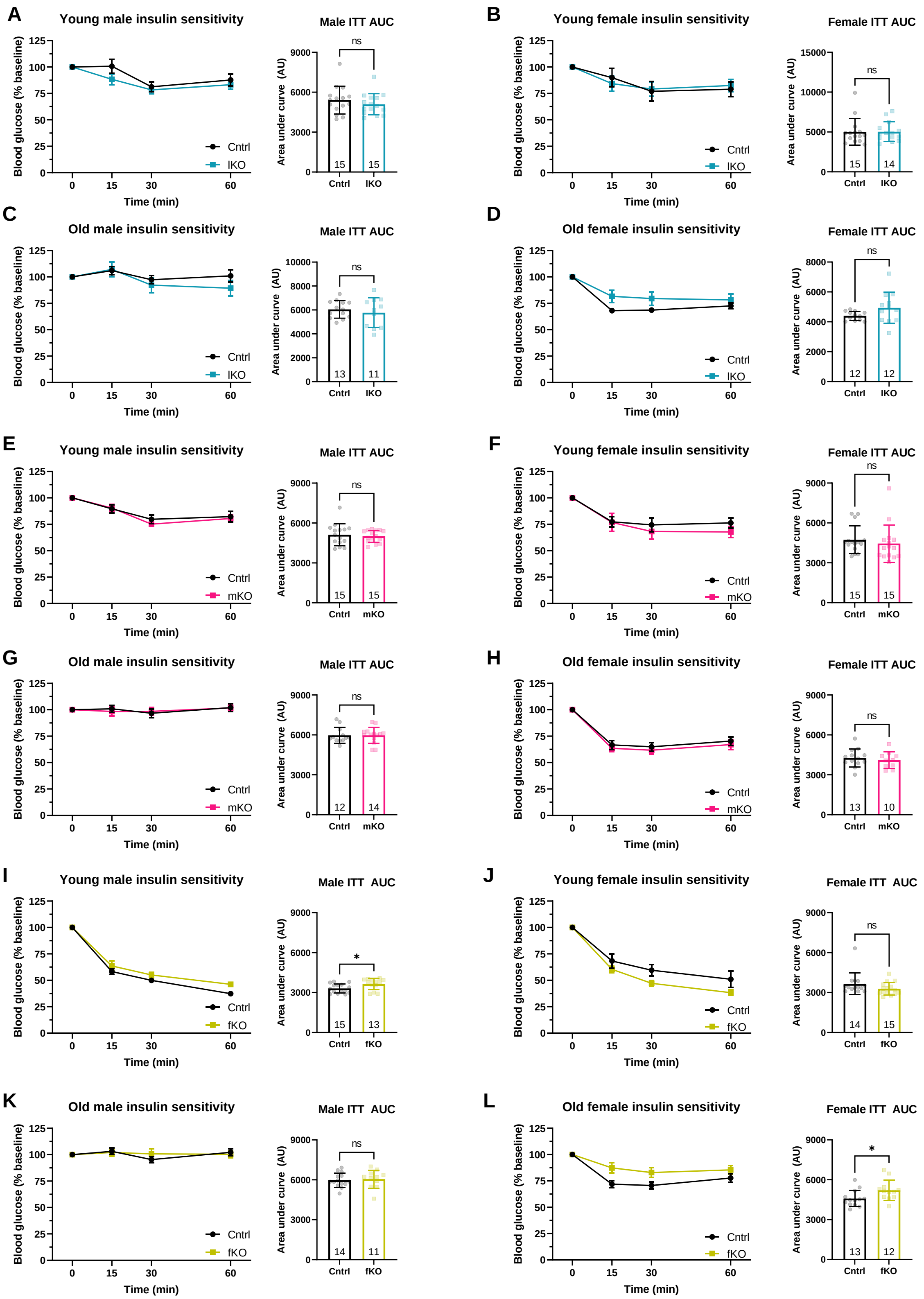


**Supplementary Figure 7: Insulin tolerance test of tissue-specific Irs1KO mice**

Insulin tolerance test (ITT) revealed no significant difference in insulin sensitivity of young male lKO **(A)**, young female lKO **(B),** old male lKO **(C)** and old female lKO **(D)** compared to their respective control littermates as assessed by AUC analysis. No significant difference in insulin sensitivity of young male mKO **(E)**, young female mKO **(F),** old male mKO **(G)** and old female mKO **(H)** compared to their respective control littermates as assessed by AUC analysis. ITT of young male fKO **(I)** revealed a significant reduction in insulin sensitivity between fKO and control littermates. ITT analysis of young female fKO **(J)** and old male fKO **(K)** did not detect any significant differences between fKO mice and their control littermates. ITT of old female fKO **(L)** revealed a significant reduction in insulin sensitivity between fKO and control littermates. Detailed statistical values found in Table S1.

*Supplementary Figure 8*

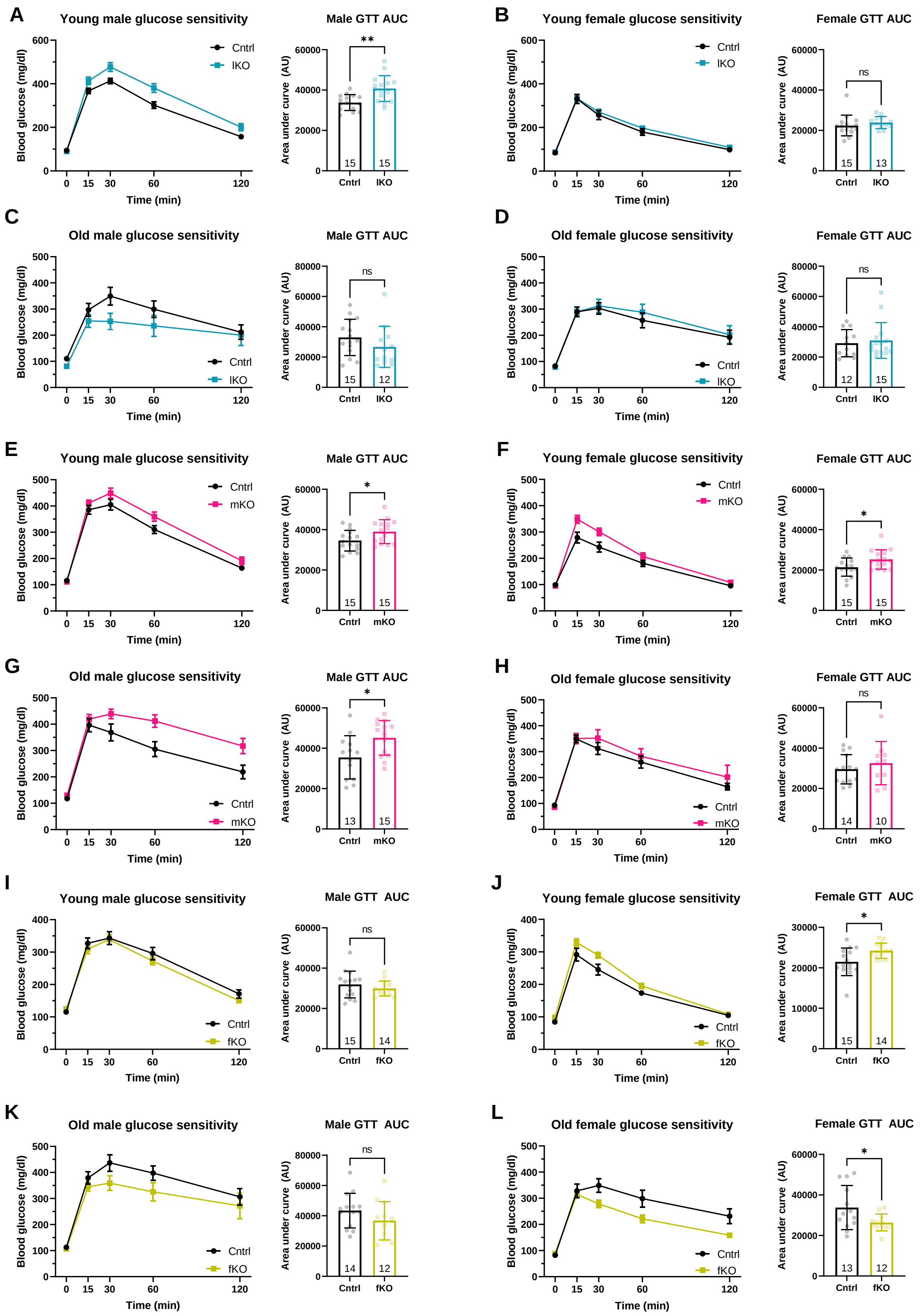


**Supplementary Figure 8: Glucose tolerance test of tissue-specific Irs1KO mice (below)**

Glucose tolerance test (GTT) revealed a significant reduction in glucose tolerance of young male lKO **(A)** compared to control mice as assessed by AUC analysis. AUC analysis of GTT of young female lKO **(B)**, old male lKO **(C)** and old female lKO **(D)** did not detect any significant differences between lKO mice and their control littermates. GTT revealed a significant reduction in glucose tolerance of young male mKO **(E)**, young female mKO **(F)** and old male mKO **(G)** as assessed by AUC analysis. GTT of old female mKO **(H)** did not detect any significant differences between mKO mice and their control littermates. GTT of young male fKO **(I)** did not detect any significant differences between fKO mice and their control littermates. GTT of young female fKO **(J)** revealed a significant reduction in glucose tolerance as assessed by AUC analysis. GTT of old male fKO **(K)** and old female fKO **(L)** did not detect any significant differences between fKO mice and their control littermates. Detailed statistical values found in Table S1.

*Supplementary Figure 9*

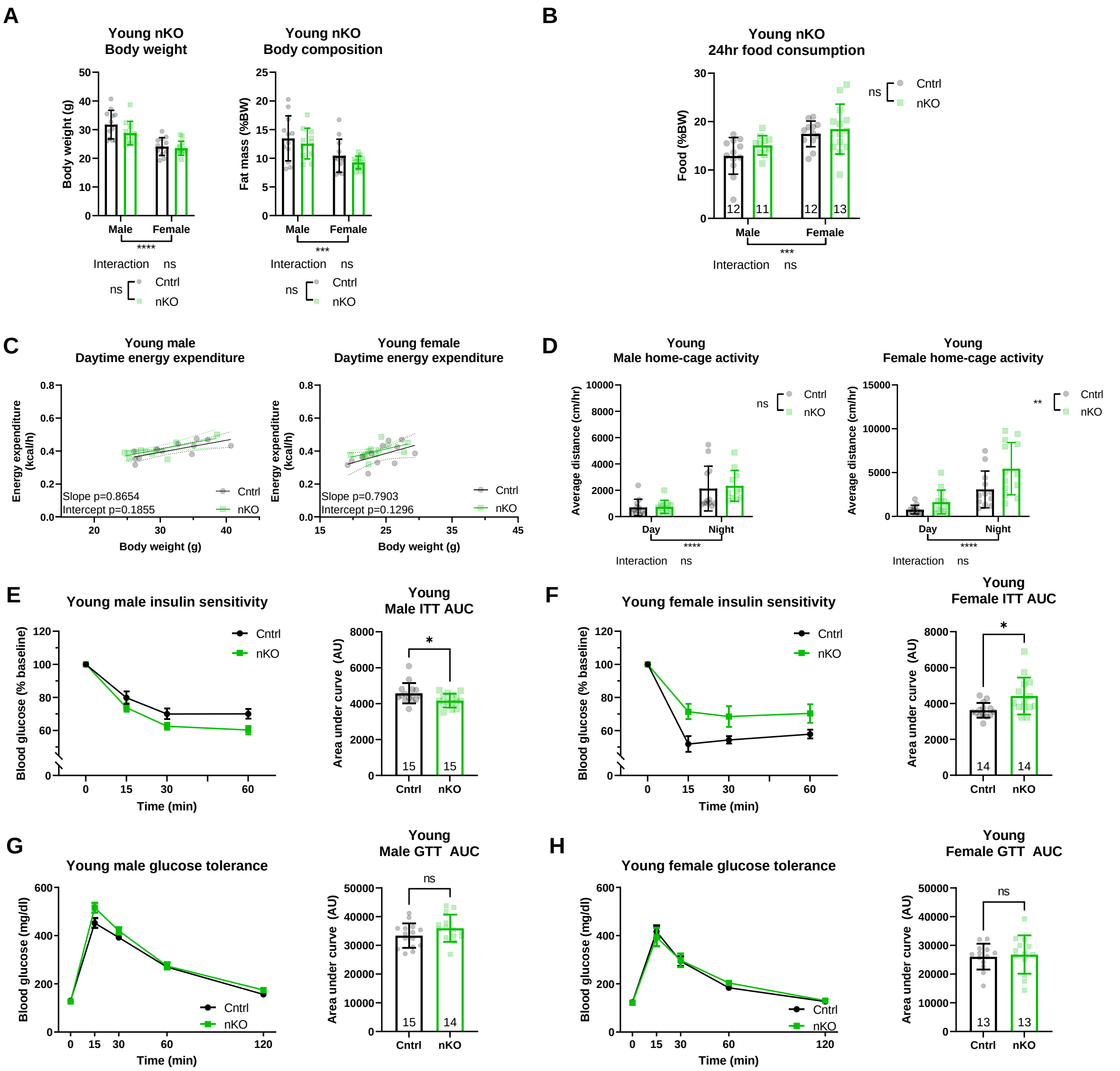


### Supplementary Figure 9: Characterisation of young nKO

**(A)** No significant difference was observed in body weight or body composition at young age (3 months) of nKO and control littermates (male nKO and controls n=15, female nKO and control n=14). **(B)** Measurement of food consumption of young nKO and control mice revealed no significant difference in food consumption of single housed animals. **(C)** Daytime energy expenditure of male nKO and female nKO and control mice was analysed by linear regression of energy expenditure by body weight (ANCOVA). No difference in energy expenditure between young male nKO (n=11) and controls (n=12) or female nKO (n=13) and controls (n=12) was observed. **(D)** No significant difference was observed in spontaneous activity of young single-housed male nKO (n=12) and littermate control (n=11) mice. A significant increase in daytime and nighttime activity was observed between female nKO (n=13) and littermate control (n=12) mice. Insulin sensitivity of male nKO and control mice **(E)** showed a significantly improved insulin sensitivity in male nKO mice. However, a significant reduction in insulin sensitivity was detected between female **(F)** nKO and control mice. Analysis of GTT of male **(G)** and female **(H)** nKO mice did not reveal any significant difference in young nKO glucose tolerance compared to control littermates. All error bars correspond to standard deviation except for longitudinal glucose and insulin sensitivity where standard error of the mean is reported. Number of animals reported at the bottom of the bars for each condition. Detailed statistical values found in Table S1.

*Supplementary Figure 10*

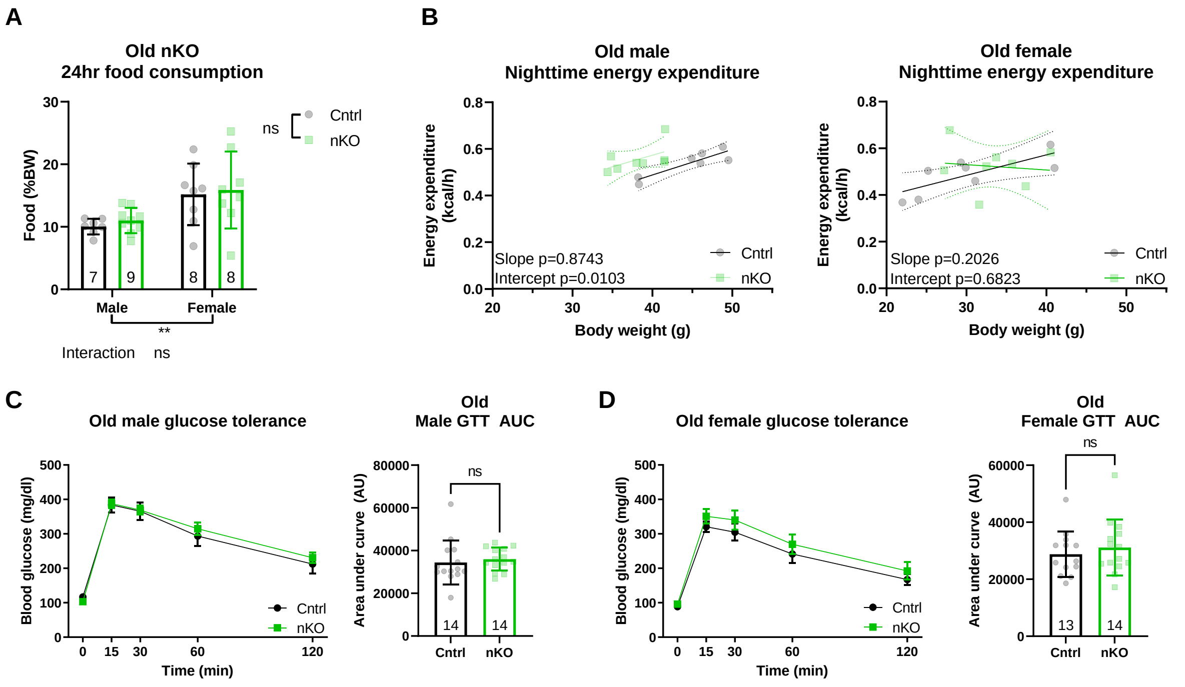


### Supplementary Figure 10: Additional parameters of old nKO

**(A)** No difference in food consumption was detected in single housed old (16 months) nKO and control littermates. **(B)** Nighttime energy expenditure of male and female mice was analysed by linear regression of energy expenditure by body weight (ANCOVA). Unlike female nKO mice (n=8 female control and nKO mice) where no significant difference between intercepts was detected, male nKO (n=8) mice showed significant increase in energy expenditure compared to male controls (n=7). Analysis of GTT did not detect any significant difference between the ability of old male **(C)** and female **(D)** nKO mice compared to their respective littermate controls in lowering blood glucose levels. All error bars correspond to standard deviation except for longitudinal glucose sensitivity where standard error of the mean is reported. Number of animals reported at the bottom of the bars for each condition. Detailed statistical values found in Table S1.

*Supplementary Figure 11*

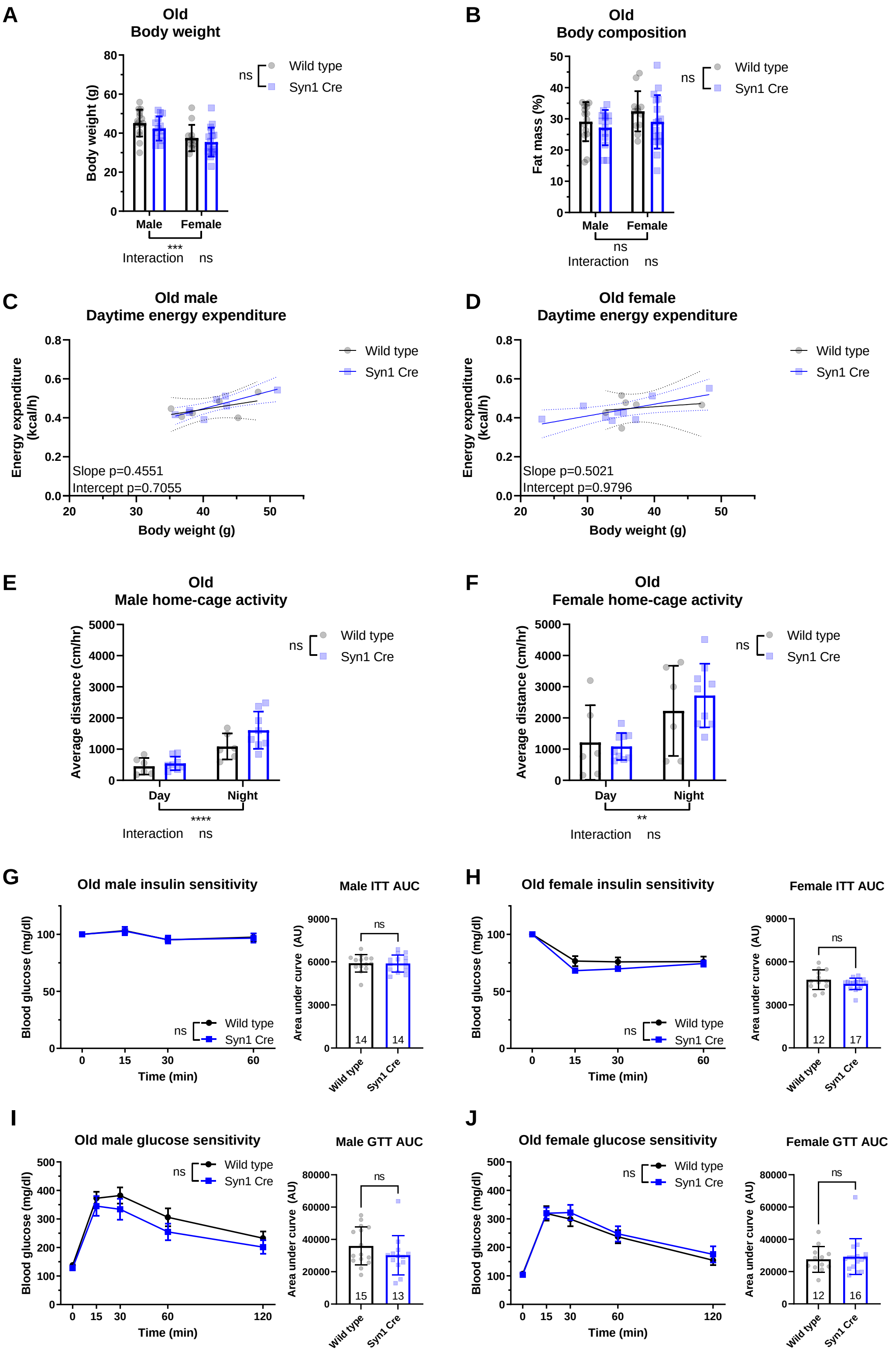


**Supplementary Figure 11: Neuronal Syn1Cre expression does not affect peripheral metabolism (below)**

**(A)** No significant difference was observed in body weight or **(B)** body composition at old age (16 months) of Syn1Cre and wild type littermates (male wild type n=15 and Syn1Cre n=14, female wild type n=12 and Syn1Cre n=17). Daytime energy expenditure of **(C)** male and **(D)** female Syn1Cre and wild type mice was analysed by linear regression of energy expenditure by body weight (ANCOVA). No difference in energy expenditure between old male Syn1Cre (n=8) and wild type (n=6) as well as between female Syn1Cre (n=9) and wild type (n=6) was observed. No significant difference was observed in spontaneous activity of old single-housed **(E)** male Syn1Cre (n=8) and littermate wild type (n=6) mice or female **(F)** Syn1Cre (n=9) and wild type (n=6) mice. Insulin sensitivity of old **(G)** male or **(H)** female Syn1Cre and wild type mice showed no significant difference in insulin sensitivity. Analysis of GTT of male **(I)** and female **(J)** Syn1Cre mice did not reveal any significant difference in old Syn1Cre glucose tolerance compared to wild type littermates. All error bars correspond to standard deviation except for longitudinal glucose and insulin sensitivity where standard error of the mean is reported. Number of animals reported at the bottom of the bars for each condition. Detailed statistical values found in Table S1.

*Supplementary Figure 12*

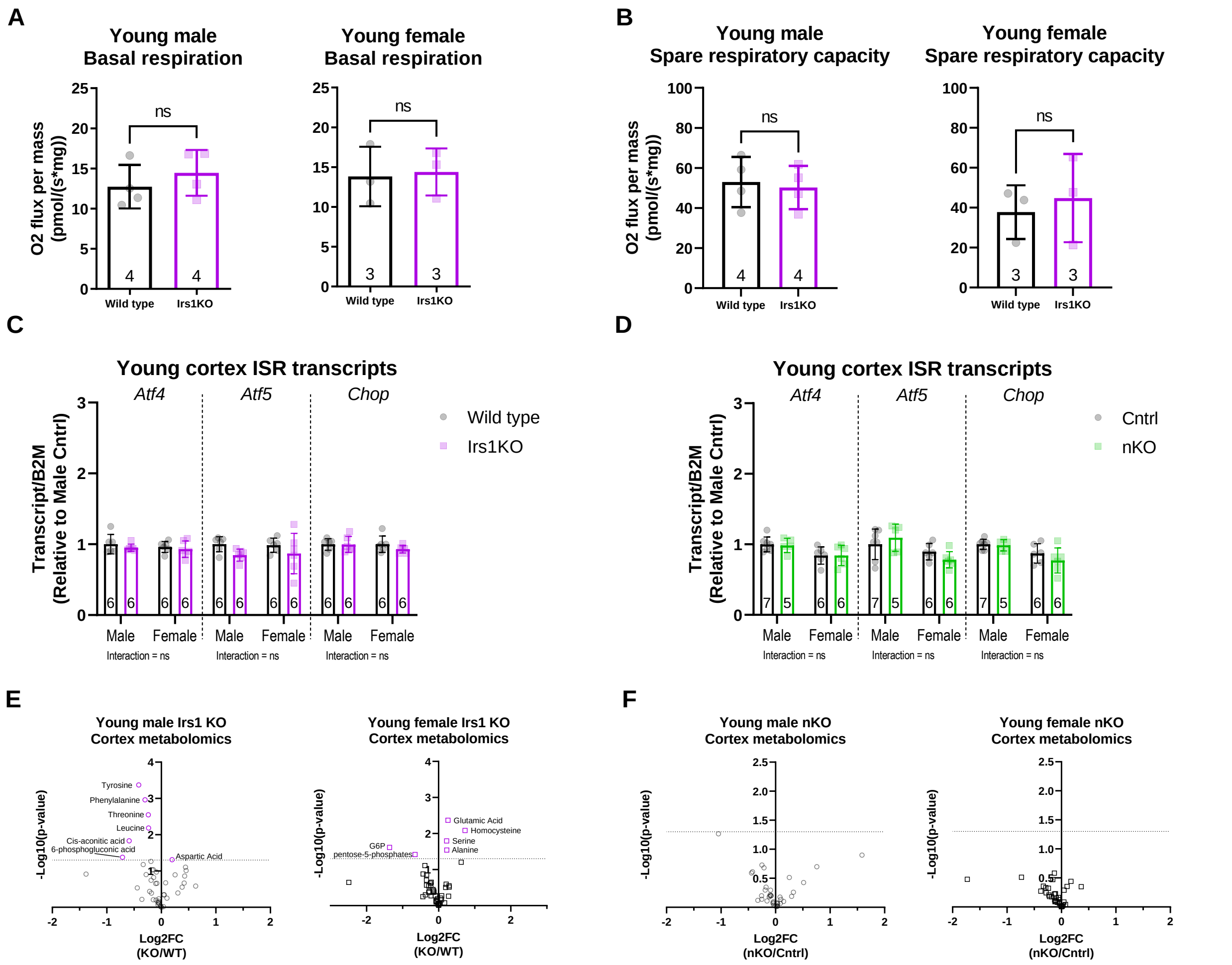


### Supplementary Figure 12: No activation of ISR in brains of young Irs1KO and nKO mice

**(A)** Basal oxygen consumption of brain tissue showed no difference in the basal respiration of mitochondria of young (6 months) male or female Irs1KO mice. **(B)** Mitochondrial spare respiratory capacity in brain tissue revealed no significant difference in young male or female Irs1KO mitochondrial function. **(C)** Quantitative real-time PCR was performed on brains of young male and female Irs1KO mice and their wild type littermates to measure transcript levels of several integrated stress response (ISR) markers revealed no significant differences. **(D)** Transcripts of ISR markers were measured in brains of young male and female nKO mice and their control littermates showed no significant differences. **(E)** Semi-targeted metabolomics revealed down-regulation in metabolites in male Irs1KO mice and up-regulation in metabolites in female Irs1KO mice (male wild type and Irs1KO n=6, female wild type and Irs1KO n=6). **(F)** No significant change in metabolites in either male or female young nKO brain tissue (male control n=7 and nKO n=5, female control and nKO n=6). All error bars correspond to standard deviation. Detailed statistical values found in Table S1. Full metabolites measured in Table S2.

*Supplementary Figure 13*

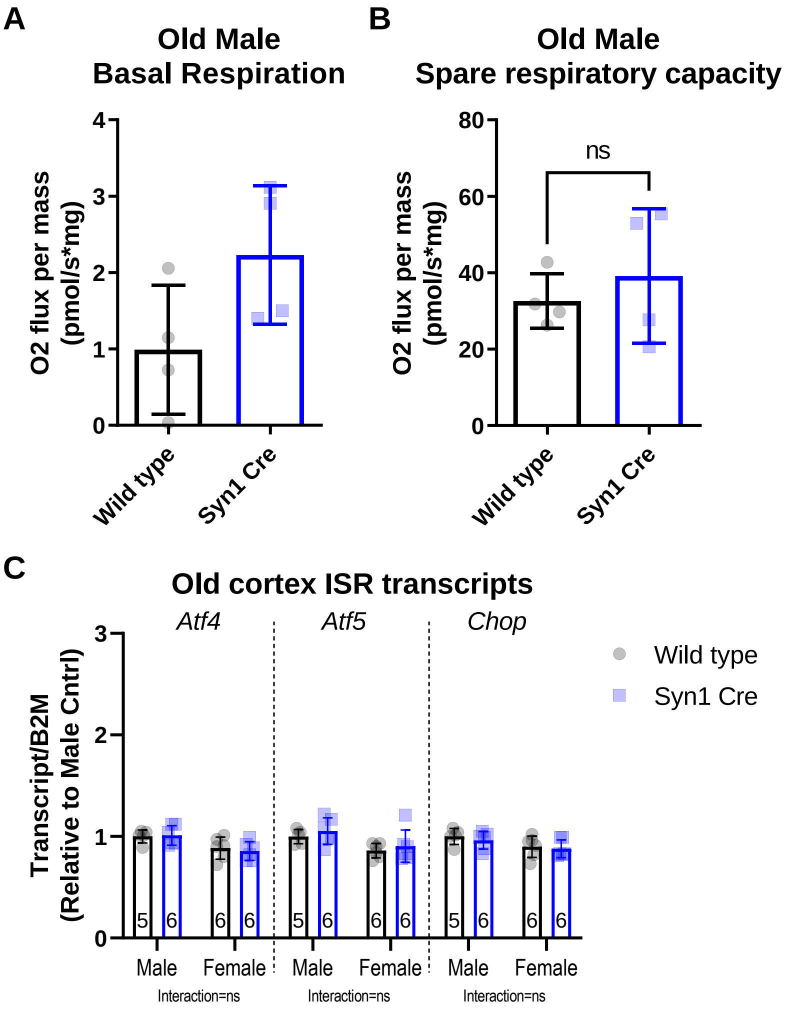


**Supplementary Figure 13: Syn1Cre expression does not lead to brain ISR activation**

**(A)** Basal oxygen consumption of brain tissue showed no difference in the basal respiration of mitochondria of old (22 months) male Syn1Cre mice compared to their wild type littermates. **(B)** Mitochondrial spare respiratory capacity in brain tissue revealed no significant difference in old male Syn1Cre mitochondrial function. **(C)** Quantitative real-time PCR was performed on brains of old male and female Syn1Cre mice and their wild type littermates to measure transcript levels of several integrated stress response (ISR) markers revealed no significant differences. All error bars correspond to standard deviation. Detailed statistical values found in Table S1.

*Supplementary Figure 14*

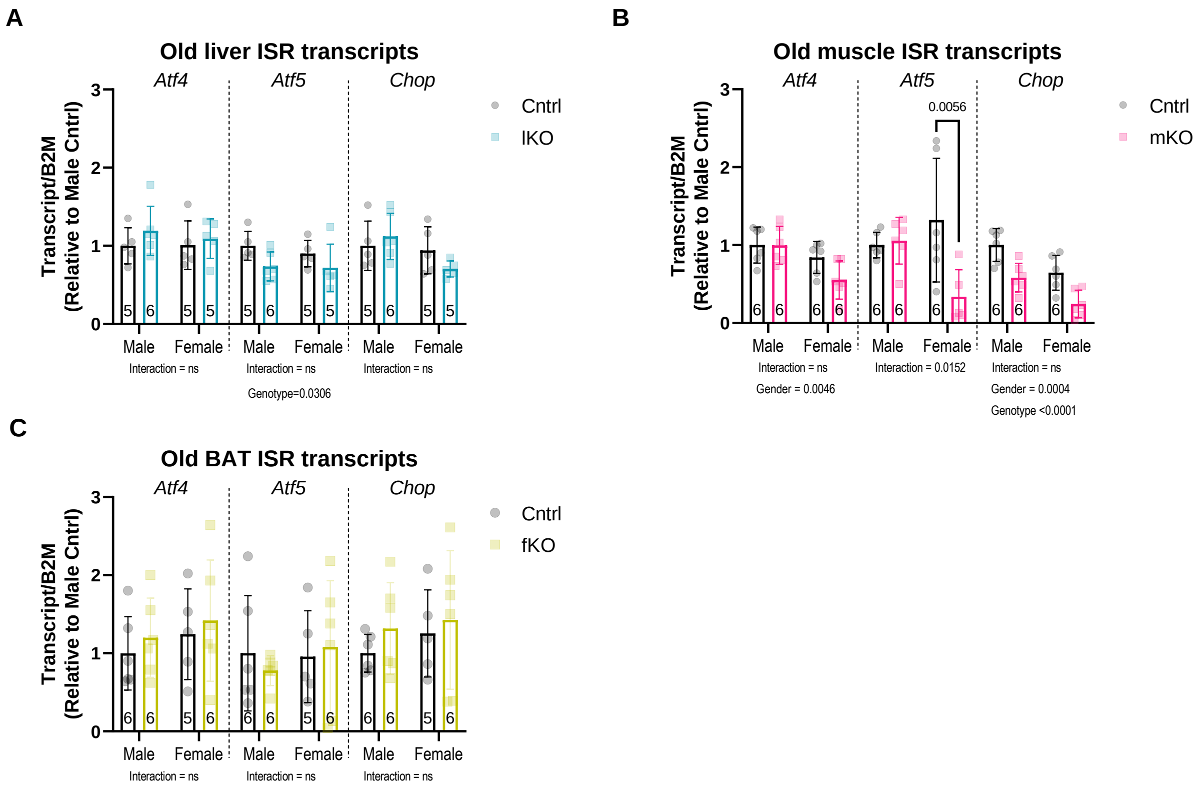


#### Supplementary Figure 14: IRS1 deletion in peripheral tissues is insufficient to induce local ISR signature in old mice

Quantitative real-time PCR performed on metabolic organs of Irs1 tissue-specific deletion in old (16 months) mice targeting various integrated stress response (ISR) markers. **(A)** No significant difference in ISR transcripts was detected in the liver of lKO mice (male control n=5 and lKO n=6, female control n=5 and lKO n=5). **(B)** No significant increase in ISR transcript levels in hind limb muscle tissue of mKO mice, however we did detect a sex-specific downregulation of Atf5 transcripts in female mKO mice and genotype specific downregulation of Chop levels in mKO mice (n=6 biologically independent animals for all groups). **(C)** No difference in ISR transcripts was found in supraclavicular brown adipose tissue (BAT) of fKO mice (male control n=6 and fKO n=6, female control n=5 and fKO n=6). Detailed statistical values found in Table S1.

*Supplementary Figure 15*

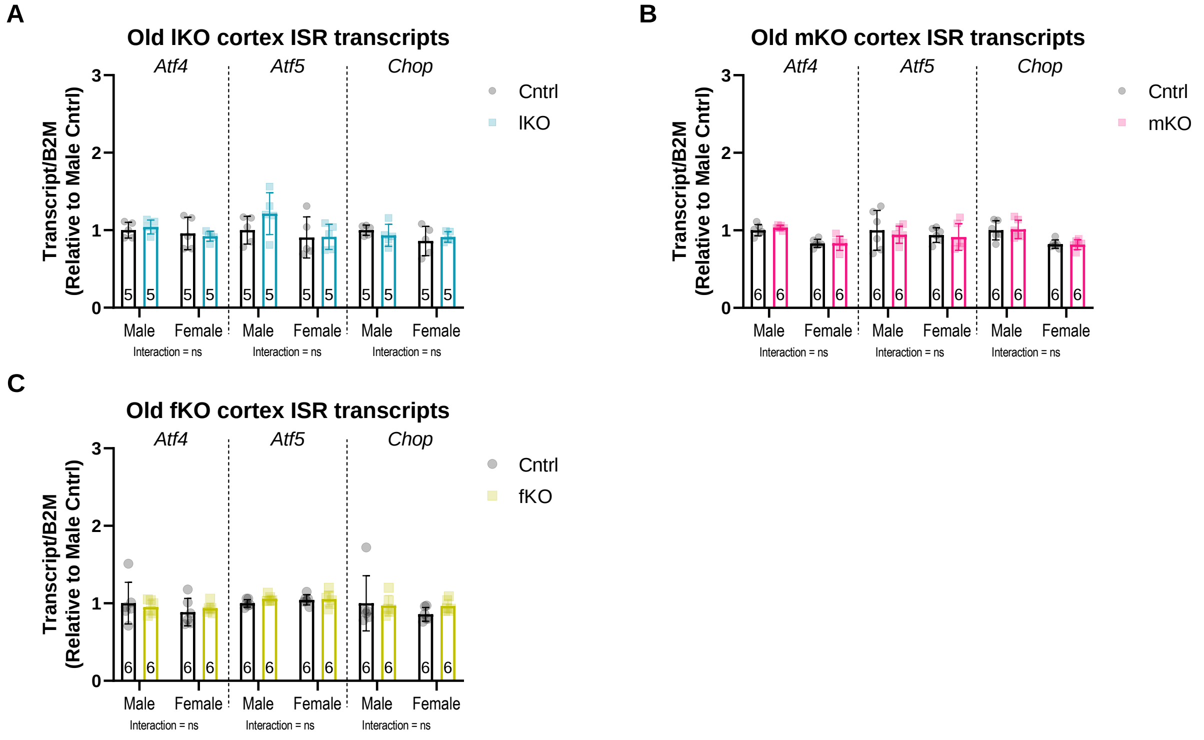


#### Supplementary Figure 15: IRS1 deletion in peripheral tissues does not induce ISR signature in brains of old mice

Quantitative real-time PCR performed on cortex samples of Irs1 tissue-specific deletion in old (16 months) mice targeting various integrated stress response (ISR) markers. No significant difference in ISR transcripts was detected in **(A)** lKO, **(B)** mKO, or **(C)** fKO mice. Detailed statistical values found in Table S1.

### Supplementary Table 1

**Figure 1: Increased lifespan and improved health parameters in Irs1KO mice**

**Panel a**

| Type of test | Comparison | P value<br>(Multiplicity adjusted P value reported for Post-tests) | Test statistic | Degrees of Freedom<br>(F(df numerator, df denominator)) |
|---|---|---|---|---|
| Log-rank test for median lifespan | Male Irs1 KO vs wild type | 0,0021 | 9,496 | 1 |
| Mann Whitney test for max lifespan | Male Irs1 KO vs wild type | < 0,0001 | | |

**Panel b**

| Type of test | Comparison | P value | Test statistic | Degrees of Freedom |
|---|---|---|---|---|
| Log-rank test for median lifespan | Female Irs1 KO vs wild type | 0,0056 | 7,667 | 1 |
| Mann Whitney test for max lifespan | Female Irs1 KO vs wild type | 0,0063 | | |

**Panel c**

Old Irs1 KO body weight

| Type of test | Comparison | P value | Test statistic | Degrees of Freedom |
|---|---|---|---|---|
| Two-way ANOVA | Interaction (sex*genotype) | 0,9049 | 0,01438 | F (1, 74) |
| Two-way ANOVA | Sex (main effect) | 0,4678 | 0,5327 | F (1, 74) |
| Two-way ANOVA | Genotype (main effect) | <0,0001 | 410,3 | F (1, 74) |
| Sidak's Post-test | Male Irs1 KO vs wild type | <0,0001 | 14,05 | 74 |
| Sidak's Post-test | Female Irs1 KO vs wild type | <0,0001 | 14,62 | 74 |

Old Irs1 KO body composition

| Type of test | Comparison | P value | Test statistic | Degrees of Freedom |
|---|---|---|---|---|
| Two-way ANOVA | Interaction (sex*genotype) | 0,1787 | 1,844 | F (1, 74) |
| Two-way ANOVA | Sex (main effect) | <0,0001 | 136,1 | F (1, 74) |
| Two-way ANOVA | Genotype (main effect) | <0,0001 | 33,97 | F (1, 74) |
| Sidak's Post-test | Male Irs1 KO vs wild type | <0,0001 | 3,082 | 74 |
| Sidak's Post-test | Female Irs1 KO vs wild type | <0,0001 | 5,219 | 74 |

**Panel d**

Old Irs1 KO average energy expenditure during daytime

| Type of test | Comparison | P value | Test statistic | Degrees of Freedom |
|---|---|---|---|---|
| Two-way ANOVA | Interaction (sex*genotype) | 0,2969 | 1,116 | F (1, 42) |
| Two-way ANOVA | Sex (main effect) | 0,0143 | 6,53 | F (1, 42) |
| Two-way ANOVA | Genotype (main effect) | <0,0001 | 56,11 | F (1, 42) |
| Sidak's Post-test | Male Irs1 KO vs wild type | <0,0001 | 4,533 | 42 |
| Sidak's Post-test | Female Irs1 KO vs wild type | <0,0001 | 6,067 | 42 |

**Panel e**

Old Irs1 KO male activity

| Type of test | Comparison | P value | Test statistic | Degrees of Freedom |
|---|---|---|---|---|
| Two-way ANOVA | Interaction (phase*genotype) | 0,039 | 4,539 | F (1, 42) |
| Two-way ANOVA | Phase (main effect) | <0,0001 | 109,9 | F (1, 42) |
| Two-way ANOVA | Genotype (main effect) | 0,0005 | 14,18 | F (1, 42) |
| Sidak's Post-test | Male Irs1 KO vs wild type day | 0,4437 | 1,156 | 42 |
| Sidak's Post-test | Male Irs1 KO vs wild type night | 0,0003 | 4,169 | 42 |

Old Irs1 KO female activity

| Type of test | Comparison | P value | Test statistic | Degrees of Freedom |
|---|---|---|---|---|
| Two-way ANOVA | Interaction (phase*genotype) | 0,0242 | 5,468 | F (1, 42) |
| Two-way ANOVA | Phase (main effect) | <0,0001 | 122,8 | F (1, 42) |
| Two-way ANOVA | Genotype (main effect) | 0,0216 | 5,699 | F (1, 42) |
| Sidak's Post-test | Female Irs1 KO vs wild type day | 0,9993 | 0,03453 | 42 |
| Sidak's Post-test | Female Irs1 KO vs wild type night | 0,0035 | 3,341 | 42 |

**Panel f**

Old Irs1 KO male insulin tolerance test

| Type of test | Comparison | P value | Test statistic | Degrees of Freedom |
|---|---|---|---|---|
| Two-sided t test | AUC male Irs1 KO vs wild type | 0,0162 | 2,525 | 35 |

**Panel g**

Old Irs1 KO female insulin tolerance test

| Type of test | Comparison | P value | Test statistic | Degrees of Freedom |
|---|---|---|---|---|
| Two-sided t test | AUC female Irs1 KO vs wild type | 0,0084 | 2,776 | 39 |

#### Figure 2: Tissue-specific deletion of IRS1 is not sufficient for lifespan extension

**Panel a**

| Type of test | Comparison | P value (Multiplicity adjusted P value reported for Post-tests) | Test statistic | Degrees of Freedom (F(df numerator, df denominator)) |
|---|---|---|---|---|
| Log-rank test for median lifespan | Male lKO vs control | 0,1071 | 2,596 | 1 |
| Mann Whitney test for max lifespan | Male lKO vs control | 0,0095 | | |

**Panel b**

| Type of test | Comparison | P value | Test statistic | Degrees of Freedom |
|---|---|---|---|---|
| Log-rank test for median lifespan | Female lKO vs control | 0,8323 | 0,04484 | 1 |
| Mann Whitney test for max lifespan | Female lKO vs control | 0,5661 | | |

**Panel c**

| Type of test | Comparison | P value | Test statistic | Degrees of Freedom |
|---|---|---|---|---|
| Log-rank test for median lifespan | Male mKO vs control | 0,0605 | 3,525 | 1 |
| Mann Whitney test for max lifespan | Male mKO vs control | 0,0303 | | |

**Panel d**

| Type of test | Comparison | P value | Test statistic | Degrees of Freedom |
|---|---|---|---|---|
| Log-rank test for median lifespan | Female mKO vs control | 0,6842 | 0,1654 | 1 |
| Mann Whitney test for max lifespan | Female mKO vs control | 0,5787 | | |

**Panel e**

| Type of test | Comparison | P value | Test statistic | Degrees of Freedom |
|---|---|---|---|---|
| Log-rank test for median lifespan | Male fKO vs control | 0,4777 | 0,5042 | 1 |
| Mann Whitney test for max lifespan | Male fKO vs control | 0,4813 | | |

**Panel f**

| Type of test | Comparison | P value | Test statistic | Degrees of Freedom |
|---|---|---|---|---|
| Log-rank test for median lifespan | Female fKO vs control | 0,029 | 4,767 | 1 |
| Mann Whitney test for max lifespan | Female fKO vs control | 0,0029 | | |

**Panel g**

| Type of test | Comparison | P value | Test statistic | Degrees of Freedom |
|---|---|---|---|---|
| Log-rank test for median lifespan | Male nKO vs control | 0,0874 | 2,921 | 1 |
| Mann Whitney test for max lifespan | Male nKO vs control | 0,2393 | | |

**Panel h**

| Type of test | Comparison | P value | Test statistic | Degrees of Freedom |
|---|---|---|---|---|
| Log-rank test for median lifespan | Female nKO vs control | 0,8656 | 0,02865 | 1 |
| Mann Whitney test for max lifespan | Female nKO vs control | 0,1051 | | |

**Figure 3: Neuron-specific Irs1 knockout (nKO) mice show male-specific improvement in metabolic health**

| | | Multiplicity adjusted P value reported for Post-tests | | F(df numerator, df denominator) |
|---|---|---|---|---|

**Panel a**

Old nKO body weight

| Type of test | Comparison | P value | Test statistic | Degrees of Freedom |
|---|---|---|---|---|
| Two-way ANOVA | Interaction (sex*genotype) | 0,0172 | 6,069 | F (1, 51) |
| Two-way ANOVA | Sex (main effect) | <0,0001 | 40,08 | F (1, 51) |
| Two-way ANOVA | Genotype (main effect) | 0,0327 | 4,819 | F (1, 51) |
| Sidak's Post-test | Male nKO vs control | 0,0033 | 3,326 | 51 |
| Sidak's Post-test | Female nKO vs control | 0,978 | 0,1879 | 51 |

Old nKO body composition

| Type of test | Comparison | P value | Test statistic | Degrees of Freedom |
|---|---|---|---|---|
| Two-way ANOVA | Interaction (sex*genotype) | 0,2394 | 1,417 | F (1, 51) |
| Two-way ANOVA | Sex (main effect) | 0,5624 | 0,34 | F (1, 51) |
| Two-way ANOVA | Genotype (main effect) | 0,5262 | 0,4072 | F (1, 51) |
| Sidak's Post-test | Male nKO vs control | 0,3562 | 1,305 | 51 |
| Sidak's Post-test | Female nKO vs control | 0,9103 | 0,3868 | 51 |

**Panel b**

Old male nKO average energy expenditure during daytime

| Type of test | Comparison | P value | Test statistic | Degrees of Freedom |
|---|---|---|---|---|
| Simple linear regression | Slopes of male nKO and control | 0,7126 | 0,1429 | Dfn=1, Dfd11 |
| Simple linear regression | Intercepts of male nKO and control | 0,0159 | 7,859 | Dfn=1, Dfd12 |

Old female nKO average energy expenditure during daytime

| Type of test | Comparison | P value | Test statistic | Degrees of Freedom |
|---|---|---|---|---|
| Simple linear regression | Slopes of female nKO and control | 0,2152 | 1,712 | Dfn=1, Dfd12 |
| Simple linear regression | Intercepts of female nKO and control | 0,4371 | 0,6429 | Dfn=1, Dfd13 |

**Panel c**

Old nKO male activity

| Type of test | Comparison | P value | Test statistic | Degrees of Freedom |
|---|---|---|---|---|
| Two-way ANOVA | Interaction (phase*genotype) | 0,019 | 6,201 | F (1, 28) |
| Two-way ANOVA | Phase (main effect) | <0,0001 | 33,54 | F (1, 28) |
| Two-way ANOVA | Genotype (main effect) | 0,0038 | 9,957 | F (1, 28) |
| Sidak's Post-test | Male nKO vs control day | 0,8716 | 0,4704 | 28 |
| Sidak's Post-test | Male nKO vs control night | 0,0009 | 3,992 | 28 |

Old nKO female activity

| Type of test | Comparison | P value | Test statistic | Degrees of Freedom |
|---|---|---|---|---|
| Two-way ANOVA | Interaction (phase*genotype) | 0,5589 | 0,35 | F (1, 28) |
| Two-way ANOVA | Phase (main effect) | <0,0001 | 45,51 | F (1, 28) |
| Two-way ANOVA | Genotype (main effect) | 0,5668 | 0,3359 | F (1, 28) |
| Sidak's Post-test | Female nKO vs control day | 0,6573 | 0,8282 | 28 |
| Sidak's Post-test | Female nKO vs control night | >0,9999 | 0,008474 | 28 |

**Panel d**

Old male nKO insulin tolerance test

| Type of test | Comparison | P value | Test statistic | Degrees of Freedom |
|---|---|---|---|---|
| Two-sided t test | AUC male nKO vs control | 0,0037 | 3,19 | 26 |

**Panel e**

Old female nKO insulin tolerance test

| Type of test | Comparison | P value | Test statistic | Degrees of Freedom |
|---|---|---|---|---|
| Two-sided t test | AUC female nKO vs control | 0,2167 | 1,267 | 25 |

**Figure 4: Investigation of mitochondrial function implicated the integrated stress response**

| | | Multiplicity adjusted P value reported for Post-tests | | F(df numerator, df denominator) |
|---|---|---|---|---|

**Panel a**

Old Irs1KO mitochondrial basal respiration

| Type of test | Comparison | P value | Test statistic | Degrees of Freedom |
|---|---|---|---|---|
| Two-sided t test | Male Irs1 KO vs wild type | 0,4457 | 0,8160 | 6 |
| Two-sided t test | Female Irs1 KO vs wild type | 0,4888 | 0,7255 | 8 |

**Panel b**

Old nKO mitochondrial basal respiration

| Type of test | Comparison | P value | Test statistic | Degrees of Freedom |
|---|---|---|---|---|
| Two-sided t test | Male nKO vs control | 0,9443 | 0,0734 | 5 |
| Two-sided t test | Female nKO vs control | 0,6294 | 0,5083 | 6 |

**Panel c**

Old Irs1KO mitochondria spare respiratory capacity

| Type of test | Comparison | P value | Test statistic | Degrees of Freedom |
|---|---|---|---|---|
| Two-sided t test | Male Irs1 KO vs wild type | 0,0016 | 5,4210 | 6 |
| Two-sided t test | Female Irs1 KO vs wild type | 0,4128 | 0,8639 | 8 |

**Panel d**

Old nKO mitochondria spare respiratory capacity

| Type of test | Comparison | P value | Test statistic | Degrees of Freedom |
|---|---|---|---|---|
| Two-sided t test | Male nKO vs control | 0,0153 | 3,6130 | 5 |
| Two-sided t test | Female nKO vs control | 0,9966 | 0,0044 | 6 |

**Panel e**

Old Irs1 KO cortex Atf4 transcripts

| Type of test | Comparison | P value | Test statistic | Degrees of Freedom |
|---|---|---|---|---|
| Two-way ANOVA | Interaction (sex*genotype) | 0,0203 | 5,924 | F (1, 34) |
| Two-way ANOVA | Sex (main effect) | 0,0003 | 16,480 | F (1, 34) |
| Two-way ANOVA | Genotype (main effect) | 0,1272 | 2,4440 | F (1, 34) |
| Sidak's Post-test | Male Irs1 KO vs wild type | 0,0156 | 2,8270 | 34 |
| Sidak's Post-test | Female Irs1 KO vs wild type | 0,7905 | 0,6156 | 34 |

Old Irs1 KO cortex Atf5 transcripts

| Type of test | Comparison | P value | Test statistic | Degrees of Freedom |
|---|---|---|---|---|
| Two-way ANOVA | Interaction (sex*genotype) | 0,0846 | 3,1460 | F (1, 36) |
| Two-way ANOVA | Sex (main effect) | 0,0292 | 5,1620 | F (1, 36) |
| Two-way ANOVA | Genotype (main effect) | 0,0293 | 5,1490 | F (1, 36) |
| Sidak's Post-test | Male Irs1 KO vs wild type | 0,0140 | 2,8590 | 36 |
| Sidak's Post-test | Female Irs1 KO vs wild type | 0,9261 | 0,3503 | 36 |

Old Irs1 KO cortex Chop transcripts

| Type of test | Comparison | P value | Test statistic | Degrees of Freedom |
|---|---|---|---|---|
| Two-way ANOVA | Interaction (sex*genotype) | 0,0445 | 4,326 | F (1, 37) |
| Two-way ANOVA | Sex (main effect) | 0,0460 | 4,264 | F (1, 37) |
| Two-way ANOVA | Genotype (main effect) | 0,0294 | 5,137 | F (1, 37) |
| Sidak's Post-test | Male Irs1 KO vs wild type | 0,0126 | 2,895 | 37 |

| | | | | |
|---|---|---|---|---|
| Sidak's Post-test | Female Irs1 KO vs wild type | 0,9875 | 0,1413 | 37 |

**Panel f**

Old nKO cortex Atf4 transcripts

| Type of test | Comparison | P value | Test statistic | Degrees of Freedom |
|---|---|---|---|---|
| Two-way ANOVA | Interaction (sex*genotype) | 0,0518 | 4,276 | F (1, 20) |
| Two-way ANOVA | Sex (main effect) | 0,4213 | 0,674 | F (1, 20) |
| Two-way ANOVA | Genotype (main effect) | 0,3268 | 1,0100 | F (1, 20) |
| Sidak's Post-test | Male nKO vs control | 0,0797 | 2,1880 | 20 |
| Sidak's Post-test | Female nKO vs control | 0,713 | 0,7462 | 20 |

Old nKO cortex Atf5 transcripts

| Type of test | Comparison | P value | Test statistic | Degrees of Freedom |
|---|---|---|---|---|
| Two-way ANOVA | Interaction (sex*genotype) | 0,0244 | 5,9260 | F (1, 20) |
| Two-way ANOVA | Sex (main effect) | 0,5151 | 0,4391 | F (1, 20) |
| Two-way ANOVA | Genotype (main effect) | 0,0397 | 4,8420 | F (1, 20) |
| Sidak's Post-test | Male nKO vs control | 0,0071 | 3,3010 | 20 |
| Sidak's Post-test | Female nKO vs control | 0,9834 | 0,1642 | 20 |

Old nKO cortex Chop transcripts

| Type of test | Comparison | P value | Test statistic | Degrees of Freedom |
|---|---|---|---|---|
| Two-way ANOVA | Interaction (sex*genotype) | 0,0375 | 4,904 | F (1, 22) |
| Two-way ANOVA | Sex (main effect) | 0,0123 | 7,434 | F (1, 22) |
| Two-way ANOVA | Genotype (main effect) | 0,0454 | 4,498 | F (1, 22) |
| Sidak's Post-test | Male nKO vs control | 0,0127 | 3,014 | 22 |
| Sidak's Post-test | Female nKO vs control | 0,9972 | 0,06735 | 22 |

#### Figure 5: Sex-specific mitochondrial ISR can be activated in non-affected tissues

**Panel a**

Old nKO liver Atf4 transcripts

| Type of test | Comparison | P value (Multiplicity adjusted P value reported for Post-tests) | Test statistic | Degrees of Freedom (F(df numerator, df denominator)) |
|---|---|---|---|---|
| Two-way ANOVA | Interaction (sex*genotype) | 0,2842 | 1,211 | F (1, 20) |
| Two-way ANOVA | Sex (main effect) | 0,3619 | 0,8706 | F (1, 20) |
| Two-way ANOVA | Genotype (main effect) | 0,0722 | 3,603 | F (1, 20) |
| Sidak's Post-test | Male nKO vs control | 0,0885 | 2,135 | 20 |
| Sidak's Post-test | Female nKO vs control | 0,825 | 0,5601 | 20 |

Old nKO liver Atf5 transcripts

| Type of test | Comparison | P value | Test statistic | Degrees of Freedom |
|---|---|---|---|---|
| Two-way ANOVA | Interaction (sex*genotype) | 0,0017 | 13,61 | F (1, 18) |
| Two-way ANOVA | Sex (main effect) | 0,0002 | 20,92 | F (1, 18) |
| Two-way ANOVA | Genotype (main effect) | 0,0164 | 7,008 | F (1, 18) |
| Sidak's Post-test | Male nKO vs control | 0,0005 | 4,56 | 18 |
| Sidak's Post-test | Female nKO vs control | 0,7279 | 0,724 | 18 |

Old nKO liver Chop transcripts

| Type of test | Comparison | P value | Test statistic | Degrees of Freedom |
|---|---|---|---|---|
| Two-way ANOVA | Interaction (sex*genotype) | 0,6412 | 0,2246 | F (1, 18) |
| Two-way ANOVA | Sex (main effect) | 0,6881 | 0,1664 | F (1, 18) |
| Two-way ANOVA | Genotype (main effect) | 0,2631 | 1,334 | F (1, 18) |
| Sidak's Post-test | Male nKO vs control | 0,863 | 0,4902 | 18 |
| Sidak's Post-test | Female nKO vs control | 0,4704 | 1,133 | 18 |

**Panel b**

Old nKO muscle Atf4 transcripts

| Type of test | Comparison | P value | Test statistic | Degrees of Freedom |
|---|---|---|---|---|
| Two-way ANOVA | Interaction (sex*genotype) | 0,0009 | 15,14 | F (1, 20) |
| Two-way ANOVA | Sex (main effect) | 0,0266 | 5,728 | F (1, 20) |
| Two-way ANOVA | Genotype (main effect) | 0,8477 | 0,03788 | F (1, 20) |
| Sidak's Post-test | Male nKO vs control | 0,0173 | 2,909 | 20 |
| Sidak's Post-test | Female nKO vs control | 0,0343 | 2,595 | 20 |

Old nKO muscle Atf5 transcripts

| Type of test | Comparison | P value | Test statistic | Degrees of Freedom |
|---|---|---|---|---|
| Two-way ANOVA | Interaction (sex*genotype) | 0,2282 | 1,545 | F (1, 20) |
| Two-way ANOVA | Sex (main effect) | 0,0898 | 3,178 | F (1, 20) |
| Two-way ANOVA | Genotype (main effect) | 0,4276 | 0,6559 | F (1, 20) |
| Sidak's Post-test | Male nKO vs control | 0,9428 | 0,3085 | 20 |
| Sidak's Post-test | Female nKO vs control | 0,3026 | 1,442 | 20 |

Old nKO muscle Chop transcripts

| Type of test | Comparison | P value | Test statistic | Degrees of Freedom |
|---|---|---|---|---|
| Two-way ANOVA | Interaction (sex*genotype) | 0,0043 | 10,36 | F (1, 20) |
| Two-way ANOVA | Sex (main effect) | 0,7172 | 0,1349 | F (1, 20) |
| Two-way ANOVA | Genotype (main effect) | 0,3795 | 0,8078 | F (1, 20) |
| Sidak's Post-test | Male nKO vs control | 0,2152 | 1,652 | 20 |
| Sidak's Post-test | Female nKO vs control | 0,018 | 2,891 | 20 |

**Panel c**

Old nKO WAT Atf4 transcripts

| Type of test | Comparison | P value | Test statistic | Degrees of Freedom |
|---|---|---|---|---|
| Two-way ANOVA | Interaction (sex*genotype) | 0,4829 | 0,5123 | F (1, 19) |
| Two-way ANOVA | Sex (main effect) | 0,0055 | 9,819 | F (1, 19) |
| Two-way ANOVA | Genotype (main effect) | 0,1335 | 2,458 | F (1, 19) |
| Sidak's Post-test | Male nKO vs control | 0,2049 | 1,685 | 19 |
| Sidak's Post-test | Female nKO vs control | 0,8145 | 0,5792 | 19 |

Old nKO WAT Atf5 transcripts

| Type of test | Comparison | P value | Test statistic | Degrees of Freedom |
|---|---|---|---|---|
| Two-way ANOVA | Interaction (sex*genotype) | 0,0136 | 7,399 | F (1, 19) |
| Two-way ANOVA | Sex (main effect) | 0,0608 | 3,974 | F (1, 19) |
| Two-way ANOVA | Genotype (main effect) | 0,1882 | 1,863 | F (1, 19) |
| Sidak's Post-test | Male nKO vs control | 0,0142 | 3,015 | 19 |
| Sidak's Post-test | Female nKO vs control | 0,6012 | 0,9212 | 19 |

Old nKO WAT Chop transcripts

| Type of test | Comparison | P value | Test statistic | Degrees of Freedom |
|---|---|---|---|---|
| Two-way ANOVA | Interaction (sex*genotype) | 0,0725 | 3,615 | F (1, 19) |
| Two-way ANOVA | Sex (main effect) | 0,1206 | 2,642 | F (1, 19) |
| Two-way ANOVA | Genotype (main effect) | 0,1526 | 2,22 | F (1, 19) |
| Sidak's Post-test | Male nKO vs control | 0,0428 | 2,503 | 19 |
| Sidak's Post-test | Female nKO vs control | 0,9528 | 0,2796 | 19 |

**Panel d**

Old nKO BAT Atf4 transcripts

| Type of test | Comparison | P value | Test statistic | Degrees of Freedom |
|---|---|---|---|---|
| Two-way ANOVA | Interaction (sex*genotype) | 0,486 | 0,5039 | F (1, 20) |
| Two-way ANOVA | Sex (main effect) | 0,6602 | 0,1991 | F (1, 20) |
| Two-way ANOVA | Genotype (main effect) | 0,5033 | 0,4646 | F (1, 20) |
| Sidak's Post-test | Male nKO vs control | 0,5558 | 0,9909 | 20 |
| Sidak's Post-test | Female nKO vs control | 0,9998 | 0,01984 | 20 |

Old nKO BAT Atf5 transcripts

| Type of test | Comparison | P value | Test statistic | Degrees of Freedom |
|---|---|---|---|---|
| Two-way ANOVA | Interaction (sex*genotype) | 0,7277 | 0,1247 | F (1, 20) |
| Two-way ANOVA | Sex (main effect) | 0,7605 | 0,09548 | F (1, 20) |
| Two-way ANOVA | Genotype (main effect) | 0,9763 | 0,0009065 | F (1, 20) |
| Sidak's Post-test | Male nKO vs control | 0,9549 | 0,2729 | 20 |
| Sidak's Post-test | Female nKO vs control | 0,9686 | 0,2268 | 20 |

Old nKO BAT Chop transcripts

| Type of test | Comparison | P value | Test statistic | Degrees of Freedom |
|---|---|---|---|---|
| Two-way ANOVA | Interaction (sex*genotype) | 0,8745 | 0,02558 | F (1, 20) |
| Two-way ANOVA | Sex (main effect) | 0,5862 | 0,3062 | F (1, 20) |
| Two-way ANOVA | Genotype (main effect) | 0,5201 | 0,4287 | F (1, 20) |
| Sidak's Post-test | Male nKO vs control | 0,8136 | 0,5802 | 20 |
| Sidak's Post-test | Female nKO vs control | 0,9281 | 0,3474 | 20 |

**Panel e**

Old nKO gut(ileum) Atf4 transcripts

| Type of test | Comparison | P value | Test statistic | Degrees of Freedom |
|---|---|---|---|---|

| | | | | |
|---|---|---|---|---|
| Two-way ANOVA | Interaction (sex*genotype) | 0,8238 | 0,05104 | F (1, 18) |
| Two-way ANOVA | Sex (main effect) | 0,4608 | 0,5681 | F (1, 18) |
| Two-way ANOVA | Genotype (main effect) | 0,5355 | 0,3991 | F (1, 18) |
| Sidak's Post-test | Male nKO vs control | 0,9487 | 0,2921 | 18 |
| Sidak's Post-test | Female nKO vs control | 0,805 | 0,5963 | 18 |

Old nKO gut(ileum) Atf5 transcripts

| **Type of test** | **Comparison** | **P value** | **Test statistic** | **Degrees of Freedom** |
|---|---|---|---|---|
| Two-way ANOVA | Interaction (sex*genotype) | 0,7719 | 0,08662 | F (1, 18) |
| Two-way ANOVA | Sex (main effect) | 0,0562 | 4,165 | F (1, 18) |
| Two-way ANOVA | Genotype (main effect) | 0,9589 | 0,00273 | F (1, 18) |
| Sidak's Post-test | Male nKO vs control | 0,9814 | 0,1742 | 18 |
| Sidak's Post-test | Female nKO vs control | 0,9648 | 0,2409 | 18 |

Old nKO gut(ileum) Chop transcripts

| **Type of test** | **Comparison** | **P value** | **Test statistic** | **Degrees of Freedom** |
|---|---|---|---|---|
| Two-way ANOVA | Interaction (sex*genotype) | 0,7127 | 0,1399 | F (1, 18) |
| Two-way ANOVA | Sex (main effect) | 0,9085 | 0,01357 | F (1, 18) |
| Two-way ANOVA | Genotype (main effect) | 0,4677 | 0,5505 | F (1, 18) |
| Sidak's Post-test | Male nKO vs control | 0,6778 | 0,8031 | 18 |
| Sidak's Post-test | Female nKO vs control | 0,9604 | 0,2558 | 18 |

#### Supplementary Figure 1: Characterisation of young Irs1KO

| | | Multiplicity adjusted P value reported for Post-tests | | F(df numerator, df denominator) |
|---|---|---|---|---|

**Panel a**

Young Irs1 KO body weight

| Type of test | Comparison | P value | Test statistic | Degrees of Freedom |
|---|---|---|---|---|
| Two-way ANOVA | Interaction (sex*genotype) | <0,0001 | 22,74 | F (1, 76) |
| Two-way ANOVA | Sex (main effect) | <0,0001 | 72,67 | F (1, 76) |
| Two-way ANOVA | Genotype (main effect) | <0,0001 | 332,8 | F (1, 76) |
| Sidak's Post-test | Male Irs1 KO vs wild type | <0,0001 | 16,07 | 76 |
| Sidak's Post-test | Female Irs1 KO vs wild type | <0,0001 | 9,651 | 76 |

Young Irs1 KO body composition

| Type of test | Comparison | P value | Test statistic | Degrees of Freedom |
|---|---|---|---|---|
| Two-way ANOVA | Interaction (sex*genotype) | 0,0041 | 8,758 | F (1, 76) |
| Two-way ANOVA | Sex (main effect) | 0,2301 | 1,464 | F (1, 76) |
| Two-way ANOVA | Genotype (main effect) | 0,0151 | 6,179 | F (1, 76) |
| Sidak's Post-test | Male Irs1 KO vs wild type | 0,0006 | 3,803 | 76 |
| Sidak's Post-test | Female Irs1 KO vs wild type | 0,9299 | 0,3393 | 76 |

**Panel b**

Young Irs1 KO food consumption relative to body weight

| Type of test | Comparison | P value | Test statistic | Degrees of Freedom |
|---|---|---|---|---|
| Two-way ANOVA | Interaction (sex*genotype) | 0,7167 | 0,1335 | F (1, 40) |
| Two-way ANOVA | Sex (main effect) | 0,0007 | 13,44 | F (1, 40) |
| Two-way ANOVA | Genotype (main effect) | 0,0005 | 14,56 | F (1, 40) |
| Sidak's Post-test | Male Irs1 KO vs wild type | 0,0123 | 2,892 | 40 |
| Sidak's Post-test | Female Irs1 KO vs wild type | 0,0331 | 2,498 | 40 |

**Panel c**

Young Irs1 KO average energy expenditure during daytime

| Type of test | Comparison | P value | Test statistic | Degrees of Freedom |
|---|---|---|---|---|
| Two-way ANOVA | Interaction (sex*genotype) | 0,7997 | 0,06525 | F (1, 39) |
| Two-way ANOVA | Sex (main effect) | <0,0001 | 44,27 | F (1, 39) |
| Two-way ANOVA | Genotype (main effect) | <0,0001 | 85,67 | F (1, 39) |
| Sidak's Post-test | Male Irs1 KO vs wild type | <0,0001 | 6,645 | 39 |
| Sidak's Post-test | Female Irs1 KO vs wild type | <0,0001 | 6,443 | 39 |

**Panel d**

Young Irs1 KO male activity

| Type of test | Comparison | P value | Test statistic | Degrees of Freedom |
|---|---|---|---|---|
| Two-way ANOVA | Interaction (phase*genotype) | 0,1718 | 1,939 | F (1, 38) |
| Two-way ANOVA | Phase (main effect) | <0,0001 | 40,97 | F (1, 42) |
| Two-way ANOVA | Genotype (main effect) | 0,1136 | 2,623 | F (1, 42) |
| Sidak's Post-test | Male Irs1 KO vs wild type day | 0,984 | 0,1605 | 38 |
| Sidak's Post-test | Male Irs1 KO vs wild type night | 0,0778 | 2,13 | 38 |

Young Irs1 KO female activity

| Type of test | Comparison | P value | Test statistic | Degrees of Freedom |
|---|---|---|---|---|
| Two-way ANOVA | Interaction (phase*genotype) | 0,4898 | 0,4858 | F (1, 40) |

| | | | | |
|---|---|---|---|---|
| Two-way ANOVA | Phase (main effect) | <0,0001 | 38,66 | F (1, 40) |
| Two-way ANOVA | Genotype (main effect) | 0,6168 | 0,2544 | F (1, 40) |
| Sidak's Post-test | Female Irs1 KO vs wild type day | 0,9884 | 0,1362 | 40 |
| Sidak's Post-test | Female Irs1 KO vs wild type night | 0,6408 | 0,8495 | 40 |

**Panel e**

Young Irs1 KO male insulin tolerance test

| Type of test | Comparison | P value | Test statistic | Degrees of Freedom |
|---|---|---|---|---|
| Two-sided t test | AUC male Irs1 KO vs wild type | <0,0001 | 11,57 | 37 |

**Panel f**

Young Irs1 KO female insulin tolerance test

| Type of test | Comparison | P value | Test statistic | Degrees of Freedom |
|---|---|---|---|---|
| Two-sided t test | AUC female Irs1 KO vs wild type | 0,1672 | 1,407 | 40 |

**Panel g**

Young Irs1 KO male glucose tolerance test

| Type of test | Comparison | P value | Test statistic | Degrees of Freedom |
|---|---|---|---|---|
| Two-sided t test | AUC male Irs1 KO vs wild type | 0,2963 | 1,06 | 35 |

**Panel h**

Young Irs1 KO female glucose tolerance test

| Type of test | Comparison | P value | Test statistic | Degrees of Freedom |
|---|---|---|---|---|
| Two-sided t test | AUC female Irs1 KO vs wild type | 0,0002 | 4,227 | 35 |

#### Supplementary Figure 2: Additional parameters of old Irs1KO

**Panel a**

Multiplicity adjusted P value reported for Post-tests

F(df numerator, df denominator)

Old Irs1 KO food consumption relative to body weight

| Type of test | Comparison | P value | Test statistic | Degrees of Freedom |
|---|---|---|---|---|
| Two-way ANOVA | Interaction (sex*genotype) | 0,614 | 0,2582 | F (1, 42) |
| Two-way ANOVA | Sex (main effect) | 0,7402 | 0,1114 | F (1, 42) |
| Two-way ANOVA | Genotype (main effect) | 0,317 | 1,026 | F (1, 42) |
| Sidak's Post-test | Male Irs1 KO vs wild type | 0,9238 | 0,3555 | 42 |
| Sidak's Post-test | Female Irs1 KO vs wild type | 0,4909 | 1,08 | 42 |

**Panel b**

Old Irs1 KO average energy expenditure during nighttime

| Type of test | Comparison | P value | Test statistic | Degrees of Freedom |
|---|---|---|---|---|
| Two-way ANOVA | Interaction (sex*genotype) | 0,0687 | 3,491 | F (1, 42) |
| Two-way ANOVA | Sex (main effect) | 0,0257 | 5,346 | F (1, 42) |
| Two-way ANOVA | Genotype (main effect) | <0,0001 | 45,18 | F (1, 42) |
| Sidak's Post-test | Male Irs1 KO vs wild type | 0,0028 | 3,418 | 42 |
| Sidak's Post-test | Female Irs1 KO vs wild type | <0,0001 | 6,097 | 42 |

**Panel c**

Old Irs1 KO male glucose tolerance test

| Type of test | Comparison | P value | Test statistic | Degrees of Freedom |
|---|---|---|---|---|
| Two-sided t test | AUC male Irs1 KO vs wild type | 0,0107 | 2,698 | 35 |

**Panel d**

Old Irs1 KO female glucose tolerance test

| Type of test | Comparison | P value | Test statistic | Degrees of Freedom |
|---|---|---|---|---|
| Two-sided t test | AUC female Irs1 KO vs wild type | 0,3473 | 0,9512 | 39 |

#### Supplementary Figure 3: Validation of mouse models used in the study

| | | Multiplicity adjusted P value reported for Post-tests | | F(df numerator, df denominator) |
|---|---|---|---|---|

**Panel a**

Old Irs1 KO cortex Irs1 transcripts

| Type of test | Comparison | P value | Test statistic | Degrees of Freedom |
|---|---|---|---|---|
| Two-way ANOVA | Interaction (sex*genotype) | 0,2607 | 1,34 | F (1, 20) |
| Two-way ANOVA | Sex (main effect) | 0,2607 | 1,34 | F (1, 20) |
| Two-way ANOVA | Genotype (main effect) | <0,0001 | 87,02 | F (1, 20) |
| Sidak's Post-test | Male Irs1 KO vs wild type | <0,0001 | 5,778 | 20 |
| Sidak's Post-test | Female Irs1 KO vs wild type | <0,0001 | 7,415 | 20 |

Old Irs1 KO cortex Irs2 transcripts

| Type of test | Comparison | P value | Test statistic | Degrees of Freedom |
|---|---|---|---|---|
| Two-way ANOVA | Interaction (sex*genotype) | 0,3495 | 0,9256 | F (1, 17) |
| Two-way ANOVA | Sex (main effect) | 0,7437 | 0,1104 | F (1, 17) |
| Two-way ANOVA | Genotype (main effect) | 0,3167 | 1,064 | F (1, 17) |
| Sidak's Post-test | Male Irs1 KO vs wild type | 0,3694 | 1,315 | 17 |
| Sidak's Post-test | Female Irs1 KO vs wild type | 0,9982 | 0,05327 | 17 |

**Panel b**

Old lKO liver Irs1 transcripts

| Type of test | Comparison | P value | Test statistic | Degrees of Freedom |
|---|---|---|---|---|
| Two-way ANOVA | Interaction (sex*genotype) | 0,0058 | 10,34 | F (1, 15) |
| Two-way ANOVA | Sex (main effect) | 0,0054 | 10,58 | F (1, 15) |
| Two-way ANOVA | Genotype (main effect) | <0,0001 | 744,2 | F (1, 15) |
| Sidak's Post-test | Male lKO vs control | <0,0001 | 17,54 | 15 |
| Sidak's Post-test | Female lKO vs control | <0,0001 | 20,96 | 15 |

Old lKO liver Irs2 transcripts

| Type of test | Comparison | P value | Test statistic | Degrees of Freedom |
|---|---|---|---|---|
| Two-way ANOVA | Interaction (sex*genotype) | 0,156 | 2,23 | F (1, 15) |
| Two-way ANOVA | Sex (main effect) | 0,8332 | 0,04592 | F (1, 15) |
| Two-way ANOVA | Genotype (main effect) | 0,0366 | 5,267 | F (1, 15) |
| Sidak's Post-test | Male lKO vs control | 0,0289 | 2,761 | 15 |
| Sidak's Post-test | Female lKO vs control | 0,8318 | 0,5508 | 15 |

**Panel c**

Old mKO muscle Irs1 transcripts

| Type of test | Comparison | P value | Test statistic | Degrees of Freedom |
|---|---|---|---|---|
| Two-way ANOVA | Interaction (sex*genotype) | 0,6313 | 0,2384 | F (1, 18) |
| Two-way ANOVA | Sex (main effect) | 0,6668 | 0,1916 | F (1, 18) |
| Two-way ANOVA | Genotype (main effect) | <0,0001 | 28,92 | F (1, 18) |
| Sidak's Post-test | Male mKO vs control | 0,0056 | 3,457 | 18 |
| Sidak's Post-test | Female mKO vs control | 0,0012 | 4,148 | 18 |

Old mKO muscle Irs2 transcripts

| Type of test | Comparison | P value | Test statistic | Degrees of Freedom |
|---|---|---|---|---|
| Two-way ANOVA | Interaction (sex*genotype) | 0,1645 | 2,082 | F (1, 20) |
| Two-way ANOVA | Sex (main effect) | 0,0433 | 4,656 | F (1, 20) |
| Two-way ANOVA | Genotype (main effect) | 0,5562 | 0,3583 | F (1, 20) |

| | | | | |
|---|---|---|---|---|
| Sidak's Post-test | Male mKO vs control | 0,3016 | 1,444 | 20 |
| Sidak's Post-test | Female mKO vs control | 0,8038 | 0,5972 | 20 |

**Panel d**

Old fKO BAT Irs1 transcripts

| Type of test | Comparison | P value | Test statistic | Degrees of Freedom |
|---|---|---|---|---|
| Two-way ANOVA | Interaction (sex*genotype) | 0,8857 | 0,02118 | F (1, 20) |
| Two-way ANOVA | Sex (main effect) | 0,9378 | 0,006243 | F (1, 20) |
| Two-way ANOVA | Genotype (main effect) | 0,0003 | 19,69 | F (1, 20) |
| Sidak's Post-test | Male fKO vs control | 0,013 | 3,035 | 20 |
| Sidak's Post-test | Female fKO vs control | 0,0082 | 3,24 | 20 |

Old fKO BAT Irs2 transcripts

| Type of test | Comparison | P value | Test statistic | Degrees of Freedom |
|---|---|---|---|---|
| Two-way ANOVA | Interaction (sex*genotype) | 0,6979 | 0,1553 | F (1, 19) |
| Two-way ANOVA | Sex (main effect) | 0,8476 | 0,03797 | F (1, 19) |
| Two-way ANOVA | Genotype (main effect) | 0,4377 | 0,6284 | F (1, 19) |
| Sidak's Post-test | Male fKO vs control | 0,9542 | 0,2753 | 19 |
| Sidak's Post-test | Female fKO vs control | 0,6406 | 0,8599 | 19 |

**Panel e**

Old nKO cortex Irs1 transcripts

| Type of test | Comparison | P value | Test statistic | Degrees of Freedom |
|---|---|---|---|---|
| Two-way ANOVA | Interaction (sex*genotype) | 0,0529 | 4,327 | F (1, 17) |
| Two-way ANOVA | Sex (main effect) | 0,609 | 0,2715 | F (1, 17) |
| Two-way ANOVA | Genotype (main effect) | <0,0001 | 32,54 | F (1, 17) |
| Sidak's Post-test | Male nKO vs control | 0,0409 | 2,551 | 17 |
| Sidak's Post-test | Female nKO vs control | <0,0001 | 5,529 | 17 |

Old nKO cortex Irs2 transcripts

| Type of test | Comparison | P value | Test statistic | Degrees of Freedom |
|---|---|---|---|---|
| Two-way ANOVA | Interaction (sex*genotype) | 0,8129 | 0,05749 | F (1, 20) |
| Two-way ANOVA | Sex (main effect) | 0,6219 | 0,251 | F (1, 20) |
| Two-way ANOVA | Genotype (main effect) | 0,6705 | 0,1865 | F (1, 20) |
| Sidak's Post-test | Male nKO vs control | 0,9885 | 0,1368 | 20 |
| Sidak's Post-test | Female nKO vs control | 0,8721 | 0,4716 | 20 |

**Panel f**

Old gKO ileum Irs1 transcripts

| Type of test | Comparison | P value | Test statistic | Degrees of Freedom |
|---|---|---|---|---|
| Two-way ANOVA | Interaction (sex*genotype) | 0,4878 | 0,5123 | F (1, 12) |
| Two-way ANOVA | Sex (main effect) | 0,1908 | 1,922 | F (1, 12) |
| Two-way ANOVA | Genotype (main effect) | 0,5755 | 0,3313 | F (1, 12) |
| Sidak's Post-test | Male gKO vs control | 0,994 | 0,0991 | 12 |
| Sidak's Post-test | Female gKO vs control | 0,6145 | 0,9132 | 12 |

Old gKO ileum Irs2 transcripts

| Type of test | Comparison | P value | Test statistic | Degrees of Freedom |
|---|---|---|---|---|
| Two-way ANOVA | Interaction (sex*genotype) | 0,5662 | 0,3479 | F (1, 12) |
| Two-way ANOVA | Sex (main effect) | 0,2196 | 1,678 | F (1, 12) |
| Two-way ANOVA | Genotype (main effect) | 0,3412 | 0,9824 | F (1, 12) |
| Sidak's Post-test | Male gKO vs control | 0,9522 | 0,2838 | 12 |

| | | | | |
|---|---|---|---|---|
| Sidak's Post-test | Female gKO vs control | 0,4895 | 1,118 | 12 |

**Panel g**

Old lKO cortex Irs1 transcripts

| Type of test | Comparison | P value | Test statistic | Degrees of Freedom |
|---|---|---|---|---|
| Two-way ANOVA | Interaction (sex*genotype) | 0,7592 | 0,0972 | F (1, 16) |
| Two-way ANOVA | Sex (main effect) | 0,7927 | 0,07141 | F (1, 16) |
| Two-way ANOVA | Genotype (main effect) | 0,9563 | 0,003099 | F (1, 16) |
| Sidak's Post-test | Male lKO vs control | 0,98 | 0,1811 | 16 |
| Sidak's Post-test | Female lKO vs control | 0,9593 | 0,2598 | 16 |

Old mKO cortex Irs1 transcripts

| Type of test | Comparison | P value | Test statistic | Degrees of Freedom |
|---|---|---|---|---|
| Two-way ANOVA | Interaction (sex*genotype) | 0,9874 | 0,000254 | F (1, 20) |
| Two-way ANOVA | Sex (main effect) | 0,4912 | 0,4918 | F (1, 20) |
| Two-way ANOVA | Genotype (main effect) | 0,3426 | 0,9452 | F (1, 20) |
| Sidak's Post-test | Male mKO vs control | 0,7566 | 0,6762 | 20 |
| Sidak's Post-test | Female mKO vs control | 0,7427 | 0,6987 | 20 |

Old fKO cortex Irs1 transcripts

| Type of test | Comparison | P value | Test statistic | Degrees of Freedom |
|---|---|---|---|---|
| Two-way ANOVA | Interaction (sex*genotype) | 0,3282 | 1,004 | F (1, 20) |
| Two-way ANOVA | Sex (main effect) | 0,8837 | 0,02195 | F (1, 20) |
| Two-way ANOVA | Genotype (main effect) | 0,6865 | 0,1677 | F (1, 20) |
| Sidak's Post-test | Male fKO vs control | 0,5512 | 0,9982 | 20 |
| Sidak's Post-test | Female fKO vs control | 0,8974 | 0,419 | 20 |

**Panel h**

Old nKO liver Irs1 transcripts

| Type of test | Comparison | P value | Test statistic | Degrees of Freedom |
|---|---|---|---|---|
| Two-way ANOVA | Interaction (sex*genotype) | 0,3664 | 0,854 | F (1, 20) |
| Two-way ANOVA | Sex (main effect) | 0,0002 | 20,38 | F (1, 20) |
| Two-way ANOVA | Genotype (main effect) | 0,6442 | 0,2199 | F (1, 20) |
| Sidak's Post-test | Male nKO vs control | 0,5551 | 0,992 | 20 |
| Sidak's Post-test | Female nKO vs control | 0,9388 | 0,3197 | 20 |

#### Supplementary Figure 4: Body weight and composition of tissue-specific Irs1KO mice

Multiplicity adjusted P value reported for Post-tests

F(df numerator, df denominator)

**Panel a**

Young lKO body weight

| Type of test | Comparison | P value | Test statistic | Degrees of Freedom |
|---|---|---|---|---|
| Two-way ANOVA | Interaction (sex*genotype) | 0,286 | 1,162 | F (1, 53) |
| Two-way ANOVA | Sex (main effect) | 0,0001 | 307,5 | F (1, 53) |
| Two-way ANOVA | Genotype (main effect) | 0,9321 | 0,007319 | F (1, 53) |
| Sidak's Post-test | Male lKO vs control | 0,7233 | 0,7212 | 53 |
| Sidak's Post-test | Female lKO vs control | 0,671 | 0,8016 | 53 |

**Panel b**

Young lKO body composition

| Type of test | Comparison | P value | Test statistic | Degrees of Freedom |
|---|---|---|---|---|
| Two-way ANOVA | Interaction (sex*genotype) | 0,9112 | 0,01256 | F (1, 53) |
| Two-way ANOVA | Sex (main effect) | 0,0001 | 19,02 | F (1, 53) |
| Two-way ANOVA | Genotype (main effect) | 0,6027 | 0,2743 | F (1, 53) |
| Sidak's Post-test | Male lKO vs control | 0,8746 | 0,462 | 53 |
| Sidak's Post-test | Female lKO vs control | 0,9506 | 0,2836 | 53 |

**Panel c**

Old lKO body weight

| Type of test | Comparison | P value | Test statistic | Degrees of Freedom |
|---|---|---|---|---|
| Two-way ANOVA | Interaction (sex*genotype) | 0,0015 | 11,38 | F (1, 45) |
| Two-way ANOVA | Sex (main effect) | 0,003 | 9,84 | F (1, 45) |
| Two-way ANOVA | Genotype (main effect) | 0,0963 | 2,886 | F (1, 45) |
| Sidak's Post-test | Male lKO vs control | 0,0019 | 3,546 | 45 |
| Sidak's Post-test | Female lKO vs control | 0,4178 | 1,199 | 45 |

**Panel d**

Old lKO body composition

| Type of test | Comparison | P value | Test statistic | Degrees of Freedom |
|---|---|---|---|---|
| Two-way ANOVA | Interaction (sex*genotype) | 0,0284 | 5,132 | F (1, 45) |
| Two-way ANOVA | Sex (main effect) | 0,012 | 6,863 | F (1, 45) |
| Two-way ANOVA | Genotype (main effect) | 0,0157 | 6,305 | F (1, 45) |
| Sidak's Post-test | Male lKO vs control | 0,0034 | 3,339 | 45 |
| Sidak's Post-test | Female lKO vs control | 0,9808 | 0,1758 | 45 |

**Panel e**

Young mKO body weight

| Type of test | Comparison | P value | Test statistic | Degrees of Freedom |
|---|---|---|---|---|
| Two-way ANOVA | Interaction (sex*genotype) | 0,4394 | 0,6065 | F (1, 56) |
| Two-way ANOVA | Sex (main effect) | 0,0001 | 277,7 | F (1, 56) |
| Two-way ANOVA | Genotype (main effect) | 0,3302 | 0,9649 | F (1, 56) |
| Sidak's Post-test | Male mKO vs control | 0,3888 | 1,245 | 56 |
| Sidak's Post-test | Female mKO vs control | 0,987 | 0,1439 | 56 |

**Panel f**

Young mKO body composition

| Type of test | Comparison | P value | Test statistic | Degrees of Freedom |
|---|---|---|---|---|
| Two-way ANOVA | Interaction (sex*genotype) | 0,9278 | 0,00828 | F (1, 56) |
| Two-way ANOVA | Sex (main effect) | 0,0743 | 3,308 | F (1, 56) |
| Two-way ANOVA | Genotype (main effect) | 0,0022 | 10,33 | F (1, 56) |
| Sidak's Post-test | Male mKO vs control | 0,0456 | 2,337 | 56 |
| Sidak's Post-test | Female mKO vs control | 0,0617 | 2,208 | 56 |

**Panel g**

Old mKO body weight

| Type of test | Comparison | P value | Test statistic | Degrees of Freedom |
|---|---|---|---|---|
| Two-way ANOVA | Interaction (sex*genotype) | 0,1221 | 2,478 | F (1, 47) |
| Two-way ANOVA | Sex (main effect) | 0,0001 | 67,16 | F (1, 47) |
| Two-way ANOVA | Genotype (main effect) | 0,45 | 0,5803 | F (1, 47) |
| Sidak's Post-test | Male mKO vs control | 0,1773 | 1,715 | 47 |
| Sidak's Post-test | Female mKO vs control | 0,825 | 0,5548 | 47 |

**Panel h**

Old mKO body composition

| Type of test | Comparison | P value | Test statistic | Degrees of Freedom |
|---|---|---|---|---|
| Two-way ANOVA | Interaction (sex*genotype) | 0,0629 | 3,63 | F (1, 47) |
| Two-way ANOVA | Sex (main effect) | 0,8109 | 0,05789 | F (1, 47) |
| Two-way ANOVA | Genotype (main effect) | 0,0013 | 11,64 | F (1, 47) |
| Sidak's Post-test | Male mKO vs control | 0,0006 | 3,903 | 47 |
| Sidak's Post-test | Female mKO vs control | 0,5225 | 1,029 | 47 |

**Panel i**

Young fKO body weight

| Type of test | Comparison | P value | Test statistic | Degrees of Freedom |
|---|---|---|---|---|
| Two-way ANOVA | Interaction (sex*genotype) | 0,8097 | 0,05855 | F (1, 55) |
| Two-way ANOVA | Sex (main effect) | 0,0001 | 94,13 | F (1, 55) |
| Two-way ANOVA | Genotype (main effect) | 0,4254 | 0,6449 | F (1, 55) |
| Sidak's Post-test | Male fKO vs control | 0,9073 | 0,3933 | 55 |
| Sidak's Post-test | Female fKO vs control | 0,7075 | 0,7455 | 55 |

**Panel j**

Young fKO body composition

| Type of test | Comparison | P value | Test statistic | Degrees of Freedom |
|---|---|---|---|---|
| Two-way ANOVA | Interaction (sex*genotype) | 0,4919 | 0,4786 | F (1, 55) |
| Two-way ANOVA | Sex (main effect) | 0,0033 | 9,466 | F (1, 55) |
| Two-way ANOVA | Genotype (main effect) | 0,0141 | 6,436 | F (1, 55) |
| Sidak's Post-test | Male fKO vs control | 0,0544 | 2,263 | 55 |
| Sidak's Post-test | Female fKO vs control | 0,3497 | 1,316 | 55 |

**Panel k**

Old fKO body weight

| Type of test | Comparison | P value | Test statistic | Degrees of Freedom |
|---|---|---|---|---|
| Two-way ANOVA | Interaction (sex*genotype) | 0,7564 | 0,09735 | F (1, 49) |

| | | | | |
|---|---|---|---|---|
| Two-way ANOVA | Sex (main effect) | 0,0001 | 40,23 | F (1, 49) |
| Two-way ANOVA | Genotype (main effect) | 0,1521 | 2,116 | F (1, 49) |
| Sidak's Post-test | Male fKO vs control | 0,6726 | 0,7996 | 49 |
| Sidak's Post-test | Female fKO vs control | 0,3801 | 1,263 | 49 |

**Panel l**

Old fKO body composition

| Type of test | Comparison | P value | Test statistic | Degrees of Freedom |
|---|---|---|---|---|
| Two-way ANOVA | Interaction (sex*genotype) | 0,7702 | 0,08632 | F (1, 49) |
| Two-way ANOVA | Sex (main effect) | 0,1344 | 2,317 | F (1, 49) |
| Two-way ANOVA | Genotype (main effect) | 0,0008 | 12,86 | F (1, 49) |
| Sidak's Post-test | Male fKO vs control | 0,0504 | 2,304 | 49 |
| Sidak's Post-test | Female fKO vs control | 0,0156 | 2,773 | 49 |

**Panel m**

Young lKO food consumption relative to body weight

| Type of test | Comparison | P value | Test statistic | Degrees of Freedom |
|---|---|---|---|---|
| Two-way ANOVA | Interaction (sex*genotype) | 0,6145 | 0,2602 | F (1, 25) |
| Two-way ANOVA | Sex (main effect) | 0,0033 | 10,53 | F (1, 25) |
| Two-way ANOVA | Genotype (main effect) | 0,1665 | 2,031 | F (1, 25) |
| Sidak's Post-test | Male lKO vs control | 0,3441 | 1,347 | 25 |
| Sidak's Post-test | Female lKO vs control | 0,7664 | 0,6578 | 25 |

**Panel n**

Old lKO food consumption relative to body weight

| Type of test | Comparison | P value | Test statistic | Degrees of Freedom |
|---|---|---|---|---|
| Two-way ANOVA | Interaction (sex*genotype) | 0,1202 | 2,582 | F (1, 26) |
| Two-way ANOVA | Sex (main effect) | 0,0266 | 5,523 | F (1, 26) |
| Two-way ANOVA | Genotype (main effect) | 0,9516 | 0,003755 | F (1, 26) |
| Sidak's Post-test | Male lKO vs control | 0,5479 | 0,9977 | 26 |
| Sidak's Post-test | Female lKO vs control | 0,358 | 1,319 | 26 |

**Panel o**

Young mKO food consumption relative to body weight

| Type of test | Comparison | P value | Test statistic | Degrees of Freedom |
|---|---|---|---|---|
| Two-way ANOVA | Interaction (sex*genotype) | 0,0811 | 3,393 | F (1, 19) |
| Two-way ANOVA | Sex (main effect) | 0,9117 | 0,01264 | F (1, 19) |
| Two-way ANOVA | Genotype (main effect) | 0,7425 | 0,1112 | F (1, 19) |
| Sidak's Post-test | Male mKO vs control | 0,4282 | 1,203 | 19 |
| Sidak's Post-test | Female mKO vs control | 0,3255 | 1,396 | 19 |

**Panel p**

Old mKO food consumption relative to body weight

| Type of test | Comparison | P value | Test statistic | Degrees of Freedom |
|---|---|---|---|---|
| Two-way ANOVA | Interaction (sex*genotype) | 0,23 | 1,52 | F (1, 23) |
| Two-way ANOVA | Sex (main effect) | 0,0011 | 13,8 | F (1, 23) |
| Two-way ANOVA | Genotype (main effect) | 0,6233 | 0,2478 | F (1, 23) |
| Sidak's Post-test | Male mKO vs control | 0,3981 | 1,249 | 23 |
| Sidak's Post-test | Female mKO vs control | 0,8518 | 0,5098 | 23 |

**Panel q**

Young fKO food consumption relative to body weight

| Type of test | Comparison | P value | Test statistic | Degrees of Freedom |
|---|---|---|---|---|
| Two-way ANOVA | Interaction (sex*genotype) | 0,4403 | 0,6228 | F (1, 18) |
| Two-way ANOVA | Sex (main effect) | 0,2234 | 1,59 | F (1, 18) |
| Two-way ANOVA | Genotype (main effect) | 0,4461 | 0,6067 | F (1, 18) |
| Sidak's Post-test | Male fKO vs control | >0,9999 | 0,007534 | 18 |
| Sidak's Post-test | Female fKO vs control | 0,5092 | 1,068 | 18 |

**Panel r**

Old fKO food consumption relative to body weight

| Type of test | Comparison | P value | Test statistic | Degrees of Freedom |
|---|---|---|---|---|
| Two-way ANOVA | Interaction (sex*genotype) | 0,3381 | 0,957 | F (1, 23) |
| Two-way ANOVA | Sex (main effect) | 0,1125 | 2,723 | F (1, 23) |
| Two-way ANOVA | Genotype (main effect) | 0,085 | 3,24 | F (1, 23) |
| Sidak's Post-test | Male fKO vs control | 0,8172 | 0,5726 | 23 |
| Sidak's Post-test | Female fKO vs control | 0,1128 | 1,994 | 23 |

#### Supplementary Figure 5: Energy expenditure of tissue-specific Irs1KO mice

| | | Multiplicity adjusted P value reported for Post-tests | | F(df numerator, df denominator) |
|---|---|---|---|---|

**Panel a**

Young male average energy expenditure during daytime

| Type of test | Comparison | P value | Test statistic | Degrees of Freedom |
|---|---|---|---|---|
| Simple linear regression | Slopes of male lKO and control | 0,7625 | 0,0964 | DFn=1 , DFd=10 |
| Simple linear regression | Intercepts of male lKO and control | 0,4052 | 0,749 | DFn=1 , DFd=11 |

Young male average energy expenditure during nighttime

| Type of test | Comparison | P value | Test statistic | Degrees of Freedom |
|---|---|---|---|---|
| Simple linear regression | Slopes of male lKO and control | 0,4589 | 0,593 | DFn=1 , DFd=10 |
| Simple linear regression | Intercepts of male lKO and control | 0,5176 | 0,447 | DFn=1 , DFd=11 |

**Panel b**

Young female average energy expenditure during daytime

| Type of test | Comparison | P value | Test statistic | Degrees of Freedom |
|---|---|---|---|---|
| Simple linear regression | Slopes of female lKO and control | 0,6147 | 0,267 | DFn=1 , DFd=12 |
| Simple linear regression | Intercepts of female lKO and control | 0,212 | 1,72 | DFn=1 , DFd=13 |

Young female average energy expenditure during nighttime

| Type of test | Comparison | P value | Test statistic | Degrees of Freedom |
|---|---|---|---|---|
| Simple linear regression | Slopes of female lKO and control | 0,4305 | 0,665 | DFn=1 , DFd=12 |
| Simple linear regression | Intercepts of female lKO and control | 0,6411 | 0,228 | DFn=1 , DFd=13 |

**Panel c**

Old male average energy expenditure during daytime

| Type of test | Comparison | P value | Test statistic | Degrees of Freedom |
|---|---|---|---|---|
| Simple linear regression | Slopes of male lKO and control | 0,0923 | 4,32 | DFn=1 , DFd=5 |
| Simple linear regression | Intercepts of male lKO and control | 0,1578 | 2,6 | DFn=1 , DFd=6 |

Old male average energy expenditure during nighttime

| Type of test | Comparison | P value | Test statistic | Degrees of Freedom |
|---|---|---|---|---|
| Simple linear regression | Slopes of male lKO and control | 0,0287 | 9,25 | DFn=1 , DFd=5 |
| Simple linear regression | Intercepts of male lKO and control | | | |

**Panel d**

Old female average energy expenditure during daytime

| Type of test | Comparison | P value | Test statistic | Degrees of Freedom |
|---|---|---|---|---|
| Simple linear regression | Slopes of female lKO and control | 0,0228 | 7,51 | DFn=1 , DFd=9 |
| Simple linear regression | Intercepts of female lKO and control | | | |

Old female average energy expenditure during nighttime

| Type of test | Comparison | P value | Test statistic | Degrees of Freedom |
|---|---|---|---|---|
| Simple linear regression | Slopes of female lKO and control | 0,281 | 1,32 | DFn=1 , DFd=9 |
| Simple linear regression | Intercepts of female lKO and control | 0,6897 | 0,169 | DFn=1 , DFd=10 |

**Panel e**

Young male average energy expenditure during daytime

| Type of test | Comparison | P value | Test statistic | Degrees of Freedom |
|---|---|---|---|---|
| Simple linear regression | Slopes of male mKO and control | 0,7283 | 0,128 | DFn=1 , DFd=10 |
| Simple linear regression | Intercepts of male mKO and control | 0,9452 | 0,00494 | DFn=1 , DFd=11 |

Young male average energy expenditure during nighttime

| Type of test | Comparison | P value | Test statistic | Degrees of Freedom |
|---|---|---|---|---|
| Simple linear regression | Slopes of male mKO and control | 0,7615 | 0,0973 | DFn=1 , DFd=10 |
| Simple linear regression | Intercepts of male mKO and control | 0,8278 | 0,0496 | DFn=1 , DFd=11 |

**Panel f**

Young female average energy expenditure during daytime

| Type of test | Comparison | P value | Test statistic | Degrees of Freedom |
|---|---|---|---|---|
| Simple linear regression | Slopes of female mKO and control | 0,9171 | 0,0118 | DFn=1 , DFd=6 |
| Simple linear regression | Intercepts of female mKO and control | 0,5128 | 0,475 | DFn=1 , DFd=7 |

Young female average energy expenditure during nighttime

| Type of test | Comparison | P value | Test statistic | Degrees of Freedom |
|---|---|---|---|---|
| Simple linear regression | Slopes of female mKO and control | 0,3969 | 0,832 | DFn=1 , DFd=6 |
| Simple linear regression | Intercepts of female mKO and control | 0,9329 | 0,00762 | DFn=1 , DFd=7 |

**Panel g**

Old male average energy expenditure during daytime

| Type of test | Comparison | P value | Test statistic | Degrees of Freedom |
|---|---|---|---|---|
| Simple linear regression | Slopes of male mKO and control | 0,0641 | 4,33 | DFn=1 , DFd=10 |
| Simple linear regression | Intercepts of male mKO and control | 0,8735 | 0,0266 | DFn=1 , DFd=11 |

Old male average energy expenditure during nighttime

| Type of test | Comparison | P value | Test statistic | Degrees of Freedom |
|---|---|---|---|---|
| Simple linear regression | Slopes of male mKO and control | 0,0768 | 3,89 | DFn=1 , DFd=10 |
| Simple linear regression | Intercepts of male mKO and control | 0,9309 | 0,00787 | DFn=1 , DFd=11 |

**Panel h**

Old female average energy expenditure during daytime

| Type of test | Comparison | P value | Test statistic | Degrees of Freedom |
|---|---|---|---|---|
| Simple linear regression | Slopes of female mKO and control | 0,9184 | 0,0111 | DFn=1 , DFd=9 |
| Simple linear regression | Intercepts of female mKO and control | 0,9382 | 0,00632 | DFn=1 , DFd=10 |

Old female average energy expenditure during nighttime

| Type of test | Comparison | P value | Test statistic | Degrees of Freedom |
|---|---|---|---|---|
| Simple linear regression | Slopes of female mKO and control | 0,9764 | 0,000929 | DFn=1 , DFd=9 |
| Simple linear regression | Intercepts of female mKO and control | 0,5742 | 0,337 | DFn=1 , DFd=10 |

**Panel i**

Young male average energy expenditure during daytime

| Type of test | Comparison | P value | Test statistic | Degrees of Freedom |
|---|---|---|---|---|
| Simple linear regression | Slopes of male fKO and control | 0,3128 | 1,16 | DFn=1 , DFd=8 |
| Simple linear regression | Intercepts of male fKO and control | 0,0802 | 3,89 | DFn=1 , DFd=9 |

Young male average energy expenditure during nighttime

| Type of test | Comparison | P value | Test statistic | Degrees of Freedom |
|---|---|---|---|---|
| Simple linear regression | Slopes of male fKO and control | 0,436 | 0,672 | DFn=1 , DFd=8 |
| Simple linear regression | Intercepts of male fKO and control | 0,1431 | 2,57 | DFn=1 , DFd=9 |

**Panel j**

Young female average energy expenditure during daytime

| Type of test | Comparison | P value | Test statistic | Degrees of Freedom |
|---|---|---|---|---|
| Simple linear regression | Slopes of female fKO and control | 0,8741 | 0,0264 | DFn=1 , DFd=10 |

| | | | | |
|---|---|---|---|---|
| Simple linear regression | Intercepts of female fKO and control | 0,9097 | 0,0135 | DFn=1 , DFd=11 |

Young female average energy expenditure during nighttime

| Type of test | Comparison | P value | Test statistic | Degrees of Freedom |
|---|---|---|---|---|
| Simple linear regression | Slopes of female fKO and control | 0,6378 | 0,236 | DFn=1 , DFd=10 |
| Simple linear regression | Intercepts of female fKO and control | 0,962 | 0,00238 | DFn=1 , DFd=11 |

**Panel k**

Old male average energy expenditure during daytime

| Type of test | Comparison | P value | Test statistic | Degrees of Freedom |
|---|---|---|---|---|
| Simple linear regression | Slopes of male fKO and control | 0,0568 | 4,77 | DFn=1 , DFd=9 |
| Simple linear regression | Intercepts of male fKO and control | 0,8595 | 0,033 | DFn=1 , DFd=10 |

Old male average energy expenditure during nighttime

| Type of test | Comparison | P value | Test statistic | Degrees of Freedom |
|---|---|---|---|---|
| Simple linear regression | Slopes of male fKO and control | 0,0658 | 4,38 | DFn=1 , DFd=9 |
| Simple linear regression | Intercepts of male fKO and control | 0,6395 | 0,233 | DFn=1 , DFd=10 |

**Panel l**

Old female average energy expenditure during daytime

| Type of test | Comparison | P value | Test statistic | Degrees of Freedom |
|---|---|---|---|---|
| Simple linear regression | Slopes of female fKO and control | 0,1516 | 2,41 | DFn=1 , DFd=10 |
| Simple linear regression | Intercepts of female fKO and control | 0,9205 | 0,0104 | DFn=1 , DFd=11 |

Old female average energy expenditure during nighttime

| Type of test | Comparison | P value | Test statistic | Degrees of Freedom |
|---|---|---|---|---|
| Simple linear regression | Slopes of female fKO and control | 0,1632 | 2,27 | DFn=1 , DFd=10 |
| Simple linear regression | Intercepts of female fKO and control | 0,9963 | 0,0000225 | DFn=1 , DFd=11 |

#### Supplementary Figure 6: Locomotor activity of tissue-specific Irs1KO mice

| | | Multiplicity adjusted P value reported for Post-tests | | F(df numerator, df denominator) |
|---|---|---|---|---|

**Panel a**

Young male lKO Home-cage activity

| Type of test | Comparison | P value | Test statistic | Degrees of Freedom |
|---|---|---|---|---|
| Two-way ANOVA | Interaction (phase*genotype) | 0,6828 | 0,1711 | F (1, 24) |
| Two-way ANOVA | Phase (main effect) | <0,0001 | 45,61 | F (1, 24) |
| Two-way ANOVA | Genotype (main effect) | 0,7112 | 0,1403 | F (1, 24) |
| Sidak's Post-test | Day lKO vs control | 0,8256 | 0,5574 | 24 |
| Sidak's Post-test | Night lKO vs control | 0,9995 | 0,02759 | 24 |

Young female lKO Home-cage activity

| Type of test | Comparison | P value | Test statistic | Degrees of Freedom |
|---|---|---|---|---|
| Two-way ANOVA | Interaction (phase*genotype) | 0,7549 | 0,09939 | F (1, 28) |
| Two-way ANOVA | Phase (main effect) | <0,0001 | 51,24 | F (1, 28) |
| Two-way ANOVA | Genotype (main effect) | 0,73 | 0,1215 | F (1, 28) |
| Sidak's Post-test | Day lKO vs control | 0,8721 | 0,4694 | 28 |
| Sidak's Post-test | Night lKO vs control | 0,9997 | 0,02354 | 28 |

**Panel b**

Old male lKO Home-cage activity

| Type of test | Comparison | P value | Test statistic | Degrees of Freedom |
|---|---|---|---|---|
| Two-way ANOVA | Interaction (phase*genotype) | 0,8435 | 0,03998 | F (1, 20) |
| Two-way ANOVA | Phase (main effect) | 0,0007 | 16,16 | F (1, 20) |
| Two-way ANOVA | Genotype (main effect) | 0,8138 | 0,05698 | F (1, 20) |
| Sidak's Post-test | Day lKO vs control | 0,9422 | 0,3102 | 20 |
| Sidak's Post-test | Night lKO vs control | 0,9995 | 0,0274 | 20 |

Old female lKO Home-cage activity

| Type of test | Comparison | P value | Test statistic | Degrees of Freedom |
|---|---|---|---|---|
| Two-way ANOVA | Interaction (phase*genotype) | 0,9361 | 0,006529 | F (1, 32) |
| Two-way ANOVA | Phase (main effect) | 0,0001 | 26,86 | F (1, 32) |
| Two-way ANOVA | Genotype (main effect) | 0,906 | 0,01416 | F (1, 32) |
| Sidak's Post-test | Day lKO vs control | 0,9995 | 0,02702 | 32 |
| Sidak's Post-test | Night lKO vs control | 0,9876 | 0,1413 | 32 |

**Panel c**

Young male mKO Home-cage activity

| Type of test | Comparison | P value | Test statistic | Degrees of Freedom |
|---|---|---|---|---|
| Two-way ANOVA | Interaction (phase*genotype) | <0,0001 | 27,44 | F (1, 22) |
| Two-way ANOVA | Phase (main effect) | <0,0001 | 151,6 | F (1, 22) |
| Two-way ANOVA | Genotype (main effect) | <0,0001 | 42,73 | F (1, 22) |
| Sidak's Post-test | Day mKO vs control | 0,6013 | 0,9181 | 22 |
| Sidak's Post-test | Night mKO vs control | <0,0001 | 8,326 | 22 |

Young female mKO Home-cage activity

| Type of test | Comparison | P value | Test statistic | Degrees of Freedom |
|---|---|---|---|---|
| Two-way ANOVA | Interaction (phase*genotype) | 0,6829 | 0,1731 | F (1, 16) |

| | | | | |
|---|---|---|---|---|
| Two-way ANOVA | Phase (main effect) | <0,0001 | 46,31 | F (1, 16) |
| Two-way ANOVA | Genotype (main effect) | 0,7618 | 0,09512 | F (1, 16) |
| Sidak's Post-test | Day mKO vs control | 0,9964 | 0,07613 | 16 |
| Sidak's Post-test | Night mKO vs control | 0,8521 | 0,5123 | 16 |

**Panel d**

Old male mKO Home-cage activity

| Type of test | Comparison | P value | Test statistic | Degrees of Freedom |
|---|---|---|---|---|
| Two-way ANOVA | Interaction (phase*genotype) | 0,4075 | 0,7109 | F (1, 24) |
| Two-way ANOVA | Phase (main effect) | 0,0001 | 26,72 | F (1, 24) |
| Two-way ANOVA | Genotype (main effect) | 0,071 | 3,569 | F (1, 24) |
| Sidak's Post-test | Day mKO vs control | 0,7155 | 0,7397 | 24 |
| Sidak's Post-test | Night mKO vs control | 0,1262 | 1,932 | 24 |

Old female mKO Home-cage activity

| Type of test | Comparison | P value | Test statistic | Degrees of Freedom |
|---|---|---|---|---|
| Two-way ANOVA | Interaction (phase*genotype) | 0,5929 | 0,2944 | F (1, 22) |
| Two-way ANOVA | Phase (main effect) | 0,0092 | 8,143 | F (1, 22) |
| Two-way ANOVA | Genotype (main effect) | 0,2188 | 1,603 | F (1, 22) |
| Sidak's Post-test | Day mKO vs control | 0,8511 | 0,5115 | 22 |
| Sidak's Post-test | Night mKO vs control | 0,3826 | 1,279 | 22 |

**Panel e**

Young male fKO Home-cage activity

| Type of test | Comparison | P value | Test statistic | Degrees of Freedom |
|---|---|---|---|---|
| Two-way ANOVA | Interaction (phase*genotype) | 0,4884 | 0,4984 | F (1, 20) |
| Two-way ANOVA | Phase (main effect) | 0,0003 | 19,6 | F (1, 20) |
| Two-way ANOVA | Genotype (main effect) | 0,9135 | 0,01209 | F (1, 20) |
| Sidak's Post-test | Day fKO vs control | 0,8963 | 0,4214 | 20 |
| Sidak's Post-test | Night fKO vs control | 0,8155 | 0,5769 | 20 |

Young female fKO Home-cage activity

| Type of test | Comparison | P value | Test statistic | Degrees of Freedom |
|---|---|---|---|---|
| Two-way ANOVA | Interaction (phase*genotype) | 0,7711 | 0,08679 | F (1, 22) |
| Two-way ANOVA | Phase (main effect) | <0,0001 | 38,18 | F (1, 22) |
| Two-way ANOVA | Genotype (main effect) | 0,2246 | 1,562 | F (1, 22) |
| Sidak's Post-test | Day fKO vs control | 0,4912 | 1,092 | 22 |
| Sidak's Post-test | Night fKO vs control | 0,7565 | 0,6753 | 22 |

**Panel g**

Old male fKO Home-cage activity

| Type of test | Comparison | P value | Test statistic | Degrees of Freedom |
|---|---|---|---|---|
| Two-way ANOVA | Interaction (phase*genotype) | 0,408 | 0,7094 | F (1, 24) |
| Two-way ANOVA | Phase (main effect) | 0,011 | 7,598 | F (1, 24) |
| Two-way ANOVA | Genotype (main effect) | 0,2655 | 1,3 | F (1, 24) |
| Sidak's Post-test | Day fKO vs control | 0,9728 | 0,2107 | 24 |
| Sidak's Post-test | Night fKO vs control | 0,3174 | 1,402 | 24 |

Old female fKO Home-cage activity

| Type of test | Comparison | P value | Test statistic | Degrees of Freedom |
|---|---|---|---|---|

| | | | | |
|---|---|---|---|---|
| Two-way ANOVA | Interaction (phase*genotype) | 0,7895 | 0,07288 | F (1, 24) |
| Two-way ANOVA | Phase (main effect) | <0,0001 | 24,74 | F (1, 24) |
| Two-way ANOVA | Genotype (main effect) | 0,2984 | 1,13 | F (1, 24) |
| Sidak's Post-test | Day fKO vs control | 0,5844 | 0,9424 | 24 |
| Sidak's Post-test | Night fKO vs control | 0,8238 | 0,5607 | 24 |

#### Supplementary Figure 7: Insulin tolerance test of tissue-specific Irs1KO mice

Multiplicity adjusted P value reported for Post-tests

F(df numerator, df denominator)

**Panel a**

Young lKO male insulin tolerance test

| Type of test | Comparison | P value | Test statistic | Degrees of Freedom |
|---|---|---|---|---|
| Two-sided t test | AUC male lKO vs control | 0,3541 | 0,9423 | 28 |

**Panel b**

Young lKO female insulin tolerance test

| Type of test | Comparison | P value | Test statistic | Difference between medians |
|---|---|---|---|---|
| Mann Whitney test | AUC female lKO vs control | 0,7472 | 97 | 209,5 |

**Panel c**

Old lKO male insulin tolerance test

| Type of test | Comparison | P value | Test statistic | Degrees of Freedom |
|---|---|---|---|---|
| Two-sided t test | AUC male lKO vs control | 0,5051 | 0,6777 | 22 |

**Panel d**

Old lKO female insulin tolerance test

| Type of test | Comparison | P value | Test statistic | Degrees of Freedom |
|---|---|---|---|---|
| Two-sided t test | AUC female lKO vs control | 0,1016 | 1,708 | 22 |

**Panel e**

Young mKO male insulin tolerance test

| Type of test | Comparison | P value | Test statistic | Degrees of Freedom |
|---|---|---|---|---|
| Two-sided t test | AUC male mKO vs control | 0,6305 | 0,4863 | 28 |

**Panel f**

Young mKO female insulin tolerance test

| Type of test | Comparison | P value | Test statistic | Difference between medians |
|---|---|---|---|---|
| Mann Whitney test | AUC female mKO vs control | 0,2671 | 85 | -430 |

**Panel g**

Old mKO male insulin tolerance test

| Type of test | Comparison | P value | Test statistic | Degrees of Freedom |
|---|---|---|---|---|
| Two-sided t test | AUC male mKO vs control | 0,9914 | 0,01086 | 24 |

**Panel h**

Old mKO female insulin tolerance test

| Type of test | Comparison | P value | Test statistic | Degrees of Freedom |
|---|---|---|---|---|
| Two-sided t test | AUC female mKO vs control | 0,5385 | 0,6254 | 21 |

**Panel i**

Young fKO male insulin tolerance test

| Type of test | Comparison | P value | Test statistic | Degrees of Freedom |
|---|---|---|---|---|
| Two-sided t test | AUC male fKO vs control | 0,0347 | 2,229 | 26 |

**Panel j**

Young fKO female insulin tolerance test

| Type of test | Comparison | P value | Test statistic | Degrees of Freedom |
|---|---|---|---|---|
| Two-sided t test | AUC female fKO vs control | 0,1592 | 1,448 | 27 |

**Panel k**

Old fKO male insulin tolerance test

| Type of test | Comparison | P value | Test statistic | Degrees of Freedom |
|---|---|---|---|---|
| Two-sided t test | AUC male fKO vs control | 0,7411 | 0,3344 | 23 |

**Panel l**

Old fKO female insulin tolerance test

| Type of test | Comparison | P value | Test statistic | Difference between medians |
|---|---|---|---|---|
| Mann Whitney test | AUC female fKO vs control | 0,022 | 36 | 644,5 |

**Supplementary Figure 8: Glucose tolerance test of tissue-specific Irs1KO mice**

| | | Multiplicity adjusted P value reported for Post-tests | | F(df numerator, df denominator) |
|---|---|---|---|---|

**Panel a**

Young lKO male glucose tolerance test

| Type of test | Comparison | P value | Test statistic | Degrees of Freedom |
|---|---|---|---|---|
| Two-sided t test | AUC male lKO vs control | 0,0012 | 3,598 | 28 |

**Panel b**

Young lKO female glucose tolerance test

| Type of test | Comparison | P value | Test statistic | Difference between medians |
|---|---|---|---|---|
| Mann Whitney test | AUC female lKO vs control | 0,1555 | 66 | 1103 |

**Panel c**

Old lKO male glucose tolerance test

| Type of test | Comparison | P value | Test statistic | Degrees of Freedom |
|---|---|---|---|---|
| Two-sided t test | AUC male lKO vs control | 0,2167 | 1,267 | 25 |

**Panel d**

Old lKO female glucose tolerance test

| Type of test | Comparison | P value | Test statistic | Difference between medians |
|---|---|---|---|---|
| Mann Whitney test | AUC female lKO vs control | 0,6833 | 81 | 292 |

**Panel e**

Young mKO male glucose tolerance test

| Type of test | Comparison | P value | Test statistic | Degrees of Freedom |
|---|---|---|---|---|
| Two-sided t test | AUC male mKO vs control | 0,0375 | 2,184 | 28 |

**Panel f**

Young mKO female glucose tolerance test

| Type of test | Comparison | P value | Test statistic | Degrees of Freedom |
|---|---|---|---|---|
| Two-sided t test | AUC female mKO vs control | 0,0298 | 2,289 | 28 |

**Panel g**

Old mKO male glucose tolerance test

| Type of test | Comparison | P value | Test statistic | Degrees of Freedom |
|---|---|---|---|---|
| Two-sided t test | AUC male mKO vs control | 0,0126 | 2,679 | 26 |

**Panel h**

Old mKO female glucose tolerance test

| Type of test | Comparison | P value | Test statistic | Degrees of Freedom |
|---|---|---|---|---|
| Two-sided t test | AUC female mKO vs control | 0,4192 | 0,8232 | 22 |

**Panel i**

Young fKO male glucose tolerance test

| Type of test | Comparison | P value | Test statistic | Degrees of Freedom |
|---|---|---|---|---|

| | | | | |
|---|---|---|---|---|
| Two-sided t test | AUC male fKO vs control | 0,3383 | 0,9749 | 27 |

**Panel j**

Young fKO female glucose tolerance test

| Type of test | Comparison | P value | Test statistic | Degrees of Freedom |
|---|---|---|---|---|
| Two-sided t test | AUC female fKO vs control | 0,0127 | 2,669 | 27 |

**Panel k**

Old fKO male glucose tolerance test

| Type of test | Comparison | P value | Test statistic | Degrees of Freedom |
|---|---|---|---|---|
| Two-sided t test | AUC male fKO vs control | 0,1753 | 1,397 | 24 |

**Panel l**

Old fKO female glucose tolerance test

| Type of test | Comparison | P value | Test statistic | Degrees of Freedom |
|---|---|---|---|---|
| Two-sided t test | AUC female fKO vs control | 0,0359 | 2,228 | 23 |

#### Supplementary Figure 9: Characterisation of young nKO

| | | Multiplicity adjusted P value reported for Post-tests | | F(df numerator, df denominator) |
|---|---|---|---|---|

**Panel a**

Young nKO body weight

| Type of test | Comparison | P value | Test statistic | Degrees of Freedom |
|---|---|---|---|---|
| Two-way ANOVA | Interaction (sex*genotype) | 0,2754 | 1,22 | F (1, 44) |
| Two-way ANOVA | Sex (main effect) | <0,0001 | 35,7 | F (1, 44) |
| Two-way ANOVA | Genotype (main effect) | 0,1078 | 2,696 | F (1, 44) |
| Sidak's Post-test | Male nKO vs control | 0,1233 | 1,903 | 44 |
| Sidak's Post-test | Female nKO vs control | 0,9098 | 0,3883 | 44 |

Young nKO body composition

| Type of test | Comparison | P value | Test statistic | Degrees of Freedom |
|---|---|---|---|---|
| Two-way ANOVA | Interaction (sex*genotype) | 0,8834 | 0,02176 | F (1, 44) |
| Two-way ANOVA | Sex (main effect) | 0,0003 | 15,07 | F (1, 44) |
| Two-way ANOVA | Genotype (main effect) | 0,2046 | 1,658 | F (1, 44) |
| Sidak's Post-test | Male nKO vs control | 0,6795 | 0,7898 | 44 |
| Sidak's Post-test | Female nKO vs control | 0,5178 | 1,037 | 44 |

**Panel b**

Young nKO food consumption relative to body weight

| Type of test | Comparison | P value | Test statistic | Degrees of Freedom |
|---|---|---|---|---|
| Two-way ANOVA | Interaction (sex*genotype) | 0,5879 | 0,2979 | F (1, 44) |
| Two-way ANOVA | Sex (main effect) | 0,0006 | 13,84 | F (1, 44) |
| Two-way ANOVA | Genotype (main effect) | 0,1444 | 2,209 | F (1, 44) |
| Sidak's Post-test | Male nKO vs control | 0,3049 | 1,408 | 44 |
| Sidak's Post-test | Female nKO vs control | 0,7505 | 0,6793 | 44 |

**Panel c**

Young male nKO average energy expenditure during nighttime

| Type of test | Comparison | P value | Test statistic | Degrees of Freedom |
|---|---|---|---|---|
| Simple linear regression | Slopes of male nKO and control | 0,5153 | 0,4396 | Dfn=1, Dfd19 |
| Simple linear regression | Intercepts of male nKO and control | 0,1811 | 1,92 | Dfn=1, Dfd20 |

Young female nKO average energy expenditure during nighttime

| Type of test | Comparison | P value | Test statistic | Degrees of Freedom |
|---|---|---|---|---|
| Simple linear regression | Slopes of female nKO and control | 0,602 | 0,2804 | Dfn=1, Dfd21 |
| Simple linear regression | Intercepts of female nKO and control | 0,2013 | 1,735 | Dfn=1, Dfd22 |

**Panel d**

Young nKO male activity

| Type of test | Comparison | P value | Test statistic | Degrees of Freedom |
|---|---|---|---|---|
| Two-way ANOVA | Interaction (phase*genotype) | 0,8071 | 0,06035 | F (1, 42) |
| Two-way ANOVA | Phase (main effect) | <0,0001 | 21,11 | F (1, 42) |
| Two-way ANOVA | Genotype (main effect) | 0,704 | 0,1463 | F (1, 42) |
| Sidak's Post-test | Male nKO vs control | 0,9941 | 0,09673 | 42 |
| Sidak's Post-test | Female nKO vs control | 0,8839 | 0,4442 | 42 |

Young nKO female activity

| Type of test | Comparison | P value | Test statistic | Degrees of Freedom |
|---|---|---|---|---|
| Two-way ANOVA | Interaction (phase*genotype) | 0,184 | 1,819 | F (1, 46) |
| Two-way ANOVA | Phase (main effect) | <0,0001 | 29,76 | F (1, 46) |
| Two-way ANOVA | Genotype (main effect) | 0,0061 | 8,259 | F (1, 46) |
| Sidak's Post-test | Male nKO vs control | 0,4909 | 1,078 | 46 |
| Sidak's Post-test | Female nKO vs control | 0,009 | 2,986 | 46 |

**Panel e**

Young nKO male insulin tolerance test

| Type of test | Comparison | P value | Test statistic | Degrees of Freedom |
|---|---|---|---|---|
| Two-sided t test | AUC male nKO vs control | 0,0295 | 2,293 | 28 |

**Panel f**

Young nKO female insulin tolerance test

| Type of test | Comparison | P value | Test statistic | Degrees of Freedom |
|---|---|---|---|---|
| Two-sided t test | AUC female nKO vs control | 0,0118 | 2,708 | 26 |

**Panel g**

Young nKO male glucose tolerance test

| Type of test | Comparison | P value | Test statistic | Degrees of Freedom |
|---|---|---|---|---|
| Two-sided t test | AUC male nKO vs control | 0,1312 | 1,557 | 27 |

**Panel h**

Young nKO female glucose tolerance test

| Type of test | Comparison | P value | Test statistic | Degrees of Freedom |
|---|---|---|---|---|
| Two-sided t test | AUC female nKO vs control | 0,738 | 0,3383 | 24 |

#### Supplementary Figure 10: Additional parameters of old nKO

**Panel a**

Old nKO food consumption relative to body weight

| Type of test | Comparison | P value (Multiplicity adjusted P value reported for Post-tests) | Test statistic | Degrees of Freedom (F(df numerator, df denominator)) |
|---|---|---|---|---|
| Two-way ANOVA | Interaction (sex*genotype) | 0,9281 | 0,008291 | F (1, 28) |
| Two-way ANOVA | Sex (main effect) | 0,0019 | 11,7 | F (1, 28) |
| Two-way ANOVA | Genotype (main effect) | 0,5717 | 0,3274 | F (1, 28) |
| Sidak's Post-test | Male Irs1 KO vs wild type | 0,8733 | 0,4672 | 28 |
| Sidak's Post-test | Female Irs1 KO vs wild type | 0,9299 | 0,3416 | 28 |

**Panel b**

Old male nKO average energy expenditure during nighttime

| Type of test | Comparison | P value | Test statistic | Degrees of Freedom |
|---|---|---|---|---|
| Simple linear regression | Slopes of male nKO and control | 0,8743 | 0,02621 | Dfn=1, Dfd11 |
| Simple linear regression | Intercepts of male nKO and control | 0,0103 | 9,242 | Dfn=1, Dfd12 |

Old female nKO average energy expenditure during nighttime

| Type of test | Comparison | P value | Test statistic | Degrees of Freedom |
|---|---|---|---|---|
| Simple linear regression | Slopes of female nKO and control | 0,2026 | 1,817 | Dfn=1, Dfd12 |
| Simple linear regression | Intercepts of female nKO and control | 0,6823 | 0,1752 | Dfn=1, Dfd13 |

**Panel c**

Old male nKO glucose tolerance test

| Type of test | Comparison | P value | Test statistic | Difference between medians |
|---|---|---|---|---|
| Mann Whitney test | AUC male nKO vs control | 0,2905 | 74,5 | 4530 |

**Panel d**

Old female nKO glucose tolerance test

| Type of test | Comparison | P value | Test statistic | Difference between medians |
|---|---|---|---|---|
| Mann Whitney test | AUC female nKO vs control | 0,4296 | 74 | 2993 |

#### Supplementary Figure 11: Neuronal Syn1Cre expression does not affect peripheral metabolism

| | | Multiplicity adjusted P value reported for Post-tests | | F(df numerator, df denominator) |
|---|---|---|---|---|

**Panel a**

Old Syn1Cre mice body weight

| Type of test | Comparison | P value | Test statistic | Degrees of Freedom |
|---|---|---|---|---|
| Two-way ANOVA | Interaction (sex*genotype) | 0,8334 | 0,04466 | F (1, 54) |
| Two-way ANOVA | Sex (main effect) | 0,0002 | 16,24 | F (1, 54) |
| Two-way ANOVA | Genotype (main effect) | 0,1826 | 1,823 | F (1, 54) |
| Sidak's Post-test | Male Syn1Cre vs wild type | 0,4685 | 1,112 | 54 |
| Sidak's Post-test | Female Syn1Cre vs wild type | 0,6723 | 0,7995 | 54 |

**Panel b**

Old Syn1Cre mice body composition

| Type of test | Comparison | P value | Test statistic | Degrees of Freedom |
|---|---|---|---|---|
| Two-way ANOVA | Interaction (sex*genotype) | 0,6914 | 0,1592 | F (1, 54) |
| Two-way ANOVA | Sex (main effect) | 0,1653 | 1,978 | F (1, 54) |
| Two-way ANOVA | Genotype (main effect) | 0,1554 | 2,076 | F (1, 54) |
| Sidak's Post-test | Male Syn1Cre vs wild type | 0,7098 | 0,742 | 54 |
| Sidak's Post-test | Female Syn1Cre vs wild type | 0,3632 | 1,292 | 54 |

**Panel c**

Old male average energy expenditure during daytime

| Type of test | Comparison | P value | Test statistic | Degrees of Freedom |
|---|---|---|---|---|
| Simple linear regression | Slopes of male Syn1Cre vs wild type | 0,4451 | 0,632 | Dfn=1, Dfd10 |
| Simple linear regression | Intercepts of male Syn1Cre vs wild type | 0,7055 | 0,1504 | Dfn=1, Dfd11 |

**Panel d**

Old female average energy expenditure during daytime

| Type of test | Comparison | P value | Test statistic | Degrees of Freedom |
|---|---|---|---|---|
| Simple linear regression | Slopes of female Syn1Cre vs wild type | 0,5021 | 0,4815 | Dfn=1, Dfd11 |
| Simple linear regression | Intercepts of female Syn1Cre vs wild type | 0,9796 | 0,0006829 | Dfn=1, Dfd12 |

**Panel e**

Old Syn1Cre male activity

| Type of test | Comparison | P value | Test statistic | Degrees of Freedom |
|---|---|---|---|---|
| Two-way ANOVA | Interaction (phase*genotype) | 0,1842 | 1,87 | F (1, 24) |
| Two-way ANOVA | Phase (main effect) | <0,0001 | 29,16 | F (1, 24) |
| Two-way ANOVA | Genotype (main effect) | 0,064 | 3,77 | F (1, 24) |
| Sidak's Post-test | Male Syn1Cre vs wild type day | 0,9029 | 0,406 | 24 |
| Sidak's Post-test | Male Syn1Cre vs wild type night | 0,0551 | 2,34 | 24 |

**Panel f**

Old Syn1Cre female activity

| Type of test | Comparison | P value | Test statistic | Degrees of Freedom |
|---|---|---|---|---|
| Two-way ANOVA | Interaction (phase*genotype) | 0,4245 | 0,6584 | F (1, 26) |

| | | | | |
|---|---|---|---|---|
| Two-way ANOVA | Phase (main effect) | 0,0019 | 11,94 | F (1, 26) |
| Two-way ANOVA | Genotype (main effect) | 0,6397 | 0,2244 | F (1, 26) |
| Sidak's Post-test | Male Syn1Cre vs wild type day | 0,9651 | 0,2388 | 26 |
| Sidak's Post-test | Male Syn1Cre vs wild type night | 0,6055 | 0,9087 | 26 |

**Panel g**

Old Irs1 KO male insulin tolerance test

| Type of test | Comparison | P value | Test statistic | Degrees of Freedom |
|---|---|---|---|---|
| Two-sided t test | AUC male Syn1Cre vs wild type | 0,9433 | 0,07179 | 26 |

**Panel h**

Old Irs1 KO female insulin tolerance test

| Type of test | Comparison | P value | Test statistic | Degrees of Freedom |
|---|---|---|---|---|
| Two-sided t test | AUC female Syn1Cre vs wild type | 0,1634 | 1,433 | 27 |

**Panel j**

Old Irs1 KO male glucose tolerance test

| Type of test | Comparison | P value | Test statistic | Degrees of Freedom |
|---|---|---|---|---|
| Two-sided t test | AUC male Syn1Cre vs wild type | 0,2176 | 1,264 | 26 |

**Panel k**

Old Irs1 KO female glucose tolerance test

| Type of test | Comparison | P value | Test statistic | Difference between medians |
|---|---|---|---|---|
| Mann Whitney test | AUC female Syn1Cre vs wild type | 0,9454 | 94 | 1721 |

#### Supplementary Figure 12: No activation of ISR in brains of young Irs1KO and nKO mice

| | | Multiplicity adjusted P value reported for Post-tests | | F(df numerator, df denominator) |
|---|---|---|---|---|

**Panel a**

Young mitochondrial basal respiration

| Type of test | Comparison | P value | Test statistic | Degrees of Freedom |
|---|---|---|---|---|
| Two-sided t test | Male Irs1 KO vs wild type | 0,3794 | 1,0280 | 3 |

Young mitochondrial basal respiration

| Type of test | Comparison | P value | Test statistic | Degrees of Freedom |
|---|---|---|---|---|
| Two-sided t test | Female Irs1 KO vs wild type | 0,8231 | 0,2542 | 2 |

**Panel b**

Young Irs1KO mitochondria spare respiratory capacity

| Type of test | Comparison | P value | Test statistic | Degrees of Freedom |
|---|---|---|---|---|
| Two-sided t test | Male Irs1 KO vs wild type | 0,7541 | 0,3279 | 6 |

Young Irs1KO mitochondria spare respiratory capacity

| Type of test | Comparison | P value | Test statistic | Degrees of Freedom |
|---|---|---|---|---|
| Two-sided t test | Female Irs1 KO vs wild type | 0,6635 | 0,4689 | 4 |

**Panel c**

Young Irs1 KO cortex Atf4 transcripts

| Type of test | Comparison | P value | Test statistic | Degrees of Freedom |
|---|---|---|---|---|
| Two-way ANOVA | Interaction (sex*genotype) | 0,8113 | 0,0586 | F (1, 20) |
| Two-way ANOVA | Sex (main effect) | 0,4763 | 0,5269 | F (1, 20) |
| Two-way ANOVA | Genotype (main effect) | 0,3254 | 1,0160 | F (1, 20) |
| Sidak's Post-test | Male Irs1 KO vs wild type | 0,6245 | 0,8840 | 20 |
| Sidak's Post-test | Female Irs1 KO vs wild type | 0,8351 | 0,5418 | 20 |

Young Irs1 KO cortex Atf5 transcripts

| Type of test | Comparison | P value | Test statistic | Degrees of Freedom |
|---|---|---|---|---|
| Two-way ANOVA | Interaction (sex*genotype) | 0,798 | 0,0673 | F (1, 20) |
| Two-way ANOVA | Sex (main effect) | 0,9514 | 0,0038 | F (1, 20) |
| Two-way ANOVA | Genotype (main effect) | 0,0578 | 4,0520 | F (1, 20) |
| Sidak's Post-test | Male Irs1 KO vs wild type | 0,2322 | 1,6070 | 20 |
| Sidak's Post-test | Female Irs1 KO vs wild type | 0,406 | 1,24 | 20 |

Young Irs1 KO cortex Mthfd2 transcripts

| Type of test | Comparison | P value | Test statistic | Degrees of Freedom |
|---|---|---|---|---|
| Two-way ANOVA | Interaction (sex*genotype) | 0,3892 | 0,7747 | F (1, 20) |
| Two-way ANOVA | Sex (main effect) | 0,4123 | 0,701 | F (1, 20) |
| Two-way ANOVA | Genotype (main effect) | 0,3669 | 0,8522 | F (1, 20) |
| Sidak's Post-test | Male Irs1 KO vs wild type | 0,9994 | 0,03036 | 20 |
| Sidak's Post-test | Female Irs1 KO vs wild type | 0,3867 | 1,275 | 20 |

**Panel d**

Young nKO cortex Atf4 transcripts

| Type of test | Comparison | P value | Test statistic | Degrees of Freedom |
|---|---|---|---|---|
| Two-way ANOVA | Interaction (sex*genotype) | 0,8838 | 0,0219 | F (1, 20) |

| | | | | |
|---|---|---|---|---|
| Two-way ANOVA | Sex (main effect) | 0,0065 | 9,2350 | F (1, 20) |
| Two-way ANOVA | Genotype (main effect) | 0,8838 | 0,0219 | F (1, 20) |
| Sidak's Post-test | Male nKO vs control | 0,9736 | 0,2078 | 20 |
| Sidak's Post-test | Female nKO vs control | >0,9999 | 0 | 20 |

Young nKO cortex Atf5 transcripts

| **Type of test** | **Comparison** | **P value** | **Test statistic** | **Degrees of Freedom** |
|---|---|---|---|---|
| Two-way ANOVA | Interaction (sex*genotype) | 0,1476 | 2,2690 | F (1, 20) |
| Two-way ANOVA | Sex (main effect) | 0,0066 | 9,1930 | F (1, 20) |
| Two-way ANOVA | Genotype (main effect) | 0,8897 | 0,0197 | F (1, 20) |
| Sidak's Post-test | Male nKO vs control | 0,5762 | 0,9591 | 20 |
| Sidak's Post-test | Female nKO vs control | 0,4445 | 1,173 | 20 |

Young nKO cortex Chop transcripts

| **Type of test** | **Comparison** | **P value** | **Test statistic** | **Degrees of Freedom** |
|---|---|---|---|---|
| Two-way ANOVA | Interaction (sex*genotype) | 0,4104 | 0,7068 | F (1, 20) |
| Two-way ANOVA | Sex (main effect) | 0,0032 | 11,22 | F (1, 20) |
| Two-way ANOVA | Genotype (main effect) | 0,2783 | 1,242 | F (1, 20) |
| Sidak's Post-test | Male nKO vs control | 0,9774 | 0,1922 | 20 |
| Sidak's Post-test | Female nKO vs control | 0,3261 | 1,392 | 20 |

**Supplementary Figure 13: Syn1Cre expression does not lead to brain ISR activation**

| | | Multiplicity adjusted P value reported for Post-tests | | F(df numerator, df denominator) |
|---|---|---|---|---|

**Panel a**

Old mitochondrial basal respiration

| Type of test | Comparison | P value | Test statistic | Degrees of Freedom |
|---|---|---|---|---|
| Two-sided t test | Male Syn1Cre vs wild type | 0,0926 | 1,9990 | 6 |

**Panel b**

Old mitochondrial basal respiration

| Type of test | Comparison | P value | Test statistic | Degrees of Freedom |
|---|---|---|---|---|
| Two-sided t test | Female Syn1Cre vs wild type | 0,5179 | 0,6868 | 6 |

**Panel c**

Old Syn1Cre cortex Atf4 transcripts

| Type of test | Comparison | P value | Test statistic | Degrees of Freedom |
|---|---|---|---|---|
| Two-way ANOVA | Interaction (sex*genotype) | 0,6274 | 0,2434 | F (1, 19) |
| Two-way ANOVA | Sex (main effect) | 0,0027 | 11,9300 | F (1, 19) |
| Two-way ANOVA | Genotype (main effect) | 0,816 | 0,0557 | F (1, 19) |
| Sidak's Post-test | Male Syn1Cre vs wild type | 0,9806 | 0,1778 | 19 |
| Sidak's Post-test | Female Syn1Cre vs wild type | 0,8427 | 0,5284 | 19 |

Old Syn1Cre cortex Atf5 transcripts

| Type of test | Comparison | P value | Test statistic | Degrees of Freedom |
|---|---|---|---|---|
| Two-way ANOVA | Interaction (sex*genotype) | 0,9168 | 0,0112 | F (1, 19) |
| Two-way ANOVA | Sex (main effect) | 0,0085 | 8,6070 | F (1, 19) |
| Two-way ANOVA | Genotype (main effect) | 0,3169 | 1,0570 | F (1, 19) |
| Sidak's Post-test | Male Syn1Cre vs wild type | 0,6899 | 0,7833 | 19 |
| Sidak's Post-test | Female Syn1Cre vs wild type | 0,7619 | 0,6681 | 19 |

Old Syn1Cre cortex Mthfd2 transcripts

| Type of test | Comparison | P value | Test statistic | Degrees of Freedom |
|---|---|---|---|---|
| Two-way ANOVA | Interaction (sex*genotype) | 0,8115 | 0,05846 | F (1, 19) |
| Two-way ANOVA | Sex (main effect) | 0,0247 | 5,953 | F (1, 19) |
| Two-way ANOVA | Genotype (main effect) | 0,4512 | 0,5919 | F (1, 19) |
| Sidak's Post-test | Male Syn1Cre vs wild type | 0,7433 | 0,6985 | 19 |
| Sidak's Post-test | Female Syn1Cre vs wild type | 0,9139 | 0,3822 | 19 |

#### Supplementary Figure 14: IRS1 deletion in peripheral tissues is insufficient to induce local ISR signature in old mice

| | | Multiplicity adjusted P value reported for Post-tests | | F(df numerator, df denominator) |
|---|---|---|---|---|

**Panel a**

Old lKO liver Atf4 transcripts

| Type of test | Comparison | P value | Test statistic | Degrees of Freedom |
|---|---|---|---|---|
| Two-way ANOVA | Interaction (sex*genotype) | 0,668 | 0,1905 | F (1, 17) |
| Two-way ANOVA | Sex (main effect) | 0,7148 | 0,1381 | F (1, 17) |
| Two-way ANOVA | Genotype (main effect) | 0,2793 | 1,249 | F (1, 17) |
| Sidak's Post-test | Male lKO vs control | 0,477 | 1,124 | 17 |
| Sidak's Post-test | Female lKO vs control | 0,8728 | 0,4714 | 17 |

Old lKO liver Atf5 transcripts

| Type of test | Comparison | P value | Test statistic | Degrees of Freedom |
|---|---|---|---|---|
| Two-way ANOVA | Interaction (sex*genotype) | 0,5444 | 0,3847 | F (1, 15) |
| Two-way ANOVA | Sex (main effect) | 0,732 | 0,1217 | F (1, 15) |
| Two-way ANOVA | Genotype (main effect) | 0,0385 | 5,143 | F (1, 15) |
| Sidak's Post-test | Male lKO vs control | 0,1023 | 2,105 | 15 |
| Sidak's Post-test | Female lKO vs control | 0,4748 | 1,132 | 15 |

Old lKO liver Chop transcripts

| Type of test | Comparison | P value | Test statistic | Degrees of Freedom |
|---|---|---|---|---|
| Two-way ANOVA | Interaction (sex*genotype) | 0,2129 | 1,693 | F (1, 15) |
| Two-way ANOVA | Sex (main effect) | 0,1017 | 3,04 | F (1, 15) |
| Two-way ANOVA | Genotype (main effect) | 0,7844 | 0,07757 | F (1, 15) |
| Sidak's Post-test | Male lKO vs control | 0,7166 | 0,7453 | 15 |
| Sidak's Post-test | Female lKO vs control | 0,5028 | 1,085 | 15 |

**Panel b**

Old mKO liver Atf4 transcripts

| Type of test | Comparison | P value | Test statistic | Degrees of Freedom |
|---|---|---|---|---|
| Two-way ANOVA | Interaction (sex*genotype) | 0,1464 | 2,284 | F (1, 20) |
| Two-way ANOVA | Sex (main effect) | 0,0046 | 10,18 | F (1, 20) |
| Two-way ANOVA | Genotype (main effect) | 0,1334 | 2,447 | F (1, 20) |
| Sidak's Post-test | Male mKO vs control | 0,9991 | 0,0375 | 20 |
| Sidak's Post-test | Female mKO vs control | 0,0819 | 2,175 | 20 |

Old mKO liver Atf5 transcripts

| Type of test | Comparison | P value | Test statistic | Degrees of Freedom |
|---|---|---|---|---|
| Two-way ANOVA | Interaction (sex*genotype) | 0,0152 | 7,122 | F (1, 19) |
| Two-way ANOVA | Sex (main effect) | 0,3054 | 1,11 | F (1, 19) |
| Two-way ANOVA | Genotype (main effect) | 0,033 | 5,283 | F (1, 19) |
| Sidak's Post-test | Male mKO vs control | 0,9565 | 0,2682 | 19 |
| Sidak's Post-test | Female mKO vs control | 0,0056 | 3,432 | 19 |

Old mKO liver Chop transcripts

| Type of test | Comparison | P value | Test statistic | Degrees of Freedom |
|---|---|---|---|---|
| Two-way ANOVA | Interaction (sex*genotype) | 0,9197 | 0,01042 | F (1, 20) |
| Two-way ANOVA | Sex (main effect) | 0,0004 | 18,04 | F (1, 20) |
| Two-way ANOVA | Genotype (main effect) | <0,0001 | 25,23 | F (1, 20) |

| | | | | |
|---|---|---|---|---|
| Sidak's Post-test | Male mKO vs control | 0,0034 | 3,624 | 20 |
| Sidak's Post-test | Female mKO vs control | 0,0047 | 3,479 | 20 |

**Panel c**

Old fKO WAT Atf4 transcripts

| Type of test | Comparison | P value | Test statistic | Degrees of Freedom |
|---|---|---|---|---|
| Two-way ANOVA | Interaction (sex*genotype) | 0,9595 | 0,002647 | F (1, 19) |
| Two-way ANOVA | Sex (main effect) | 0,3623 | 0,8714 | F (1, 19) |
| Two-way ANOVA | Genotype (main effect) | 0,4622 | 0,5631 | F (1, 19) |
| Sidak's Post-test | Male mKO vs control | 0,8134 | 0,581 | 19 |
| Sidak's Post-test | Female mKO vs control | 0,8666 | 0,4829 | 19 |

Old fKO WAT Atf5 transcripts

| Type of test | Comparison | P value | Test statistic | Degrees of Freedom |
|---|---|---|---|---|
| Two-way ANOVA | Interaction (sex*genotype) | 0,5293 | 0,4106 | F (1, 19) |
| Two-way ANOVA | Sex (main effect) | 0,6383 | 0,2282 | F (1, 19) |
| Two-way ANOVA | Genotype (main effect) | 0,8582 | 0,03278 | F (1, 19) |
| Sidak's Post-test | Male mKO vs control | 0,8051 | 0,5955 | 19 |
| Sidak's Post-test | Female mKO vs control | 0,9396 | 0,3176 | 19 |

Old fKO WAT Chop transcripts

| Type of test | Comparison | P value | Test statistic | Degrees of Freedom |
|---|---|---|---|---|
| Two-way ANOVA | Interaction (sex*genotype) | 0,7827 | 0,07823 | F (1, 19) |
| Two-way ANOVA | Sex (main effect) | 0,4881 | 0,4999 | F (1, 19) |
| Two-way ANOVA | Genotype (main effect) | 0,3538 | 0,9033 | F (1, 19) |
| Sidak's Post-test | Male mKO vs control | 0,6204 | 0,8913 | 19 |
| Sidak's Post-test | Female mKO vs control | 0,8763 | 0,4634 | 19 |

#### Supplementary Figure 15: IRS1 deletion in peripheral tissues does not induce ISR signature in brains of old mice

**Panel a**

Old lKO cortex Atf4 transcripts

| Type of test | Comparison | P value (Multiplicity adjusted P value reported for Post-tests) | Test statistic | Degrees of Freedom (F(df numerator, df denominator)) |
|---|---|---|---|---|
| Two-way ANOVA | Interaction (sex*genotype) | 0,5066 | 0,4615 | F (1, 16) |
| Two-way ANOVA | Sex (main effect) | 0,1774 | 1,991 | F (1, 16) |
| Two-way ANOVA | Genotype (main effect) | 0,959 | 0,002731 | F (1, 16) |
| Sidak's Post-test | Male nKO vs control | 0,8495 | 0,5173 | 16 |
| Sidak's Post-test | Female nKO vs control | 0,8867 | 0,4434 | 16 |

Old lKO cortex Atf5 transcripts

| Type of test | Comparison | P value | Test statistic | Degrees of Freedom |
|---|---|---|---|---|
| Two-way ANOVA | Interaction (sex*genotype) | 0,3199 | 1,054 | F (1, 16) |
| Two-way ANOVA | Sex (main effect) | 0,0672 | 3,855 | F (1, 16) |
| Two-way ANOVA | Genotype (main effect) | 0,285 | 1,224 | F (1, 16) |
| Sidak's Post-test | Male nKO vs control | 0,2792 | 1,508 | 16 |
| Sidak's Post-test | Female nKO vs control | 0,998 | 0,05638 | 16 |

Old lKO cortex Chop transcripts

| Type of test | Comparison | P value | Test statistic | Degrees of Freedom |
|---|---|---|---|---|
| Two-way ANOVA | Interaction (sex*genotype) | 0,3166 | 1,069 | F (1, 16) |
| Two-way ANOVA | Sex (main effect) | 0,175 | 2,014 | F (1, 16) |
| Two-way ANOVA | Genotype (main effect) | 0,9039 | 0,01504 | F (1, 16) |
| Sidak's Post-test | Male nKO vs control | 0,67 | 0,8177 | 16 |
| Sidak's Post-test | Female nKO vs control | 0,7777 | 0,6442 | 16 |

**Panel b**

Old mKO cortex Atf4 transcripts

| Type of test | Comparison | P value | Test statistic | Degrees of Freedom |
|---|---|---|---|---|
| Two-way ANOVA | Interaction (sex*genotype) | 0,561 | 0,3496 | F (1, 20) |
| Two-way ANOVA | Sex (main effect) | <0,0001 | 48,16 | F (1, 20) |
| Two-way ANOVA | Genotype (main effect) | 0,5209 | 0,4271 | F (1, 20) |
| Sidak's Post-test | Male nKO vs control | 0,6269 | 0,8802 | 20 |
| Sidak's Post-test | Female nKO vs control | 0,9988 | 0,04401 | 20 |

Old mKO cortex Atf5 transcripts

| Type of test | Comparison | P value | Test statistic | Degrees of Freedom |
|---|---|---|---|---|
| Two-way ANOVA | Interaction (sex*genotype) | 0,8228 | 0,0515 | F (1, 20) |
| Two-way ANOVA | Sex (main effect) | 0,5187 | 0,4315 | F (1, 20) |
| Two-way ANOVA | Genotype (main effect) | 0,5649 | 0,3425 | F (1, 20) |
| Sidak's Post-test | Male nKO vs control | 0,817 | 0,5743 | 20 |
| Sidak's Post-test | Female nKO vs control | 0,961 | 0,2534 | 20 |

Old mKO cortex Chop transcripts

| Type of test | Comparison | P value | Test statistic | Degrees of Freedom |
|---|---|---|---|---|
| Two-way ANOVA | Interaction (sex*genotype) | 0,7992 | 0,06646 | F (1, 20) |
| Two-way ANOVA | Sex (main effect) | <0,0001 | 23,57 | F (1, 20) |
| Two-way ANOVA | Genotype (main effect) | 0,9324 | 0,007384 | F (1, 20) |
| Sidak's Post-test | Male nKO vs control | 0,9641 | 0,243 | 20 |
| Sidak's Post-test | Female nKO vs control | 0,9909 | 0,1215 | 20 |

**Panel c**

Old fKO cortex Atf4 transcripts

| Type of test | Comparison | P value | Test statistic | Degrees of Freedom |
|---|---|---|---|---|
| Two-way ANOVA | Interaction (sex*genotype) | 0,4216 | 0,6731 | F (1, 20) |
| Two-way ANOVA | Sex (main effect) | 0,4083 | 0,7133 | F (1, 20) |
| Two-way ANOVA | Genotype (main effect) | 0,9241 | 0,009317 | F (1, 20) |
| Sidak's Post-test | Male nKO vs control | 0,7735 | 0,6484 | 20 |
| Sidak's Post-test | Female nKO vs control | 0,8513 | 0,5119 | 20 |

Old fKO cortex Atf5 transcripts

| Type of test | Comparison | P value | Test statistic | Degrees of Freedom |
|---|---|---|---|---|
| Two-way ANOVA | Interaction (sex*genotype) | 0,2976 | 1,144 | F (1, 20) |
| Two-way ANOVA | Sex (main effect) | 0,6852 | 0,1692 | F (1, 20) |
| Two-way ANOVA | Genotype (main effect) | 0,1941 | 1,806 | F (1, 20) |
| Sidak's Post-test | Male nKO vs control | 0,1961 | 1,706 | 20 |
| Sidak's Post-test | Female nKO vs control | 0,977 | 0,1939 | 20 |

Old fKO cortex Chop transcripts

| Type of test | Comparison | P value | Test statistic | Degrees of Freedom |
|---|---|---|---|---|
| Two-way ANOVA | Interaction (sex*genotype) | 0,3866 | 0,7834 | F (1, 20) |
| Two-way ANOVA | Sex (main effect) | 0,3976 | 0,7474 | F (1, 20) |
| Two-way ANOVA | Genotype (main effect) | 0,6555 | 0,2051 | F (1, 20) |
| Sidak's Post-test | Male nKO vs control | 0,9438 | 0,3056 | 20 |
| Sidak's Post-test | Female nKO vs control | 0,5845 | 0,9461 | 20 |

### Supplementary Table 3

**Genotyping**

| Primers for | Forward sequecne | Reverse Sequence | | |
|---|---|---|---|---|
| PCR LoxP floxed Irs1 | 5´- TCC TCC CAA CCC CTA ATG C -3´ | | Band at 423 | Floxed Irs1 |
| | 5´- GTG TCA CCT CCT CTC GAA AGC -3´ | | Band at 345 | Germline deleted (whole body KO) |
| | 5´- AGC TCG GGT CTA AGT CAC TGC -3´ | | Band at 266 | Wild type |
| PCR Irs1 wild type | 5´-GCC AGG CAC CAG CAT CTT CG-3´ | 5´-TGG CCG CTC CCG AAT TCA AT-3´ | Band at 360 bp | |
| PCR Irs1 knockout | 5´-GCT ACC CGT GAT ATT GCT GAA GAG-3´ | 5´-TGG CCG CTC CCG AAT TCA AT-3´ | Band at 680 bp | |
| PCR - Cre | 5´- CCC AAG AAG AAG AGG AAG GTG TCC -3 | 5´- CCC AGA AAT GCC AGA TTA CG-3´ | Band at 506 bp | Fat and Neuron specific Cre lines |
| PCR - Cre | 5´- CAC GAC CAA GTG ACA GCA AT -3´ | 5´- AGA GAC GGA AAT CCA TCG CT -3´ | Band at 371 bp | Liver, Muscle and gut specific Cre lines |

**Q-RT-PCR**

| Name | Type | Sequence |
|---|---|---|
| ATF4_For | Sybr | CAAGGAGGATGCCTTTTC |
| ATF4_Rev | Sybr | GTCATCCATTCGAAACAGAG |
| ATF5_For | Sybr | AATTTTTAAGGGAAGAGCGG |
| ATF5_Rev | Sybr | TGGAGGACCTCATCTTATTC |
| B2M_For | Sybr | TTCTGGTGCTTGTCTCACTGA |
| B2M_Rev | Sybr | CAGTATGTTCGGCTTCCCATTC |
| CHOP_For | Sybr | CGGAACCTGAGGAGAGAGTG |
| CHOP_Rev | Sybr | CGTTTCCTGGGGATGAGATA |

| Name | Type | Reference (Thermofisher) |
|---|---|---|
| Irs1 | Taqman | Mm01278327_m1 |
| Irs2 | Taqman | Mm03038438_m1 |
| B2M | Taqman | Mm00437762_m1 |
